# Improving GWAS performance in underrepresented groups by appropriate modeling of genetics, environment, and sociocultural factors

**DOI:** 10.1101/2024.10.28.620716

**Authors:** Chelsea C. Cataldo-Ramirez, Meng Lin, Aislinn Mcmahon, Christopher R. Gignoux, Timothy D. Weaver, Brenna M. Henn

## Abstract

Genome-wide association studies (GWAS) and polygenic score (PGS) development are typically constrained by the data available in biobank repositories in which European cohorts are vastly overrepresented. Here, we increase the utility of non-European participant data within the UK Biobank (UKB) by characterizing the genetic affinities of UKB participants who self-identify as Bangladeshi, Indian, Pakistani, “White and Asian” (WA), and “Any Other Asian” (AOA), towards creating a more robust South Asian sample size for future genetic analyses. We assess the relationships between genetic structure and self-selected ethnic identities and use consistent patterns of clustering in the dataset to train a support vector machine (SVM). The SVM was utilized to reassign *n* = 1,853 AOA and WA participants at the subcontinental level, and increase the sample size of the UKB South Asian group by 1,381 additional participants. We further leverage these samples to assess GWAS performance and PGS development. We include environmental covariates in the height GWAS by implementing a rigorous covariate selection procedure, and compare the outputs of two GWAS models: GWAS_null_ and GWAS_env_. We show that PGS performance derived from both GWAS models yield comparable prediction to PGS models developed with an order of magnitude larger training, and environmentally-adjusted PGS models reduce the sex-bias in predictive performance. In summary, we demonstrate how GWAS performance can be improved by leveraging ambiguous ethnicity codes, ancestry matched imputation panels, and including environmental covariates.

## Background

Unequal representation of diverse populations within biobanks has resulted in comparatively few insights into the genetic architecture of complex traits for many non-European-descent populations [1]. This Eurocentric emphasis within genetic research is often attributed to the current state of data availability, in which European-descent genomic data is most abundant. Consequently, the prevailing assumption is that smaller datasets will fail to produce informative results. This perspective acts to disincentivize prospective researchers from attempting to make the best use of the datasets currently available [2]. However, utilizing smaller samples need not be a futile undertaking. Genotype-phenotype research design can be optimized to increase the utility of smaller datasets. Here, we highlight how a detailed consideration of the intersection of genetic ancestry affinities and ethnic identifiers, in conjunction with incorporating environmental adjustments into GWAS of height can improve the quality of results output by smaller, often excluded non-European-descent samples.

PGS performance declines when applied to groups outside of the discovery GWAS training sample [3,4,5], and because European cohorts are vastly overrepresented within the available data, PGS applicability is much reduced in non-Europeans [6, 7]. Specifically, variation in frequencies and population-specificity of causal alleles, in conjunction with differences in linkage disequilibrium (LD) between populations, contribute to the limitations of out-of-sample PGS portability [8, 9]. Fine-mapping with exome data, improved imputation for GWAS, and including LD-informed analyses can ameliorate the effects of these factors [4, 10]. Methods that integrate discovery-sample GWAS summary statistics with LD structure from multi-ethnic cohorts to improve the predictive utility of European-derived PGS have been a primary focus of PGS research recently [12,13,14]. However, improving GWAS discovery by increasing the sample sizes of minority cohorts or including relevant environmental variables have not been explored as systematically as other approaches.

Using UK Biobank (UKB) South Asian participants as a case study, we aim to test how the discovery of significant genetic variants and accuracy of PGS prediction of complex traits may be improved by: 1) considering several levels of participant identity and genetic characteristics when establishing a sample dataset; and 2) adjusting for environmental confounders in the GWAS model. We focus on height as a canonical complex trait. The UKB is a vast resource for public health, epidemiology, and precision medicine research, containing genetic, phenotypic, and environmental data for over 500,000 adult UK residents [14]. While the majority of participants self-identify as “white” (defined by Data Field 21000 as including British, Irish, and “Any other white background”; *n* > 500,000 as of January 2024), there are over 28,000 individuals who do not identify as such. The Pan-UK Biobank and additional studies have identified tens of thousands of individuals with primarily non-European genetically inferred ancestry [3, 16]. The largest geographically-defined grouping of non-“white”-identifying participants is a conglomerate of South Asian identities, which include Bangladeshi, Indian, and Pakistani identifying individuals (Data Field 21000). However, ethnic identity does not necessarily reflect high genetic affinities with other individuals or populations occupying the same geographical region at the same point in time [17,18,19,20].

The use of ethnicity as a proxy, or at least as a starting point, for identifying a genetically homogenous population is further complicated when the ethnic labels participants are able to choose from are limited, discrete/categorical, follow several naming conventions, and/or are un-defined, as is the case with UKB participants who identify as “White and Asian” (WA) or “Any other Asian” (AOA). This, in conjunction with the limited UKB Data Fields associated with genetic affinity available to researchers and historical apprehension to including genetically “admixed” populations in GWAS research has resulted in reduced inclusion for AOA and WA participants [5, 21, 22]. To rectify this, we characterize in detail the genetic affinities of UKB AOA and WA participants, and provide clarification on their genetic similarities to 1000 Genomes (1KG) reference populations (Table S1.6) and UKB Bangladeshi, Indian, and Pakistani (together referred to as UKB-BIP) participants (Table 1). This, accordingly, contributes to our first aim of increasing participant inclusion by integrating several descriptors of participant identity when building an associated genetic dataset.

**Table 1.** UK Biobank sample.

| **Ethnicity\*** | **Group Acronym\*\*** | ***n*** |
| --- | --- | --- |
| Bangladeshi | BIP | 212 |
| Indian | BIP | 5016 |
| Pakistani | BIP | 1528 |
| Any other Asian | AOA | 1614 |
| White and Asian | WA | 549 |

\*The “Ethnicity” column reflects the participants’ response to UKB data field 21000 (“Ethnic background”).
\*\*The “Group Acronym” column reflects the original grouping of participants used in this study.

Previous investigations into the genetic ancestries of UKB participants have aimed to broadly categorize participants at the subcontinental level, but in-depth assessments of population affinities within the identified non-European groupings have been lacking. Privé et al. (2022) defined nine subcontinental ancestry groups within the UKB cohort and demonstrated that PGS prediction significantly decreases when PGS models trained on Northwestern European individuals are applied to the non-European ancestry groups [3]. However, non-European ancestry groups were, in part, restricted to participants whose self-reported ethnicity (as assessed by a within-UKB PCA) matched their country of origin, resulting in PGS exclusion for many participants as individuals within diaspora communities were not retained (see Privé et al., 2022, Fig. 1 and Table SA1).

**Figure 1.**
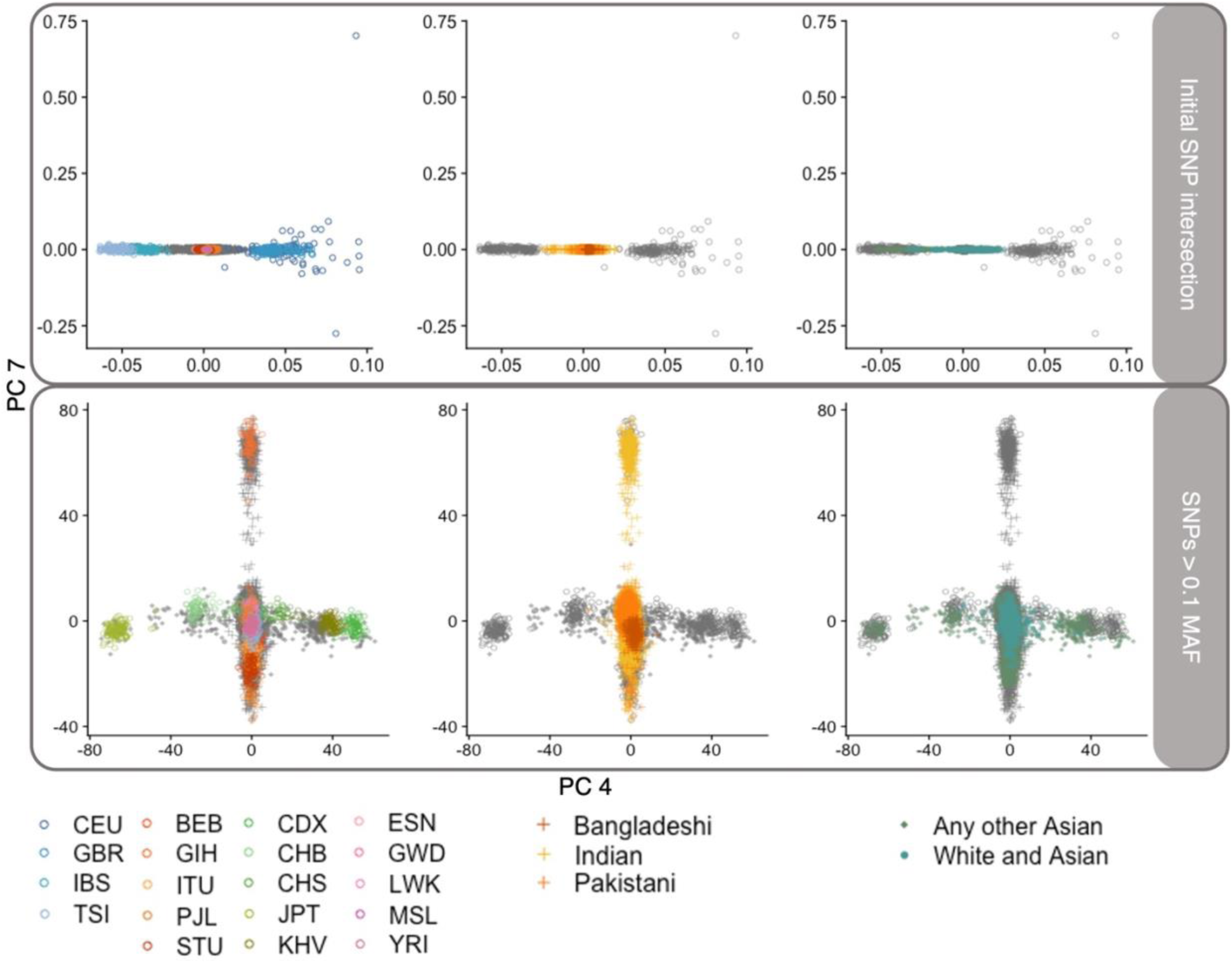
UKB BIP (middle column) and AOA and WA (right column) array data projected onto 1KG PC4 and PC7 space, when using standard QC thresholds for MAF (top row) and stringent MAF thresholds (bottom row). For each plot, 1KG data are represented by open circles, BIP are represented by plus signs, AOA are represented by filled diamonds, and WA are represented by filled circles. All data are plotted in the left column, with UKB data grayed. BIP and 1KG data are plotted in the middle column, with 1KG data grayed. AOA, WA, and 1KG data are plotted in the right column, with 1KG data grayed. Top: When using the full SNP intersection between UKB and 1KG data, PC4 and above are both driven by European genetic variation (See SI Section 1.2.3 for more information). Bottom: when more stringent MAF thresholds are used to filter out all but common SNPs (i.e., variants with MAF > 0.1 retained), PC4 is driven by East Asian genetic variation and PC7 is driven by South Asian genetic variation. Standard QC thresholds for filtering UKB array data will emphasize structure within European axes of variation and may only be appropriate for addressing within-sample structure if just the first three PCs are being considered. However, this may bias the interpretation of relative genetic variance within the sample data as well as patterns of global population structure (therefore also biasing attempts to characterize genetic ancestry).

Constantinescu et al. aimed to characterize UKB participants of “non-white British ancestry” for the explicit purpose of identifying relatively homogenous groups (2022: 2), to facilitate the inclusion of these participants in future health research [16]. We implement a similar approach to Constantinescu et al. to characterize a subset of the UKB participants of “non-white British ancestry” at a subcontinental resolution as well as at the level of 1KG reference groups [16]. Our subcontinental approach, however, differs at three primary levels from the approach taken by Constantinescu et al.: 1) we focus our analyses on a UKB South Asian metapopulation as opposed to their inclusion of all “non-white” individuals, therefore providing more detail into South Asian population substructure; 2) we utilize a subset of the 1KG-UKB genetic data intersection in our genetic affinity analyses chosen specifically to reduce the effects of ascertainment bias resulting from the use of the UKB SNP array; and 3) while our subcontinental classifications are also PCA based, we choose to utilize support vector machines (SVM) instead of K-means clustering to infer population assignments.

After creating a broader UKB South Asian dataset, our second aim is to assess how adjusting for environmental confounders in the GWAS model affects GWAS and PGS results. We implement an environmental variable selection procedure to identify a subset of covariates to control for confounding in our GWAS model (referred to here as GWAS_env_). We then assess the downstream effects of GWAS_env_ by developing a height PGS model and compare its performance to published population-matched PGS.

## RESULTS

### Genetic affinities of UKB South Asian participants

The genetic affinities of UKB participants who self-identify as Bangladeshi, Indian, Pakistani, “White and Asian” (WA), and “Any Other Asian” (AOA) were characterized via three main methods: principal components analysis (PCA), ADMIXTURE, and support vector machines (SVM). For each method, the UKB data were projected onto or trained on data from a globally distributed reference panel (1KG) to characterize genetically how UKB-BIP participants align with other South Asian groups and to identify underlying genetic patterns of participants with “ambiguously defined” ethnicities (WA and AOA). For WA and AOA participants, ascertaining genetic similarities with broader groups (of potentially differing ethnic labels) may be the only way that these participants are routinely included in future genetics research, since discretizing populations into ethnic groupings of small sample sizes typically results in their exclusion from genetic analyses [5, 16, 20]. Therefore, in addition to the PCA and ADMIXTURE assessments, an SVM was trained to quantitatively integrate information from the previously calculated PC scores and infer both population-level affinities and subcontinental assignments for each AOA and WA participant (see Methods section and SI Section 2.2).

#### Ascertainment bias in the UKB SNP array

To conduct these analyses, the QC-ed UKB SNP array (see Methods) was merged with the 1KG data (*n* = 668,051 shared SNPs). The 1KG data were used to calculate 20 PCs and the UKB data were projected onto this PC space (FRAPOSA [23]). PC plots were visually assessed to identify large-scale patterns within and between the UKB groups.

During the initial visual inspections, however, PCA results indicated that Utah residents with Northern and Western European ancestry (CEU) and British in England and Scotland (GBR) samples explained most of the genetic variance within the 1KG reference data (prior to projection of the UKB data) (Fig. 1), prompting further investigation of the data qualities. Several assessments were employed to identify the root cause(s) driving the inflated European variance observed in the PC plots, ultimately indicating ascertainment bias of the UKB genotype array as the primary cause (see Methods, SI Section 1.2.3, and Figures S1.3 – S1.5). Because the array was designed to highlight European genetic variation, most of the rare alleles on the array are found at higher frequencies within European groups (Fig. S1.2), artificially inflating the magnitude of European genetic heterogeneity relative to other globally distributed populations. When the UKB and 1KG data are intersected for population genetic analyses, globally representative data (1KG) gets reduced to the variants that, within PCA results, fail to adequately capture variation among samples with non-European genetic ancestry (i.e., the formerly globally representative sample becomes primarily representative of genetic variation within the British Isles). Therefore, for all subsequent analyses (described below), the original intersection of QC-ed SNPs within both UKB and 1KG data was further reduced to a subset chosen to reflect broad patterns of expected global genetic diversity (*n* = 199,495) following more stringent MAF thresholds and expulsion of variants that failed F_ST_ assessments (See Methods section and SI Section 1.2.3 for more details).

#### UKB Bangladeshi, Indian, and Pakistani Principal Components Analysis (PCA)

UKB Bangladeshi, Indian, and Pakistani (BIP) participants generally cluster closer to one another and to 1KG South Asian reference groups than to other reference populations included in the global PCA. The UKB Indian group demonstrates the highest variance in scores across PCs, suggesting the highest levels of genetic diversity of the UKB BIP groups (Fig. 2; SI Section 1.3.1, Fig. S1.6-S1.8), although they also comprise the largest sub-sample, possibly confounding this interpretation. Across PCs, the Pakistani scores fall within the Indian ranges, making them largely indistinguishable from Indian-identifying participants. In 5 of the 20 calculated PCs, the Indian score distributions have more than one peak, suggesting population substructure that variably aligns with the Bangladeshi and Pakistani mean scores. Additionally, substantial population structure within the Indian group is visible on PC 7, which is driven by variation in South Asian reference populations. Along this PC, the Gujarati Indians sampled from Houston, TX (GIH) exhibit two peaks, one of which is far removed from other South Asian populations (this GIH substructure has previously been described by Sengupta et al. and Reich et al. [24, 25]). Clusters of UKB Indian participants can be found within both of the GIH peaks, as well as clinally distributed between them. Bangladeshi PC scores typically fall entirely within the Indian ranges, but with means closer to the East and Southeast Asian groups. This geographic trend (a West-East cline) is further reinforced where Pakistani and Bangladeshi mean scores diverge, as the Bangladeshi group tends to fall closer to the East and Southeast Asians while the Pakistani tend to fall closer to the Europeans (see Fig.S1.5 for examples).

**Figure 2.**
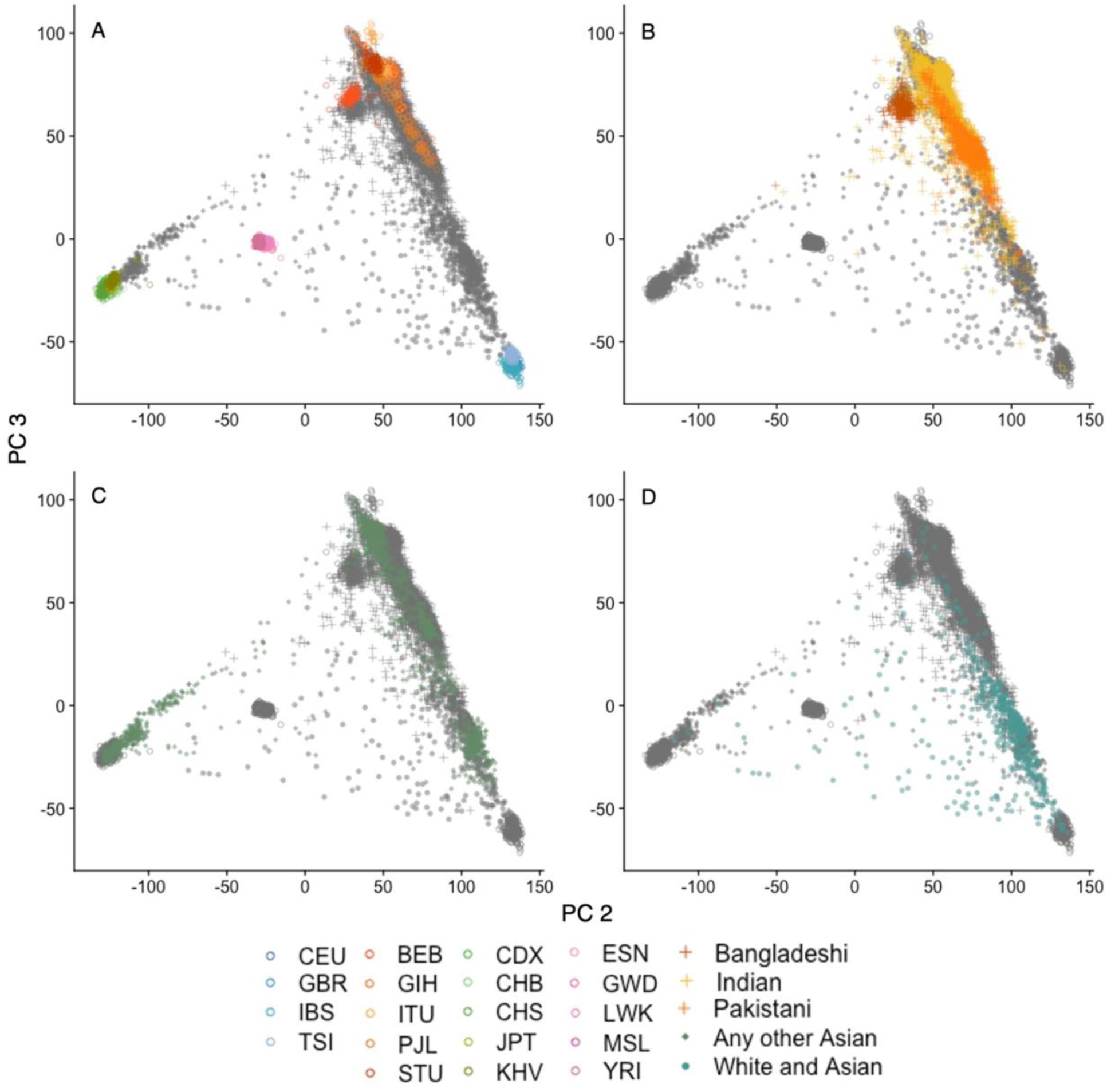
UKB self-selected ethnicity groups (UDI 21000) projected onto 1KG PCs 2 and 3. **A)** 1KG data used in the PCA, colored by reference population. **B – D)** UKB data projected onto 1KG PC space, with **B)** Bangladeshi, Indian, and Pakistani identifying participants colored; **C)** “Any other Asian” identifying participants colored; and **D)** “White and Asian” identifying participants colored.

#### UKB Bangladeshi, Indian, and Pakistani ADMIXTURE

Supervised ADMIXTURE analyses (informed by the unsupervised ADMIXTURE analyses of the 1KG references) were conducted for *k* = 3:9 and *pong* was utilized to visualize these results (Fig. 3; SI Section 1.3.3). Throughout lower and intermediate values of *k* (e.g., 4 – 6), the Indian and Pakistani groups are predominantly characterized by European and broadly South Asian components. At these *k*’s, Indian and Pakistani participants share similar proportions of these two clusters except for a small subset of Indian participants who have much higher South Asian affinities (Fig. S1.13, k = 4 – 6). However, at *k* = 7, the GIH reference group appears as its own cluster, replicating the South Asian substructure demonstrated in the PCA. This GIH-associated component is found throughout BIP groups, though in the highest proportion in the subset of Indian participants previously characterized by their higher South Asian affinity (Fig. S1.13, *k* = 7 – 9). The Pakistani participants tend to fall within the ranges of cluster proportions that also characterize the Indian participants (generally, a combination of European, broad South Asian, and GIH-associated clusters). Throughout *k*’s, the Bangladeshi participants demonstrate higher affinities with the East Asian and Southeast Asian clusters than either of the other UKB-BIP groups, and generally align with the 1KG BEB reference group. Group average percentages for cluster affinities across *k*’s can be found in Table S1.10.

**Figure 3.**
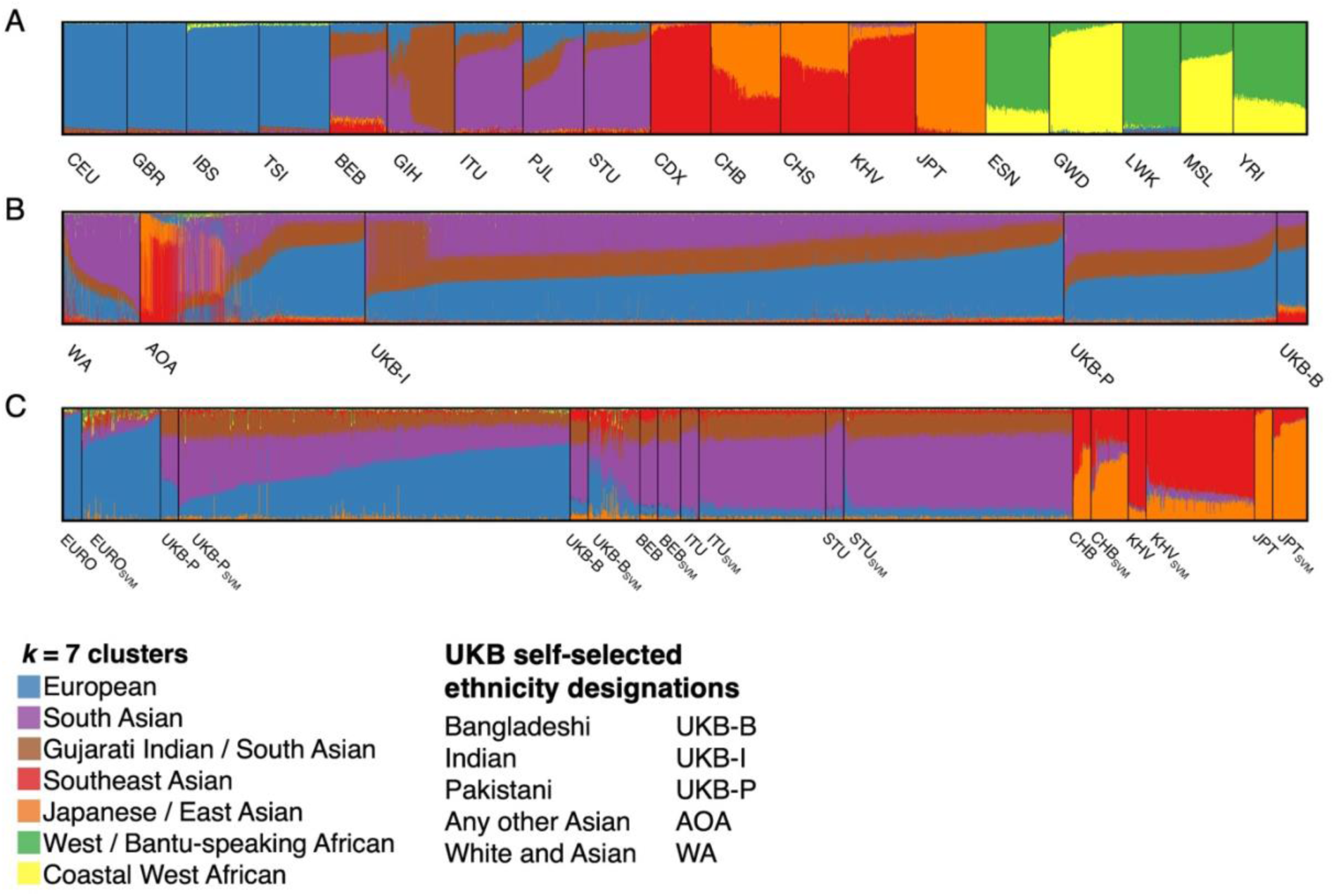
Supervised ADMIXTURE output trained on 1KG data, for *k* = 7. **A)** 1KG populations used in the unsupervised analyses. **B)** Supervised ADMIXTURE output grouped by the original UKB self-selected ethnicity labels (UDI 21000). **C)** Supervised ADMIXTURE output grouped by SVM training group (*n* = 30; left) and respective predicted group classifications (right) for UKB “White and Asian” and “Any other Asian” identifying participants. Only groups with 30 or more SVM classifications are included in this visualization. UKB Indian-identifying participants were not included in the SVM analyses and are therefore not included as a classification option here.

#### UKB “Any Other Asian”

Results of PCA, ADMIXTURE, and SVM classifications consistently demonstrate that UKB participants who self-identify as “Any Other Asian” (AOA) (*n* = 1,614) do not comprise a genetically homogeneous group (as the ethnicity label may imply). PCA results indicate that the AOA group mostly consists of participants who genetically align with Asian reference populations. Across PC’s, score distributions highlight substructure within this grouping, especially along PC’s driven by Asian genetic variance (e.g., PC’s 4, 7, and 8; Fig. 1; SI Section 1.2.1). At least four clusters can be visualized in the PCA plots, consistently aligning with Japanese sampled in Tokyo (JPT), Chinese Dai sampled in Xishuangbanna (CHB), and the conglomerate of South Asian clusters, as well as a cluster of participants that tend to fall near or between the Kinh Vietnamese sampled in Ho Chi Minh City (KHV) and Southern Han Chinese (CHS) samples. However, not all participants can be characterized solely by Asian substructure. One participant consistently aligns with 1KG African reference groups, and European ancestry is also possible for a subset of the participants (Fig. 2). ADMIXTURE analyses further replicate these results, again indicating substantial genetic substructure within this ethnic label across all values of *k* (Fig. 3B) and demonstrating that a small portion of AOA participants are more genetically similar to European reference populations as opposed to Asian reference populations (Table S1.11).

SVM classifications reiterate the genetic structure within this ethnic label, inferring affinities with reference groups across the globe (classifications at ≥ 0, ≥ 0.75, and ≥ 0.9 probabilities are provided in Table S1.8). Using four subcontinental probabilities (with summed classification probabilities ≥ 0.7), we are able to classify 1,408 AOA participants. Specifically, 2.8% of the total AOA sample classify as European, 20.6% as East Asian, and 64.2% as South Asian (Table S1.9). These subcontinental classifications are derived from grouping the population-level assignments, for example the ‘East Asian’ classification is created by summing CDX, CHB, CHS, KHV, and JPT classification probabilities. The majority of AOA participants were classified as Bangladeshi or Pakistani, as UKB Indians were not incorporated into the training data; (see SI Section 1.3.2 for more information), and Indian Telugu (ITU) or Sri Lankan Tamil (STU) at lower classification probabilities. Of these participants, 159/208 that were classified as ITU and 309/373 classified as STU (classification probability > 0) report Sri Lanka as their region of origin. In total, 500 AOA participants report Sri Lanka as their region of origin, followed by Mauritius (*n* = 85), Afghanistan (*n* = 72), Kenya (*n* = 59), Iran (*n* = 33), and Malaysia (*n* = 31) (see Fig.S1.11 for further details). Notably, no participants were classified as Punjabi from Pakistan (PJL) at any classification probability threshold. Of the 1,613 AOA participants included in these analyses, 1,036 align at the subcontinental level with South Asian samples.

#### UKB “White and Asian”

While AOA results indicate the presence of multiple structured populations within a single ethnic label, “White and Asian” (WA) analyses suggest a clinal genetic pattern between European and Asian populations (Fig. 2D). ADMIXTURE analyses also indicate high levels of European affinity throughout values of *k* (Fig. 3; Fig.S1.14; Table S1.11). SVM classifications predominantly place WA participants as European or UKB Pakistani, and to a lesser extent UKB Bangladeshi (Table S1.8; Fig. 4). Of participants with an SVM classification of UKB Pakistani (classification probability > 0), 54/93 report India as their region of origin and 11/93 report Pakistan as their region of origin. India and Pakistan reflect the most common region of origin for WA participants, followed by Myanmar (*n* = 7), The Guianas (*n* = 5), Germany (*n* = 4), and Sri Lanka (*n* = 4) (see Fig.S1.10 for further details). Probabilities summed at the subcontinental level classified 16.8% of WA participants as European, 1.5% as East Asian, and 62.8% as South Asian (Table S1.9). Of the 549 WA participants included in these analyses, 345 align at the subcontinental level with South Asian samples.

**Figure 4.**
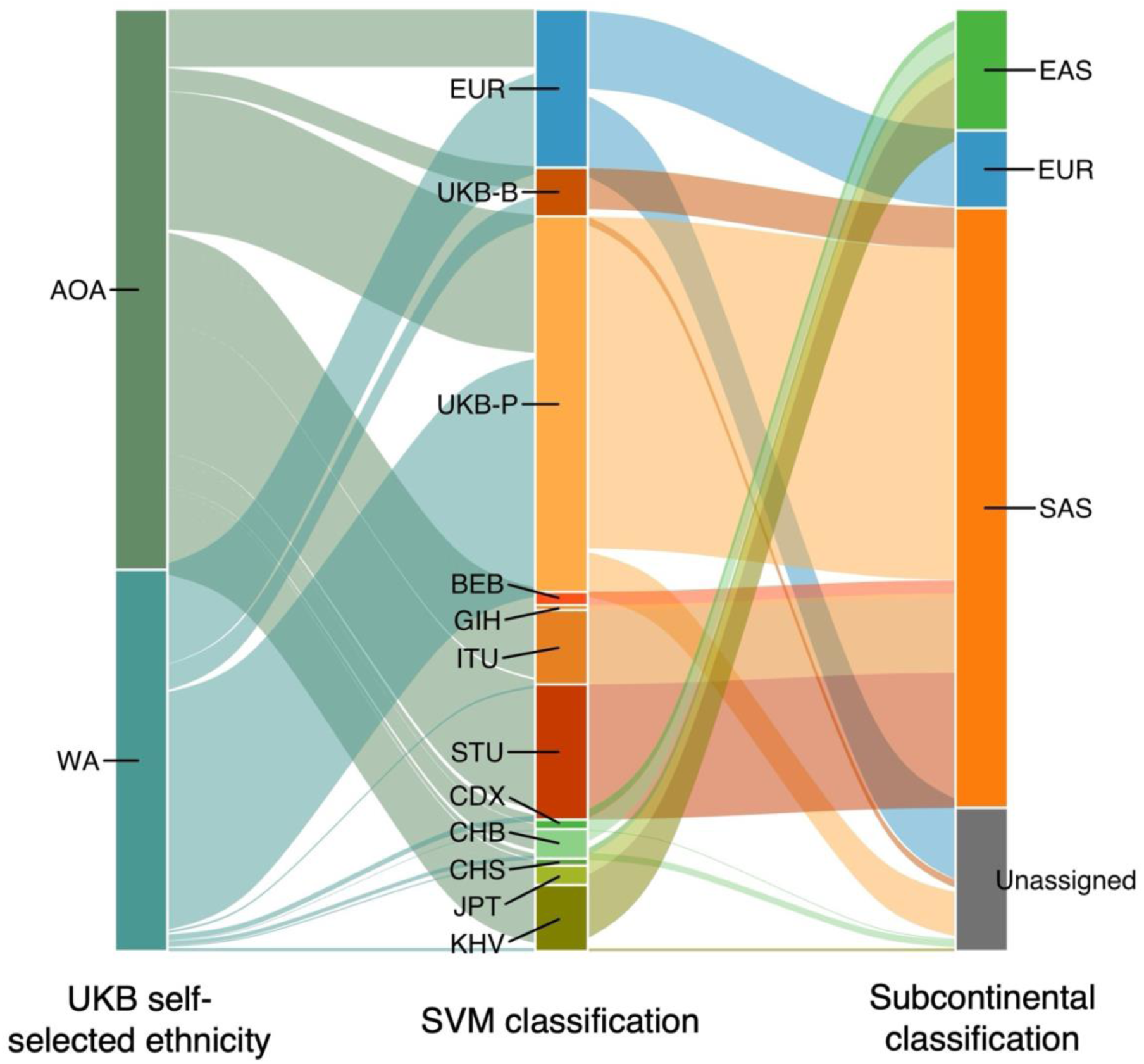
Relationships between group labels and genetic affinities throughout the SVM pipeline, for UKB participants who selected “Any other Asian” (AOA) or “White and Asian” (WA) as their ethnicity. Participants with self-selected AOA or WA ethnic backgrounds (UKB data-field 21000) (left column) were first assigned a group membership to one of the 13 reference populations used to train the SVM (AFR not pictured), based on the highest classification probability assigned (middle column). Classification probabilities for each participant were grouped by subcontinental region and summed. Those with a summed subcontinental classification probability ≥ 0.7 were assigned to that group (right column). AOA and WA participants assigned to the South Asian (SAS) group were added to the UKB-BIP sample to create a new UKB South Asian sample.

Cumulatively, the dispersed PCA scores, high European affinity demonstrated by ADMIXTURE, and low population-level SVM classification probabilities suggest that WA participants either have high levels of both European and Asian ancestry (aligning with an “admixed” interpretation of the label “White and Asian”) or that these participants share genetic affinities with populations not represented in the reference data (e.g., Central and/or West Asian groups). Future analyses of haplotype data or the inclusion of more representative reference data will be necessary to disentangle these two hypotheses.

#### Residual Population Stratification

While comparative analyses collectively suggest that UKB-BIP participants are genetically most similar to non-UK South Asian groups (i.e., the 1KG South Asian reference populations), ADMIXTURE results, specifically, indicate slightly higher European affinities within all UKB groups assessed than the 1KG South Asian samples (Fig. 3) and there are visible (and quantifiable) discrepancies between UKB WA and AOA ADMIXTURE proportions compared to the 1KG SVM associated training group (Fig. 3; Fig. S1.12 – S1.14; Tables S1.10 and S1.11). It is unclear if this is an artifact of the genotype array or a true signal of higher shared genetic ancestry between UKB participants and other Europeans.

##### Disclaimer regarding genetic affinity analyses

It should be emphasized that human genetic diversity is not categorical, and the analyses utilized here leverage the incredibly small amount of genetic diversity that differs among populations [25, 26]. The co-opting of statistical methods designed to classify and discriminate between discrete data towards analyzing human genetic variation is employed here only to facilitate the incorporation of typically excluded participants in genetic studies. The terms used in the Methods and Results sections (e.g., “classification”) reflect the statistical procedures conducted and are not meant to impose identities or group affiliations outside of this context.

### Environmentally-adjusted GWAS and PGS

#### Environmental covariates

Height is a composite phenotype made up of many anatomical regions, structures, and tissues. The primary contributor to adult stature, however, is the skeletal system. Bone mass and joint integrity (and consequently, height) peak at around 30 years of age [28, 29] before gradually breaking down over the next several decades of life. Non-genetic factors that influence an individual’s peak bone mass include primarily the environment experienced during ontogeny. To account for these effects, variables encompassing diet, general health, activity patterns, exposures, and socioeconomic status were requested from the data available in the UKB. After growth has ceased, systemic health and maintenance become the prominent factors concerning skeletal mass and morphology. Variables that affect bone mineralization, joint function, overall health, and the likelihood of incurring fractures were included to account for post-ontogenetic environmental influences.

Following assessments of the quality, missingness, and correlations among the requested environmental variables (Methods), nine data fields were selected for inclusion in the GWAS (Table 2; Figure S2.2). To test if these specific environmental variables are significantly associated with height, and to identify the variance in height explained by additive environmental components alone, an ordinary least squares (OLS) regression model was tested. The full UKB South Asian sample with un-imputed data (*n* = 7,331) was used for this assessment.

**Table 2.**
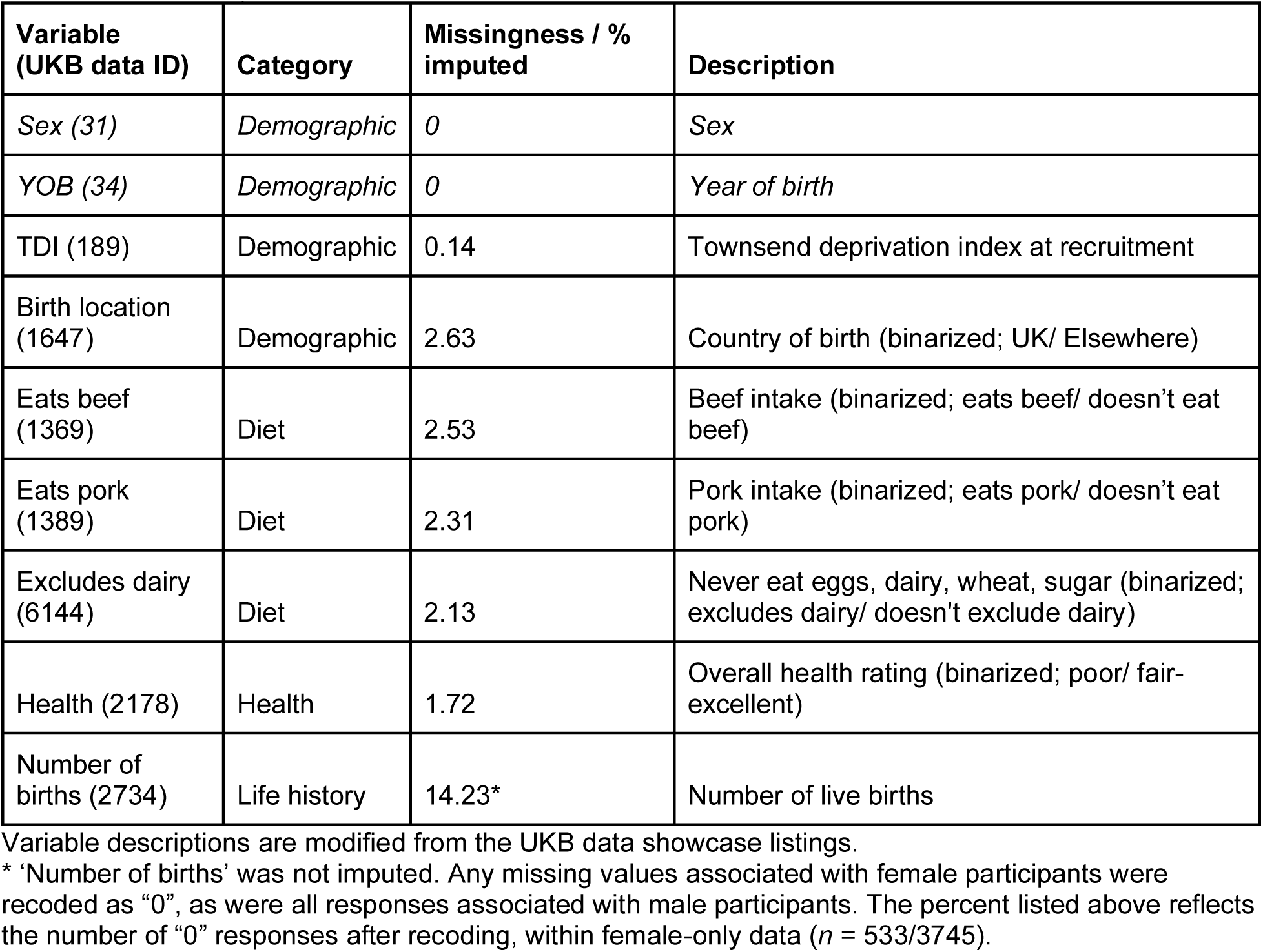
Environmental variables included as covariates in GWAS_null_ (italicized) and/or GWAS_env_ (all variables listed here).

| Variable (UKB data ID) | Category | Missingness / % imputed | Description |
| --- | --- | --- | --- |
| Sex (31) | *Demographic* | 0 | Sex |
| YOB (34) | *Demographic* | 0 | Year of birth |
| TDI (189) | Demographic | 0.14 | Townsend deprivation index at recruitment |
| Birth location (1647) | Demographic | 2.63 | Country of birth (binarized; UK/ Elsewhere) |
| Eats beef (1369) | Diet | 2.53 | Beef intake (binarized; eats beef/ doesn't eat beef) |
| Eats pork (1389) | Diet | 2.31 | Pork intake (binarized; eats pork/ doesn't eat pork) |
| Excludes dairy (6144) | Diet | 2.13 | Never eat eggs, dairy, wheat, sugar (binarized; excludes dairy/ doesn't exclude dairy) |
| Health (2178) | Health | 1.72 | Overall health rating (binarized; poor/ fair-excellent) |
| Number of births (2734) | Life history | 14.23* | Number of live births |

Variable descriptions are modified from the UKB data showcase listings.
\* 'Number of births' was not imputed. Any missing values associated with female participants were recoded as "0", as were all responses associated with male participants. The percent listed above reflects the number of "0" responses after recoding, within female-only data ( $n = 533/3745$ ).

The environment-only model explains 6 and 8% of the variance in height in males and females, respectively (Table 3; Table S2.2). When applied to the combined sex sample, all variables meet significance (p < 0.01) except for ‘Number of live births’ (Table S2.1), though visualization of the effect does suggest a weak, negative association with height (Figures S2.4 – S2.5). Variables with the strongest effects on height include sex, ‘Birth location’ (whether or not the participant was born in the UK), and ‘Excludes dairy’ (whether or not the participant includes dairy in their diet). Dietary variables reflecting meat consumption, and ‘Health’ (a self-assessed overall health indicator) are associated with moderate effects on height, while Townsend Deprivation Index (‘TDI’) and year of birth (‘YOB’) confer the weakest effects of all significant covariates included in the model.

**Table 3.** Linear regression summary statistics for non-genetic predictors on height.

| Model covariates | Sample | Adj. $R^2$ | $p$ |
| --- | --- | --- | --- |
| Sex + TDI + YOB + Birth location + Eats pork + Eats beef + Excludes dairy + Health + Number of births | All ($n = 7,331$) | 0.57 | $< 2.2 \times 10^{-16}$ |
| TDI + YOB + Birth location + Eats pork + Eats beef + Excludes dairy + Health + Number of births | F ($n = 3,430$) | 0.08 | $< 2.2 \times 10^{-16}$ |
| TDI + YOB + Birth location + Eats pork + Eats beef + Excludes dairy + Health | M ($n = 3,901$) | 0.06 | $< 2.2 \times 10^{-16}$ |

#### Height GWAS models

To assess how environmental covariate inclusion may affect GWAS results, two GWAS models were run: GWAS_null_, which includes typical covariates used in previously published height GWAS (e.g., age, sex, and genetic PCs); and GWAS_env_, which incorporates environmental covariates hypothesized or previously demonstrated to influence growth and/or systemic health, in addition to the covariates in the “null” model. These covariates can be found in Table 2. The GWAS were run on a subset of the full sample (*n* = 6,954) as 1000 randomly selected participants were “left-out” for PGS assessments and some participants did not have standing height data. SNP significance and effect sizes output by both GWAS models were compared.

#### Variant significance and effect size comparisons

Comparisons of SNP significance between GWAS_null_ and GWAS_env_ outputs demonstrate increased detection of tag-SNPs in GWAS_env_ (Table 4; Figure 5). This increase in associated SNPs did not occur in a uniform manner, however, as the incorporation of environmental covariates decreased the significance of some associations relative to GWAS_null_ while increasing others.

**Figure 5.**
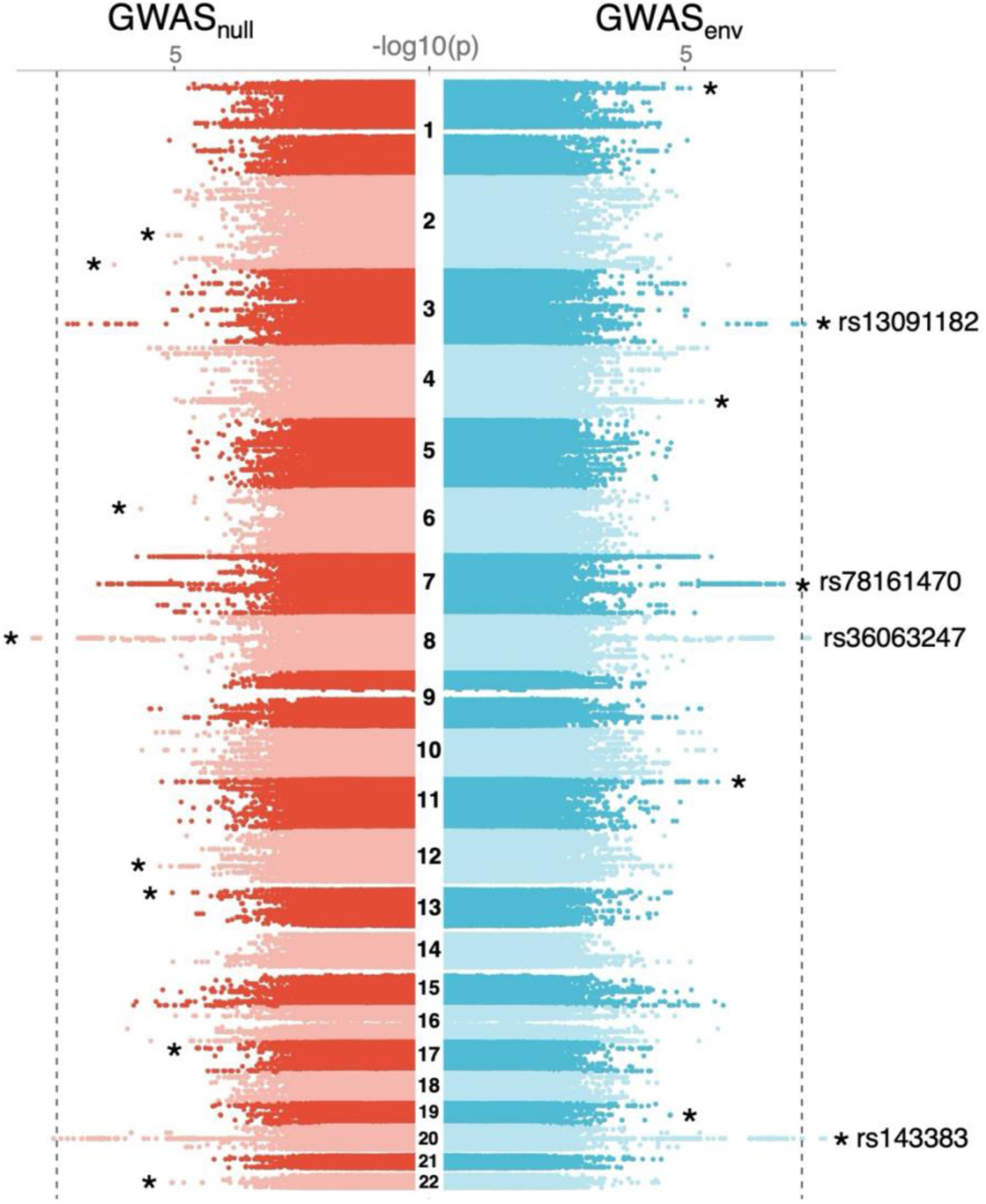
Manhattan plots of significantly associated SNPs resulting from GWAS_null_ (left panel) and GWAS_env_ (right panel). A significance threshold of *p* < 5^-8^ is designated by the dashed lines. The rsID for the variants with the lowest p-value per peak is listed in the GWAS_env_ panel. Regions with inflated significance in GWAS_null_ that are reduced in GWAS_env_ are designated by asterisks in the GWAS_null_ panel. Alternatively, regions with increased significance in GWAS_env_ in comparison to GWAS_null_ are designated by asterisks in the GWAS_env_ panel.

**Table 4.** Number of SNPs meeting different significance thresholds between GWAS models.

| $p <$ | $10^{-5}$ | $10^{-6}$ | $10^{-7}$ | $5^{-8}$ |
| --- | --- | --- | --- | --- |
| GWAS<sub>null</sub> | 530 | 248 | 11 | 8 |
| GWAS<sub>env</sub> | 558 | 294 | 17 | 11 |

Changes in SNP effect size between GWAS outputs were also investigated, demonstrating both reductions and increases in the effects of significant variants. Comparisons between SNP effect sizes between GWAS_env_ (β_env_) and GWAS_null_ (β_null_) were restricted to variants with *p* < 10^-6^ output by GWAS_env_ (*n* = 294), as these reasonably represent the most reliable results of the variants assessed (though it should be noted that these do not meet the standard threshold for genome-wide significance). As such, no sign flips (i.e., changes in effect directionality) between β_null_ and β_env_ were detected for SNPs meeting this significance threshold. However, of these 294 variants, 131 associated effect sizes were strengthened (i.e., 2 positive values increased, and 129 negative values decreased), 163 effects were attenuated (i.e., 98 positive values were reduced, and 65 negative values were increased). All β standard errors decreased in GWAS_env_ relative to GWAS_null_, indicating more precise β estimates (Tables S2.4 and S2.5). The distribution of effect sizes differs as well, primarily with regards to SNPs with large, positive effects (Fig. S2.7). The largest shifts in effect size distributions can be found in chromosomes 7 and 8 (Fig. S2.7). Interestingly, none of the chromosome 7 SNPs meeting the significance threshold for these assessments are published in the GWAS catalog for associations with height.

#### Heritability of environmental covariates

The heritability of each environmental covariate used in the GWAS_env_ model was investigated to ascertain the potential magnitude of shared genetic causality with height. As of the submission date of this manuscript, the only published SNP-based heritability (*h^2^*) estimates available for our variables of interest were those presented in the Neale Lab SNP-Heritability Browser [http://www.nealelab.is/uk-biobank/], which are derived solely from European UKB participants. The dietary variables published by the Neale Lab have low levels of *h^2^*(∼0.03 – 0.04 for pork, beef, and dairy variables). Townsend deprivation index (TDI) is comparably low (∼0.03), as is “Number of live births” (∼0.06). Overall health, however, has a larger *h^2^*estimate of ∼0.1, suggesting that inclusion of this variable as a covariate may result in biased genetic effect estimates. As these *h^2^* estimates are not directly comparable to the UKB South Asian sample, we ran an additional GWAS for overall health (following the same procedures as our height GWAS) and calculated *h^2^* from these summary statistics using LDSC [18]. 1000 Genomes South Asian data were used to create the LD scores and regression weights required to use LDSC (after merging, 4,022,277 SNPs were used in this analysis). Our results suggest a lower *h^2^*estimate (GWAS_null_ = 0.08 [SE = 0.048]; GWAS_env_ = 0.05 [SE = 0.047]) than derived from the UKB European sample, however, SEs are large and 95% CI’s overlap 0 (Figure S2.11 and Table S2.16). This suggests that “overall health rating” is a (predictably) noisy variable likely capturing many proxies for actual health, and our data are underpowered for a reliable *h^2^* estimate. The low *h^2^* estimates suggest that any shared causal loci between height and health likely do not represent a large, direct genetic effect on height.

#### Polygenic score (PGS) construction and assessments

Following the GWAS, we developed one height PGS model for each of the two GWAS using LDpred2-auto [30, 31]. We assessed the overall performance of each model (GWAS_null_ PGS and GWAS_env_ PGS) by calculating *R*^2^ between PGS and height, both with and without the demographic or environmental covariates used in the associated GWAS, for UKB South Asian participants held out from the sample prior to GWAS analyses (*n* = 997). We additionally assessed sex biases in performance and investigated the effects of residual population stratification on PGS scores (see Methods section *Polygenic score (PGS) assessments and comparisons* and SI section *Environmental adjustments and population stratification*). Overall, performance is similar between GWAS_null_ PGS and GWAS_env_ PGS scores when using PGS alone, explaining 2.6% and 2.2% of the variance in height, respectively, though this difference is significant (Figure 6, Tables 5 and 6, and Table S2.13). When we incorporate the non-genetic covariates used in the discovery GWAS (i.e., all covariates excluding genetic PCs or GRM; see Table S2.13), performance of this larger model increases substantially (GWAS_null_ PGS *R*^2^ = 0.574; GWAS_env_ PGS *R*^2^ = 0.587) as expected, though this is predominantly due to including sex, since height is a sexually dimorphic phenotype. For sex-stratified samples, neither PGS score shows a significant difference in variance explained between sex, despite the higher point estimate observed in females. Inclusion of the environmental covariates additionally reduces the difference between estimates of sex-stratified performance (GWAS_null_: *R^2^*_F_ = 0.12; *R^2^*_M_ = 0.077; GWAS_env_: *R^2^*_F_ = 0.123; *R^2^*_M_ = 0.106) (Table S2.13).

**Figure 6.**
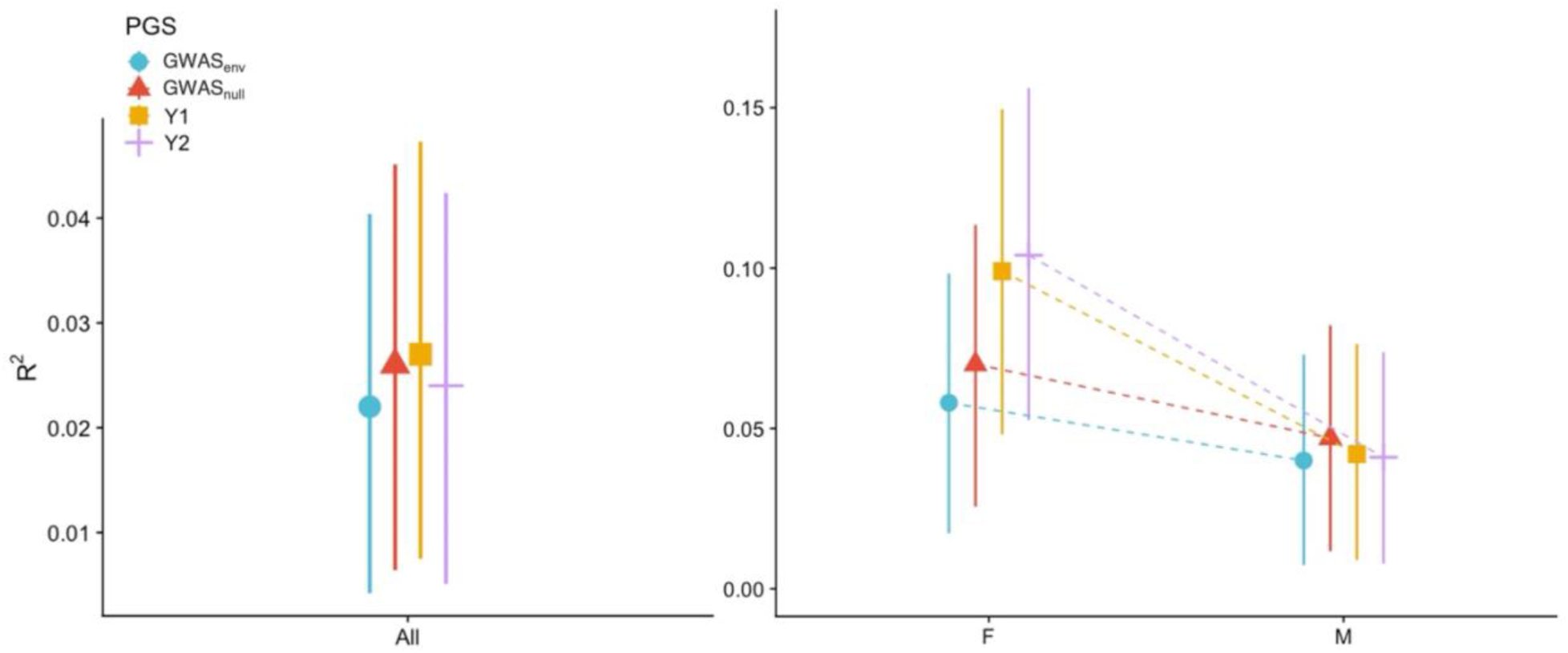
Comparison of correlations between PGS score and participant height (*n* = 997). *R^2^*values (symbols) and 95% CI (solid lines) reflect PGS performance across the full test sample (left panel) and sex stratified samples (right panel). Dashed lines connect *R^2^* estimates between sex stratified samples to help visualize biases in PGS performance. PGS models are designated by color and symbol, and are consistent across panels.

**Table 5.** Predictive performance of PGS scores. All models meet significance at *p* < 1 x 10^-5^. Extended results can be found in Tables S2.12 and S2.13.

| PGS | $R^2$ ($n = 997$) | $R^2_F$ ($n = 473$) | $R^2_M$ ($n = 524$) | $R^2_F - R^2_M$ | Fisher's $z^{**}$ | $p^{**}$ |
| --- | --- | --- | --- | --- | --- | --- |
| Y1<sup>†</sup> | 0.027 | 0.099 | 0.042 | 0.057 | 1.82 | 0.034 |
| Y2<sup>^</sup> | 0.024 | 0.104 | 0.041 | 0.063 | 2.05 | 0.020 |
| GWAS<sub>null</sub> | 0.026 | 0.07 | 0.047 | 0.023 | 0.79 | 0.216 |
| GWAS<sub>env</sub> | 0.022 | 0.058 | 0.04 | 0.018 | 0.66 | 0.254 |

<sup>†</sup>We use the subscripts "F" and "M" to indicate inclusion of females and males, respectively.
<sup>\*\*</sup>Fisher's z-statistic and associated p-value (one-tailed) derived from testing the hypothesis that $R^2_F > R^2_M$ .
R package *cocor*, function *cocor.indep.groups()* was used for these tests [32,33].
<sup>†</sup>Y1: PGS002800 – GWAS SNP intersection ( $n = 898,486$ ).
<sup>^</sup>Y2: PGS002800 – "full" SNP intersection ( $n = 1,032,672$ ). SNPs used in this PGS score reflect those in the Yengo et al. [34] PGS weight file and our imputed data (regardless of QC filtering thresholds).

**Table 6.** Results of two-tailed significance testing between PGS *R^2^*. *R^2^* estimates compared here reflect those from PGS-only models (i.e. height regressed on PGS score without additional covariates). R package *cocor*, function *cocor.dep.groups.overlap()* was used for this analysis [32,33].

| PGS | Y1 | Y2 | GWAS<sub>null</sub> | GWAS<sub>env</sub> |
| --- | --- | --- | --- | --- |
| Y1 |  | 0.0347 | 0.8947 | 0.6618 |
| Y2 | -0.0114 |  | 0.864 | 0.8967 |
| GWAS<sub>null</sub> | -0.0049 | 0.0064 |  | 0.0891 |
| GWAS<sub>env</sub> | -0.0163 | -0.0049 | -0.0113 |  |

Lower triangle) difference between $R^2$ estimates (row PGS minus column PGS). Upper triangle) two-tailed p-value of test for differences between Pearson and Filon's z transformed PGS correlations.

#### Comparisons with Yengo et al., 2022

We further tested our PGS by assessing its performance relative to a published South Asian PGS [34; PGS002800 in the PGS Catalog], which was developed from a GWAS meta-analysis (*n* = 77,890) and tested on UKB South Asian participants (*n* = 9,257). To our knowledge, this PGS equation was developed on the largest ancestry-matched dataset to our sample, and therefore represents the upper limit of predictive performance currently achievable. The PGS002800 score [35] was applied to our test sample using ESCALATOR, a PGS harmonization pipeline [36]. Although the specific participants used in the Yengo et al. [34 PGS test sample are unknown, it is likely that a large proportion are also included in our GWAS and PGS test sample, making our data appropriate for direct comparisons. For all analyses, two versions of the Yengo PGS models were developed: PGS_Y1_, which includes all SNPs shared between the Yengo et al. [34] weight file and our GWAS data, and PGS_Y2_, which additionally includes variants shared with the full GasP-imputed dataset, prior to QC filters. Performance accuracy (*R*^2^) was compared across several combinations of SNP sets, samples (e.g., sex-specific subsets), and covariate inclusion to account for differing variants included in the PGS as well as environmental variables incorporated in the training GWAS (Tables 5 and S2.12). Additionally, we assessed if PGS002800 performance was solely due to a larger GWAS sample size or could be partially explained by unadjusted environmental covariates that might inflate effect sizes.

The PGS002800 PGS_Y_ scores yield comparable results to our GWAS-derived PGS scores, with *R*^2^ point estimates ranging from 0.024 to 0.027 when applied to the full test sample (Tables 5 and S2.12), though this falls short of the reported range of 0.033 to 0.045 [34]. Although point estimates of PGS *R^2^* are larger for most Yengo-derived models, formal comparisons with the PGS *R^2^* developed here do not result in any significant differences (*at* p < 0.05) (Table 6, Table S2.14). Within sex-stratified subsets, male *R*^2^ values range from 0.041 to 0.042 while female *R*^2^ values range from 0.099 to 0.104. Unlike the two PGS scores developed in this study, PGS_Y_ scores explain significantly more variance in females than in males (*p_Y1_*= 0.034; *p_Y2_* = 0.02). When YOB is included in the model (Table S2.12), *R*^2^ values increase, explaining ∼14 – 15% of the variance in height in females and ∼7% in males.

To differentiate between factors impacting PGS performance (e.g., if accuracy is increased due to collinearity of genetic effects and un-adjusted environmental influences), each PGS score was then regressed on the GWAS_env_ environmental covariates (Table S2.15). Comparatively, a larger *R*^2^ (between PGS score and environmental covariates) suggests a higher likelihood that environmental influences are captured by the β weights used to calculate the PGS score (as opposed to directly causal genetic effects). Across models, *R*^2^ is predictably low, though point estimates are (weakly) higher for sex-combined PGS_Y_-derived scores than those resulting from the GWAS PGS scores developed here. Therefore, it is inconclusive if PGS_Y_ scores reflect a higher proportion of genetic effects that covary with environmental stratification relative to the GWAS_env_ PGS score, as this cannot be statistically supported given the resulting p-values and sex-stratified results. Additional investigation into the effects of population stratification on PGS performance can be found in SI Section “Environmental adjustments and population stratification”.

## DISCUSSION

Pre-existing, predominantly European-descent biobank data can be used for genotype-phenotype research, but extra steps must be taken to improve the quality of the genetic data for non-European participants. For example, as of the submission of the current study, the UK Biobank only provides genetic ancestry information for ‘White British’ participants (data field 22006), leaving the characterization of genetic affinities among non-‘White’-identifying participants to researchers. This can result in discrepancies across studies with regards to participant inclusion, and therefore, complicate the utility of out-of-sample results and inferences. By investigating the intersection of sociocultural labels and genotype data, we increase the UKB South Asian sample size by 20% (originally only comprised of Bangladeshi, Indian, and Pakistani participants), facilitating the development of larger UKB Asian analytic groups. Increasing sample sizes, including the diversification of biobank demographics, is only a partial solution. We demonstrate that including relevant environmental covariates refines the estimates of GWAS SNP effects and reduces sex-biases in PGS performance. By taking a more intensive data quality-centered approach to research design, we are able to generate PGS models that perform comparably to those trained on datasets incorporating ten times the number of participants.

### Genetic affinities of UKB South Asian participants

While the UKB ethnic categories are very limited, we find substantial genetic diversity among participants within the same label, demonstrating the limited correspondence between sociocultural labels and genetic affinities. Geopolitically-defined ethnic labels within the UKB (e.g., BIP) have more consistent patterns of genetic affinities than more ambiguous labels such as AOA or WA. While broad trends are apparent and distinguish the genetic stratification between WA and AOA groups at large, there are no definitive indicators demonstrated by the genetic data that explain participant identities or cultural affiliations at the individual level. Individual participants who report the same geographic region of origin and are assigned to the same SVM classification choose to identify as AOA versus WA (of the available options) for reasons not predictable by genetic analyses. For example, our SVM results show that a higher proportion of WA than AOA participants classify as European or UKB Pakistani, however within SVM classifications, the same regions of origin are reflected regardless of participant ethnic identity (e.g., most of the WA participants classified as UKB Bangladeshi are from India, Myanmar, The Guianas, Malaysia, and Mauritius, and most of the AOA participants classified as UKB Bangladeshi are similarly from Mauritius, Myanmar, Nepal, Sri Lanka, and Malaysia [Figures S1.10 and S1.11]). This highlights, and reiterates, that ethnic identity, genetic stratification, and shared ancestry are distinct participant descriptors that cannot be used *a priori* as proxies for one another. Therefore, ethnic groupings may represent an adequate starting point for preliminary sample selection but cannot be used, unaltered, as a homogenous or distinct genetic sample.

Additionally, we replicate previously reported trends regarding South Asian genetic affinities, particularly with regards to higher genetic similarity of UKB Pakistani participants to Europeans (Fig.3; Fig.S1.8). Based on previous research [25, 37], this is likely a result of shared ancestry derived from the prehistoric ‘Ancestral North Indian’ (ANI) population [25, 37] rather than more recent gene flow events. More recent population movements are reflected when comparing

SVM classification results with participants’ region of origin (Figures S1.9 – S1.11), exemplifying links with common diaspora communities. For example, East African countries (Uganda, Kenya, Tanzania) are predominantly comprised of participants classified via SVM as UKB Pakistani or Indian Telugu from the UK, reflecting the 19^th^ century enforced migrations of indentured servants from India, which heavily recruited from Punjab [38]. Regions of origin in the Caribbean (e.g., The Guianas) exhibit a higher proportion of participants classified as Bengali from Bangladesh, UKB Bangladeshi, and Gujarati Indians from Houston, similarly reflecting 19^th^ century enforced migration patterns of indentured servitude that derived individuals mainly from the Northeast states of India [38].

To our knowledge, we are the first group to counteract the effects of genotype array ascertainment bias in the UKB data, therefore genetically inferring a sample that is likely to be better matched with (i.e. more genetically similar to) South Asians outside of the UKB cohort. We find that typical MAF filtering thresholds of 0.01 – 0.05 (meant to reduce the effects of genotyping error) are not adequate for quality-control processing the Affymetrix array data in UKB, when the genotype data are being used to identify genetic patterns of non-European-ancestries and assess relationships with non-UKB reference data.

### Environmentally-adjusted GWAS and PGS development

Genetic associations with height are typically assessed via a linear regression model that is purely additive, ignoring epistasis, as well as gene-environment interactions (GxE). Currently, model simplification is deemed justifiable due to limitations, such as the low power of GxE tests, measurement error of the environmental variable of interest, and a lack of knowledge of the specific biological pathways underlying the interactive effects [39]. Once identified, a GxE feature can be integrated into GWAS modeling simply by entering the interaction term; however, because of the inherently non-experimental nature of human GxE discovery research, it has been difficult to disentangle confounding from causality [40]. Regardless, for complex traits that have been studied, environmental influences are relevant, significant, and often ignored [21, 41].

There are many environmental exposures that have been repeatedly demonstrated to affect human stature [42,43,44,45,46,47,48,49,50,51,52]. Yet most height GWAS only incorporate sex, age, and a genetic relatedness matrix (GRM) and/or simply the first 10 PCs to account for population stratification and adjust for confounders (e.g.: Jeon et al., 2024; Sohail et al., 2023; Wojcik et al., 2019; Yengo et al., 2022; Zoledziewska et al., 2015 [34, 53, 54, 55, 56]). In fact, a primary reason for the accessibility of ample height data in medical records is *because of the relationship between height, environmental factors, and health* including “cumulative net nutrition, biological deprivation, and standard of living between and within populations” [50: 1]. The bulk of population-level differences may be captured by within-sample PCs and GRMs, but it is unlikely that they capture fine-level environmental effects, and it has been demonstrated that controlling for population stratification in this manner isn’t always sufficient for highly polygenic traits [57].

### Environmental covariates

The addition of environmental covariates provides an opportunity to balance the introduction of precision variables with the potential to induce colliders due to shared architecture with a downstream outcome variable being used as a covariate in the model. The impact of collider bias, collinearity, and confounding on the identification and magnitude of direct genetic effects often act as justification for excluding heritable environmental covariates from GWAS and PGS development altogether [59], albeit with notable exceptions (e.g., Hoffmann et al., 2018 [60]). Here, we sought to test the introduction of environmental covariates that limit the coheritability that could drive potential bias. We focused on traits with limited absolute heritability to reduce the potential for colliders in our dataset. As we demonstrate, each of the environmental variables alone are associated with effects on the order of centimeters, whereas the largest singular allelic effect output by the GWAS_null_ or GWAS_env_ is 1.6mm (for *p* <10^-6^). This then argues for a large non-genetic component to phenotypic variability in contrast to the marginal genetic contribution per-locus. While the entirety of these effects cannot yet be fully controlled for, steps can be implemented to mitigate them. When selecting covariates for inclusion in the GWAS_env_ model, collinearity can be addressed by assessing correlations among the collection of potential covariates and removing all but one variable in a highly correlated cluster (see Methods section “Environmental covariates” and SI section 2.2 for details on how this was implemented here).

To reduce the likelihood of introducing colliders into a GWAS model, both the associations and direction of hypothesized causal effects among the phenotype of interest and possible model covariates need to be considered. For collider bias to occur here, height must have an associated effect on a covariate used in the GWAS model. It is plausible that height has an indirect and weak effect on TDI due to sociocultural contexts. Therefore, it is also possible that an endophenotype of height could be a collider, but we assume that these specific effects are likely much smaller than the collective impact of overall health on height. Prior work has demonstrated that the strength of collider bias should be almost perfectly inverse to the association between collider and exposure variables [61]. If collider bias is introduced, and our assumptions about the relative magnitude of causal effects of height on TDI or health are true, then we can expect for these biases to be minimal. Therefore, we argue that height is unlikely to exert more than minimal influence on TDI or health measures. We note that, while previous estimates of perceived health may contribute significant heritability in the white British population in UK Biobank, in our own measures we did not observe this same trend in South Asians, suggesting a minimal effect to no effect in our population of interest. However, due to the limitations of colliders within causal models, these assumptions cannot be empirically confirmed within our dataset. Further discussion on the implications of genetic colliders and confounders introduced by the model covariates can be found in SI Section 2.5 “Extended Discussion: Heritability of environmental covariates”.

Lastly, we do note that the UKB only provides data on adult participants, reflecting the participants’ recent behavior and what can be recalled from childhood. It is assumed that in most cases the participant responses regarding current diet and socioeconomic status do not deviate significantly from childhood patterns, though this cannot be explicitly tested within the data. There is some research to indicate that this is a reasonable assumption, though it is a limitation of the study [65, 66].

#### GWAS and PGS comparisons

We demonstrate that including select environmental covariates in the model refines GWAS results; specifically, we increased variant detection, reduce genomic inflation (smaller values of λ) and reduce effect size (β) standard errors when including environmental covariates in our GWAS_env_. Taken together, this suggests that: 1) adjusting for environmental influences reduces some of the noise that may be mitigating direct genetic effects; 2) the significance of some variants may be falsely inflated when environmental confounders aren’t adjusted for, leading to misidentification of causal genetic loci (e.g., the genetic variant is not causal but covaries with environmental variation); or 3) by incorporating environmental adjustments, SNP β values are more stable (i.e., have lower uncertainty) and likely reflect more accurate estimates of “true” effects. Furthermore, SNPs with large deviations in significance between the GWAS here may represent interactions with one or more of the environmental variables included in GWAS_env_. Additional investigation into these variants may generate candidate SNPs for future GxE research.

Conversely, despite a higher number of significant SNPs at different p-value thresholds, point estimates of our GWAS_env_ PGS explains less phenotypic variance than the GWAS_null_ PGS when considering PGS scores, alone. We additionally compared the performance of our GWAS derived PGS models relative to that of Yengo et al., [34], which was trained on a meta-analysis comprising the largest published South Asian sample. All models perform comparably, even though the Yengo et al. PGS were trained on a sample size that is over ten times larger than ours, suggesting that increasing sample size is not the full solution to improving PGS performance. When stratifying PGS performance by sex, the Yengo-derived PGS scores tend to explain more of the variance in height, however, performance improvement appears to be sex-biased. Within all models, performance is higher for females relative to males, but this discrepancy is most reduced in GWAS_env_ PGS results. Interestingly, PGS_Y1_ performed better than PGS_Y2_ (Table S2.11), although the latter incorporates a larger proportion of possible variants in the PGS002800 scoring file. It is possible that this is simply due to the quality of the imputed data, as PGS_Y1_ only included the intersection of SNPs meeting our GWAS inclusion info score threshold of 0.8. However, it also could be the case that the additional ∼100k SNPs in PGS_Y2_ don’t add much to model performance, which could be due to the multitude of factors generally implicated in PGS portability.

## CONCLUSIONS

Even though ancestry-diverse biobank cohorts remain small, there are steps we can take to maximize the utility of these samples within the context of genotype-phenotype research. In addition to commonly applied adjustments, e.g., incorporating LD-informed methods, some fundamental aspects of research design can be implemented to improve the quality of the results. We argue that the improving precision in trait measurement and ancestry assignment, as well as including environmental covariates facilitate the performance of PGS. If the aim of PGS implementation is not just to predict a trait expression or health outcome, but to understand *why* the outcome is predicted, then covariate effects need to be interpretable for individuals across populations. As such, if biobank-level data is accessible, it should be standard practice for GWASs of height to incorporate, at a minimum, covariates that may adjust for within-population variance in height explained by nutritional or socioeconomic variation amongst individuals. Such exploration remains a challenging, but likely fruitful area for new research.

## ACKNOWLEDGEMENTS

Above all others, we would like to thank the participants of the UKB for providing their personal health data towards furthering health research. Additionally, we would like to thank Graham Coop and Mark N. Grote for their guidance, critiques, and comments throughout the development of this project (and resulting manuscript).

## CODE REPOSITORIES / DATA AVAILABILITY

The results presented here will be returned to the UKB, providing comparable data to data field 22006 for South Asian participants, and will be available to approved researchers. We recommend replicating this pipeline on participants enrolled subsequent to our research agreement to continue building this sample. By characterizing, in depth, the genetic affinities of participants comprising the largest non-‘White British’-identifying sample, we hope that future researchers will be able to more easily incorporate these participants in their studies.

The scripts developed for the analyses discussed here will be housed in an open access repository at https://github.com/ccatram/AnthroGen. The projected PC scores and SVM classification probabilities will be added to the UKB database and made available to researchers approved by the UKB review panel.

## MATERIALS AND METHODS

### Sample datasets

All data used in this study were either obtained from the UK Biobank via project ID 54084 (downloaded in November, 2020) or, with regards to the 1KG Phase 3 data (build GRCh37), downloaded from the IGSR FTP site in April 2020 [62, 63]. All 1000 Genomes (“1KG”) reference groups reflective of European, Asian, and African super-populations were included except for American groups (e.g. MXL) and those with recent, complex admixture (e.g., ASW) (see Table S1.6 for further information). UKB participants were selected based on responses to UKB data field 21000 and were included in our starting sample if any of the following ethnic identities were chosen: Bangladeshi, Indian, Pakistani, Any other Asian, or White and Asian (*n* = 10,288 at the time of our data application [4,832 females, 5,456 males]; Table 1). Information regarding the UKB genotype array (Affymetrix UK Biobank Axiom® array) can be found in the UKB data showcase website (https://biobank.ndph.ox.ac.uk/showcase/), specifically “Resource 1955: SNP Quality Control information” and “Resource 807: Sample processing and preparation of DNA for genotyping”.

The UKB consists of over 7,000 available data fields, 283 of which were requested for this study (Figure S2.2 A). These 283 data fields were selected based on their known or hypothesized effects on skeletal growth and systemic health, or if the data field reflected contextual information surrounding data collection so that any potential biases introduced while obtaining the data could be investigated (Figure S2.2 B).

The data utilized in this study fall under the ethics approvals granted to the UKB. The UKB received approval from the National Information Governance Board for Health and Social Care and the National Health Service North West Centre for Research Ethics Committee (Ref: 11/NW/0382), maintains its own internal ethics board (UK Biobank Ethics Advisory Committee), and is compliant with the General Data Protection Regulation (GDPR) [14, 15]. Participants enrolled in the UKB provided electronic signed consent at the time of enrollment [14, 15].

### SNP array quality control

All genotype arrays are aligned to the GRCh37 build. After extracting samples from the UKB and removing duplicates, additional quality control (QC) filtering steps were taken to remove relatives, very rare variants, and filter out SNPs with characteristics more prone to, or indicative of, genotyping errors (Table S1.7). Genotype arrays for chromosomes 1-22 were merged (*n* = 784,256 variants) and filtered using PLINK v2.00a2.3 (--geno 0.10, –-hwe 0.000001, –-maf 0.000097, –-mind 0.90), removing 23,732 variants due to missing genotype data, 13,313 variants due to Hardy-Weinberg equilibrium threshold, and 51,860 variants due to allele frequency threshold. First and second-degree relatives were identified with KING v2.1.8 (--unrelated, –-degree 2) and removed, retaining 8,967 samples (1,321 individuals were removed due to relatedness). The array was then filtered once more to remove any remaining rare variants (--maf 0.0001) after relatives were removed, resulting in 692,466 variants remaining.

### Principal components analysis

Principal component analysis (PCA), support vector machines (SVM), and ADMIXTURE analyses were performed to characterize the genetic affinities of UKB participants who self-identify as Bangladeshi, Indian, Pakistani, “White and Asian” (WA), and “Any other Asian” (AOA). To identify if Bangladeshi-, Indian-, and Pakistani-identifying participants represent a genetically South Asian metapopulation and to elucidate the genetic affinities of AOA and WA participants, a principal component analysis (PCA) was performed using FRAPOSA and the 1KG data as a reference for global genetic diversity (Table S1.6) [23, 68].

To conduct these analyses, the QC-ed UKB array was merged with the 1KG WGS data, resulting in an intersection of 668,051 shared SNPs. The 1KG data were used to calculate 20 PCs and the UKB data were projected onto this PC space using FRAPOSA [23]. PC plots were visually assessed to identify large-scale patterns within and between the UKB groups.

However, unexpected data issues were discovered after visualizing the PC plots derived from the initial merge with 1KG data. PCA results indicated that CEU and GBR explained most of the genetic variance within the 1KG reference data, prompting further investigation of the data quality (see SI Section 1.2.3; Figures S1.3 – S1.5). The 1KG data were LD thinned (Plink v2.00a2.3; –-indep-pairwise 50 20 0.5), and 113,571 variants were removed from the merged dataset (retaining 514,708 variants). To test if unidentified strand-flipping between the merged datasets was contributing to the issues (i.e., if strand orientation had not been maintained throughout the plink QC pipeline), snpflip v0.0.6 was run (Biocore NTNU, 2017), but only seven variants were identified as possibly flipped. Next, F_ST_ was calculated (using PLINK v1.90p –-fst flag) for samples putatively representing the same population: UKB Bangladeshi and 1KG Bengali in Bangladesh (BEB), and UKB “White”/British with 1KG GBR. All SNPs across all checks with F_ST_ > 0.15 were subsequently removed, though this only filtered out 144 variants. The PCA was repeated but visualizations of the results demonstrated that this did not resolve the issue.

Next, outliers (samples) were sequentially removed from the 1KG PCA and the PCA was repeated. Because the first several rounds of “outlier identification” kept implicating CEU samples as the culprit, the entire CEU sample was removed from analyses. Subsequently, GBR samples began driving axes of variation across abnormally low PCs. The most obvious irregularity in PCA outputs remained the inflated weighing of European genetic variance relative to global patterns of genetic diversity, therefore, ascertainment bias in the UKB array was suspected.

To test if European genetic variance was being falsely inflated by design of the array, all but the most common genetic variants were removed and the PCA was repeated. Variants were filtered out at 0.1, 0.05, and 0.02 MAF and the results (PCs 1-10) were visualized to assess if excluding rare variants recovered expected patterns of global genetic diversity. Of these three thresholds visualized, only the most stringent (removing SNPs with MAF < 0.1) replicated expected patterns in global genetic diversity in which European variance isn’t artificially inflated. Therefore, this intersection of 199,495 shared SNPs in both the UKB and 1KG arrays were utilized for the PCA (representing variants with MAF > 0.1 and < 0.99). To additionally assess if rare variants were having undue influence on PC weights, MAF was plotted against the absolute PC weight, for PCs 1-10 (Fig. S1.2). Visual assessment indicated this was true for six of the first 10 PCs.

### Supervised genetic ancestry assignment

An SVM was then developed to quantitatively integrate information from PCs 1-15 to identify the most genetically similar reference group(s) to WA and AOA participants, using R package “e1071” [70]. The SVM was trained on a reduced sample of UKB Bangladeshi (*n* = 208) and UKB Pakistani (*n* = 1,484) participants, all 1KG Asian populations, a combined 1KG African group (AFR) (all African populations pooled; *n* = 474), and a combined 1KG European group (EUR) (all European populations pooled; *n* = 394). See SI Section 1.3.2 for further details. The SVM was then used to classify the most probable population affiliation for each AOA and WA participant. SVM classifications were compared to the participants’ region of origin (UKB data field 20115) as an applied assessment of general accuracy (e.g., if AOA or WA participants classified as STU report being from Sri Lanka). Because the training population options were limited, each AOA and WA participant was also assigned to a metapopulation based on the sum of classification probabilities across all training populations comprising a subcontinental region. AOA and WA participants with a South Asian metapopulation summed probability score ≥ 0.7 were subsequently added to the UKB South Asian group, formerly only composed of Bangladeshi-, Indian-, and Pakistani-identifying (BIP) participants.

Genetic affinities of the newly formed UKB South Asian group were assessed further by comparing SVM classifications and reference populations via ADMIXTURE analyses. The same 1KG data as utilized in the PCA were subjected to unsupervised ADMIXTURE analyses (run once for *k* = 3 – 9) and were used to guide the supervised analyses of the UKB data. The results were visualized in *pong* to determine if the SVM classifications reasonably aligned with the reference populations [71, 72].

### Environmental covariates

To maximize the retention of participants incorporated in the GWAS, environmental covariates were imputed using classification and regression trees (CART) within the R statistical package MICE (Multiple Imputation by Chained Equations), resulting in five completed datasets [73, 74] (Figure S2.3). Missingness of each covariate prior to imputation is reported in Table 2 (refer to SI Section 2.2 for further details regarding the environmental covariate imputation process).

After receiving the initial 283 environmental variables requested, the data were placed into broad categories (Figure S2.2 B). Within each domain, the etiology of each variable, and its relationships with all other variables and height were considered (Figure S2.2 B). Correlations among variables within domains were tested, and data conferring the same or highly correlated information were removed (e.g., “Long-standing illness” and “Overall health rating”). For each pair of categorical variables, we constructed 2×2 contingency tables and estimated their association using odds ratios. For all variable pairs, odds ratios were computed on the log scale, and pairs with |log(OR)| > 2 (i.e., OR < 0.14 or OR > 7.39) were flagged as strongly associated. Data hypothesized or known to have a strong genetic etiology that may be shared with the GWAS phenotype were noted (e.g., sitting height and “comparative body size at age 10”) following the guidance for covariate selection outlined in Aschard et al. [58]. Variables of this nature are not recommended for use as environmental covariates in the GWAS but were retained to maximize imputation performance (described below). The data were then assessed for missingness and all variables falling above 20% missingness were discarded prior to any further analyses.

Variables that passed the missingness threshold were then subjected to a multiple imputation (MI) procedure, in order to retain the largest sample of individuals with both genotype and standing height data. MI has been demonstrated to perform as well as or better than the methods typically employed for dealing with uncertainty in environmental data within genetic studies [75,76,77]. Additionally, utilizing MI provides an estimate of the uncertainty produced by using imputed values in downstream analyses, which more transparently reflects the final results [78,79,80].

Several of these variables (uniformly, within each of the five completed datasets) were then recoded for simplicity and to better represent the variable of interest, as it pertains to skeletal health. For example, a dietary restriction variable (UKB data field 6144) originally allowed for six categorical responses describing which food products the participant “never eats”, with “dairy products” as an available option. Of all possible dietary data fields with dairy consumption information, this data field had the least missingness (∼2%) within our study population. Therefore, this variable was recoded as a binary response, such that all responses that selected “dairy products” were recoded as “excludes dairy” and all others were recoded as “does not exclude dairy”.

Subsequent to imputation, nine variables were selected for inclusion as environmental covariates for the GWAS (Table 2). To test if these specific environmental variables are significantly associated with height, and to identify the variance in height explained by additive environmental components alone, an ordinary least squares (OLS) regression model was assessed. The full UKB South Asian sample with un-imputed data (*n* = 7,331) was used for this assessment, using the base R statistical framework (version 4.2.2).

Data recoding scripts are available at https://github.com/ccatram/AnthroGen.

### Imputation of genetic data

Imputed genetic data were utilized for the GWASs. Although the UKB provides imputed data, derived from a combined British and global reference panel (UK 10K and 1000 Genomes), imputation accuracy has not been tested for non-European participants (See Bycroft et al., 2018 [14], UKB Resource 531[https://biobank.ndph.ox.ac.uk/ukb/refer.cgi?id=531], UKB Category 100319 [https://biobank.ndph.ox.ac.uk/showcase/label.cgi?id=100319], and Marchini et al., 2016 [81] for details on the UKB imputation procedures for data versions 1-3). Performance was only tested for imputing the UKB array variants on a sample of 10 Europeans, using data inferred from the British-specific reference panel, alone. Additionally, the info score provided in the imputed data files reflects the average info score across the entire UKB sample, which is predominantly comprised of British, Irish, or “white” identifying participants. Therefore, the provided info score realistically only reflects the certainty in predicting genotypes in Europeans and is of low utility for all other participants. A different imputation was deemed necessary for our sample population (UKB whole genome sequences were not yet released), and the Genome Asia 100K Project (GAsP) was selected for use as the reference panel as it reflected the most appropriate collection of samples publicly available at the time of imputation (2023) [82]. At that time, there were 489 South Asian participants within the full reference panel used by the UKB and 724 within GAsP (phase 1 data).

To prepare the UKB genotype data for imputation, the full array (*n* = 692,466) for the UKB South Asian sample (*n* = 8,178) was QC-ed using Plink v2.00a2.3 (--geno 0.10, –-hwe 0.00001, –-maf 0.00006, –-mind 0.90) and the remaining variants (*n* = 690,869) were converted to vcf format (--recode vcf). Bcftools v1.19 [83] was used to orient the strands to the hg19 reference, split the data by chromosome, and sort by genomic position. This data were then used as the input for imputation via the Michigan Imputation Server (GAsP reference panel, pipeline version 1.7.3, using minimac 4-1.0.2 for imputation, eagle-2.4 for phasing, and *R*^2^ filter of 0) [84]. SNPs with info scores below 0.8 were removed from the returned imputed data (following the convention of Neale Lab QC for UK Biobank GWAS [85]), leaving 6,946,575 remaining variants to be used for the GWAS.

### Height GWAS models

To assess how environmental covariate inclusion may affect GWAS results, two GWAS models were run on a subset of the sample (*n* = 6,954; 1,000 randomly chosen participants were removed for PGS development) and their results compared. GWAS_null_ includes sex, year of birth (YOB), a genetic relatedness matrix (GRM), and the first 15 genetic PCs as covariates in the model, mirroring the typical covariates included in other published height GWAS. GWAS_env_ includes all variables used in GWAS_null_ plus seven additional covariates encompassing socioeconomic, dietary, and demographic influences (Table 2).

The GWASs were run using GCTA v1.93 MLMA-LOCO (--mlma-loco), a powerful mixed linear model based association analysis [86]. The genetic relatedness matrix (GRM) used in the GWAS models was derived from the UKB genotype array data, QC-ed using Plink v2.00a4 (--geno 0.05, –-maf 0.1, –-max-maf 0.99; *n* = 251,556 remaining variants) and calculated with GCTA. The GRM was then used as input for the GCTA PCA (--pca 20) that generated the 15 PCs included in the GWAS models. Although 20 PCs were calculated, visual inspection suggested that only PCs 1-15 provided relevant information regarding population substructure.

GWAS_env_ was run once for each environmental covariate imputed dataset (i.e., five GWAS were run) and the results were pooled following Rubin’s recombining rules, providing pooled effect sizes, standard errors, and p-values for each variant [79] (Figure S2.3).

The effects of environmental covariate inclusion in the GWAS models were evaluated in two primary manners: comparing the number of significant associations and assessing changes in effects for significant associations. To assess the effects of environmental variables on our ability to detect significant genetic associations with height, the number of variants meeting several p-value thresholds were calculated for both GWAS outputs and these absolute values were compared. To characterize differences in SNP effects between GWAS results, three approaches were taken. First, variants with *p* < 10^-6^ output by GWAS_env_ were identified and the GWAS_env_ beta values (β_env_) were subtracted from the GWAS_null_ beta values (β_null_) to ascertain the relative change in effect sizes from the “default” height GWAS model. Secondly, for this same sample of SNPs, the absolute differences and percent differences between β_null_ and β_env_ were calculated. Thirdly, to qualitatively assess discrepancies in β distributions, density plots were visualized, both genome-wide and within chromosomes.

Lastly, FUMA-GWAS [86] was used to summarize functional annotations of the GWAS_env_ output and assess overlap with SNP associations currently published in the GWAS catalog [88]. Input parameters and results of the analyses can be found in SI Section 2.3.1.

### Polygenic score (PGS) assessments and comparisons

Height PGS models derived from each GWAS’s summary statistics were developed using LDpred2-auto within the ‘bigsnpr’ R package [30, 31]. Only SNPs also found within HapMap3 were incorporated in the PGS (*n* = 896,160), and all default values recommended for the ‘snp_ldpred2_auto()’ function were used for the analysis (following Privé’s 2023 vignette, which can be found here: https://privefl.github.io/bigsnpr/articles/LDpred2.html [30]). LDpred2-auto was chosen for our PGS development as it infers SNP heritability (*h^2^*) and polygenicity from the LDpred model without necessitating a validation set for parameter-tuning, allowing us to retain a larger sample size than would be possible otherwise [30].

Performance was tested by obtaining *R*^2^ between PGS scores and height for UKB South Asian participants removed from the sample prior to GWAS analyses (*n* = 997; 1,000 participants were randomly selected, but three were later removed due to data missingness). Sex biases in PGS performance were additionally tested in the same manner, but within sex-stratified samples.

The utility of our PGS equations were then tested by assessing their performance relative to the South Asian PGS equation created by Yengo et al., 2022 [34], which was developed from a GWAS meta-analysis (*n* = 77,890) and tested on UKB South Asian participants (*n* = 9,257). The associated PGS scoring file (PGS002800) was downloaded from the PGS Catalog [33] and applied to our PGS test sample using ESCALATOR v.2.0.0 (Lin and Fisher, 2022 [36]; [https://github.com/menglin44/ESCALATOR]), a PGS harmonization pipeline.

PGS002800 performance was assessed for two subsets of SNPs: PGS_Y1_: the subset of SNPs found in our GWAS (and passing ESCALATOR quality control filtering) (*n* = 898,486), and PGS_Y2_: the subset of variants that were available within our imputed data (*n* = 1,032,672). The Yengo et al. South Asian PGS makes use of 1,156,741 variants in its entirety, but we were unable to include the complete set in our analyses due to poor imputation quality for 12% of SNPs [34].

Testing for significant differences across PGS model performance (both between our PGS models and Yengo-derived PGS models) was conducted using R packages *cocor* and *r2redux* [32,33]. For PGS-only comparisons (i.e., those that assess PGS *R^2^* derived from regressing height on PGS scores only), *cocor* was used. For significance testing across PGS *R^2^* derived from regressions incorporating additional covariates, *r2redux* was used.

Additionally, to test if PGS002800 performance may be inflated due to unadjusted environmental influences (as opposed to the alternative hypothesis that higher relative performance is due to a better-powered training sample), *R*^2^ between PGS scores and the model covariates were calculated for each combination of the data subsets referenced above (Table S2.15). While we expect correlations between PGS scores and environmental variables to be very low, we predict that if the effects of collinearity impacted the development of PGS002800 β weights then the resulting PGS scores will exhibit higher *R*^2^ values compared to the PGS scores derived from our environmentally adjusted GWAS data. Further investigations into the influence of environmental adjustments on PGS development can be found in SI Section 2.4.2.

