## Supplemental Information for "Improving GWAS performance in underrepresented groups by appropriate modeling of genetics, environment, and sociocultural factors"

**Affiliations:**
[1]Department of Anthropology, University of California Davis, Davis, CA, 95616, USA
[2]Department of Population and Public Health Sciences, Center for Genetic Epidemiology, Keck School of Medicine, University of Southern California, CA 91001, USA
[3]Department of Biomedical Informatics, University of Colorado Anschutz Medical Campus, Aurora, CO 80045, USA
[4]Colorado Center for Personalized Medicine, University of Colorado Anschutz Medical Campus, Aurora, CO 80045, USA
[5]UC Davis Genome Center, University of California Davis, Davis, CA, 95616, USA

**SUPPLEMENTAL INFORMATION**

Section 1: Population genetics
- 1.1: Sample demographics
- 1.2: Genetic data and quality control
  - 1.2.1: 1000G reference data
  - 1.2.2: UKB genotype array QC
  - 1.2.3: Genotype array ascertainment bias assessments
- 1.3: Genetic affinities of UKB participants
  - 1.3.1: PCA
  - 1.3.2: SVM
  - 1.3.3: ADMIXTURE

Section 2: GWAS and PGS
- 2.1: UKB South Asian height data
- 2.2: Environmental covariates
  - 2.2.1: Methods
  - 2.2.2: Extended results
- 2.3: GWAS extended results
  - 2.3.1: Functional annotation
- 2.4: PGS
  - 2.4.1: Methods
  - 2.4.2: Extended results
- 2.5: Extended discussion

References

**SECTION 1: Population genetics**
**1.1: Sample demographics**

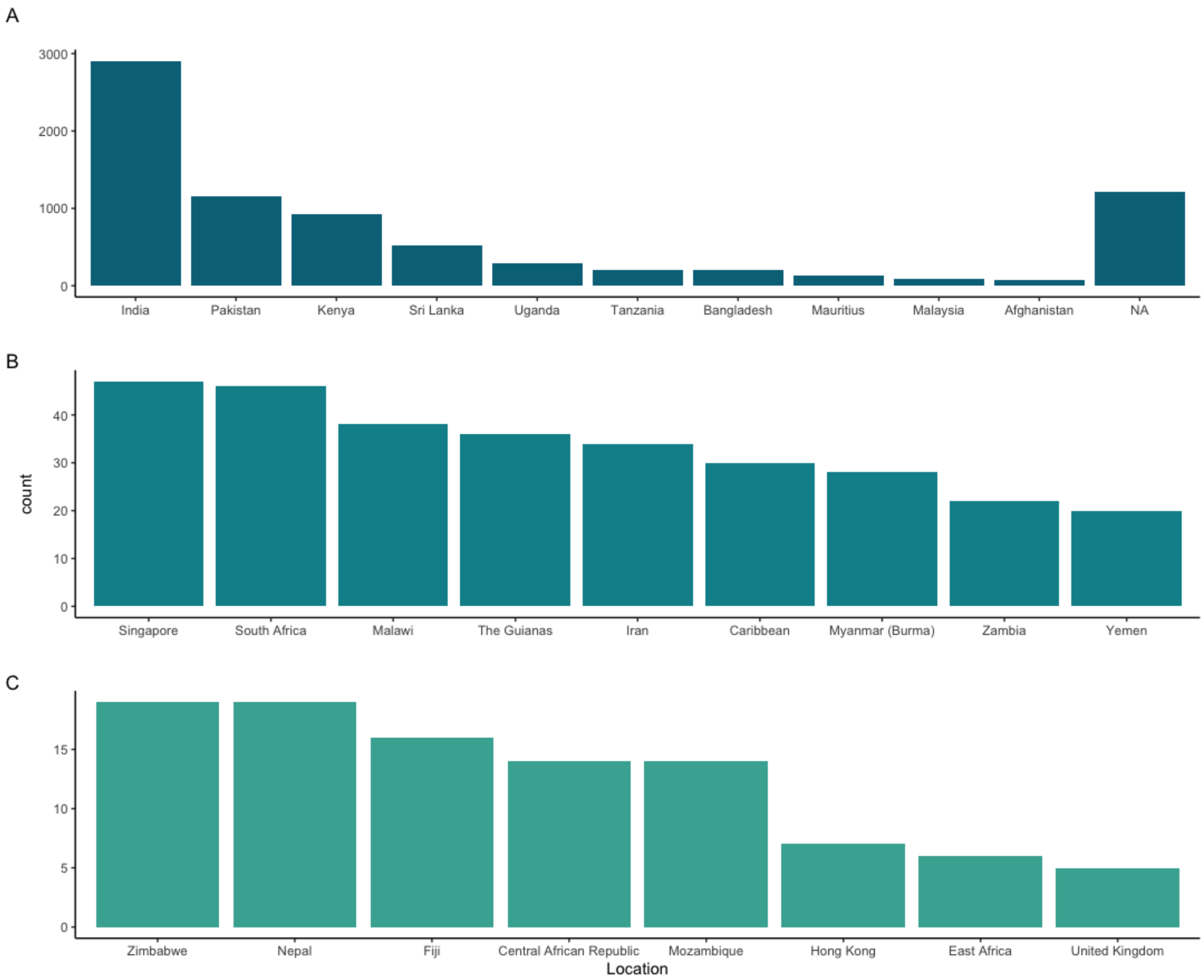

**Figure S1.1**. Country or region of origin (UKB Data ID 20115) of UKB South Asian participants. A) Locations with 50 or more participants; B) locations with 20 - 49 participants; C) locations with 5 - 19 participants. Locations with fewer than 5 participants are omitted.

**Table S1.1.** Country or region of origin (UKB Data Field 20115) of self-identified Bangladeshi participants (UKB Data Field 21000) in the UKB South Asian sample (*n* = 212).

| Bangladesh | India | Missing (NA) |
|---|---|---|
| 191 (90.1%) | 1 (0.005%) | 20 (0.09%) |

**Table S1.2.** Country or region of origin (UKB Data ID 20115) of self-identified Indian participants (UKB Data Field 21000) in the UKB South Asian sample (*n* = 5,006). Countries or regions with fewer than 5 responses are omitted.

| Caribbean | Central African Republic | Fiji | India | Kenya | Malawi | Malaysia |
|---|---|---|---|---|---|---|
| 20 (0.4%) | 11 (0.22%) | 14 (0.28%) | 2722 (54.4%) | 802 (16.02%) | 35 (0.7%) | 58 (1.16%) |
| **Mauritius** | **Myanmar (Burma)** | **Mozambique** | **Pakistan** | **Singapore** | **South Africa** | **Sri Lanka** |
| 47 (0.94%) | 7 (0.14%) | 11 (0.22%) | 35 (0.7%) | 40 (0.8%) | 37 (0.74%) | 21 (0.42%) |
| **Tanzania** | **The Guianas** | **Uganda** | **Yemen** | **Zambia** | **Zimbabwe** | **Missing (NA)** |
| 183 (3.66 %) | 19 (0.38%) | 262 (5.23%) | 17 (0.34%) | 21 (0.42%) | 15 (0.3%) | 574 (11.47%) |

**Table S1.3.** Country or region of origin (UKB Data ID 20115) of self-identified Pakistani participants (UKB Data Field 21000) in the UKB South Asian sample (*n* = 1,528). Countries or regions with fewer than 5 responses are omitted.

| India | Kenya | Pakistan | Tanzania | Uganda | Missing (NA) |
|---|---|---|---|---|---|
| 99 (6.48%) | 54 (3.53%) | 1093 (71.53%) | 6 (0.39%) | 12 (0.79%) | 240 (15.71%) |

**Table S1.4.** Country or region of origin (UKB Data ID 20115) of self-identified 'Any other Asian' participants (UKB Data Field 21000) in the UKB South Asian sample (*n* = 1,045). Countries or regions with fewer than 5 responses are omitted.

| Afghanistan | Caribbean | India | Iran | Kenya | Malaysia |
|---|---|---|---|---|---|
| 72 (6.9%) | 9 (0.86%) | 17 (1.63%) | 33 (3.16%) | 59 (5.65%) | 31 (2.97%) |
| **Mauritius** | **Myanmar (Burma)** | **Nepal** | **Pakistan** | **Singapore** | **South Africa** |
| 85 (8.13%) | 11 (1.05%) | 19 (1.82%) | 13 (1.24%) | 7 (0.67%) | 7 (0.67%) |
| **Sri Lanka** | **Tanzania** | **The Guianas** | **Uganda** | **Missing (NA)** | |
| 500 (47.85%) | 19 (1.82%) | 12 (1.15%) | 22 (2.11%) | 106 (10.14%) | |

**Table S1.5.** Country or region of origin (UKB Data ID 20115) of self-identified ‘White and Asian’ participants (UKB Data Field 21000) in the UKB South Asian sample ($n$ = 387). Countries or regions with fewer than 5 responses are omitted.

| India | Myanmar (Burma) | Pakistan | The Guianas | Missing (NA) |
|---|---|---|---|---|
| 65 (16.8%) | 7 (1.81%) | 12 (3.1%) | 5 (1.29%) | 263 (67.96%) |

### 1.2: Genetic data and quality control

#### 1.2.1: 1000 Genomes (1KG) reference data

**Table S1.6**. 1KG Phase 3 reference data used in analyses, after removing 1st and 2nd degree relatives.

| Code | Sample* | *n* |
|---|---|---|
| ESN | Esan in Nigeria | 94 |
| GWD | Gambian in Western Division, Mandinka | 109 |
| LWK | Luhya in Webuye, Kenya | 85 |
| MSL | Mende in Sierra Leone | 78 |
| YRI | Yoruba in Ibadan, Nigeria | 108 |
| CEU | Utah residents (CEPH) with Northern and Western European ancestry | 94 |
| GBR | British in England and Scotland | 88 |
| IBS | Iberian Populations in Spain | 107 |
| TSI | Toscani in Italy | 105 |
| BEB | Bengali in Bangladesh | 85 |
| GIH | Gujarati Indians in Houston, TX, USA | 100 |
| ITU | Indian Telugu in the UK | 101 |
| PJL | Punjabi in Lahore, Pakistan | 90 |
| STU | Sri Lankan Tamil in the UK | 99 |
| CDX | Chinese Dai in Xishuangbanna, China | 89 |
| CHB | Han Chinese in Beijing, China | 103 |
| CHS | Southern Han Chinese | 101 |
| JPT | Japanese in Tokyo, Japan | 104 |
| KHV | Kinh in Ho Chi Minh City, Vietnam | 99 |
| **Total** | | **1839** |

* Descriptions obtained from 1000 Genomes Project Consortium (2015) and Casillas et al. (2018) [1, 2].

#### 1.2.2: UKB genotype array quality control (QC)

**Table S1.7**. QC procedure for UKB array. Input/output *n* refers to the number of variants or samples, depending on the QC step.

| Input (*n*) | Filter(s) | Software | Output (*n*) |
|---|---|---|---|
| 784,256 SNPs | --geno 0.10<br>--hwe 0.000001<br>--maf 0.000097<br>--mind 0.90 | PLINK v2.00a2.3 | 695,351 SNPs |
| *23,732 variants removed due to missing genotype data; 13,313 variants removed due to Hardy-Weinberg exact test; 51,860 variants removed due to allele frequency threshold* | | | |
| 10,288 samples | --unrelated<br>--degree 2 | KING 2.1.8 | 8,967 samples |
| *1,321 individuals removed due to relatedness* | | | |
| 695,351 SNPs | --maf 0.0001 | PLINK v2.00a2.3 | 692,466 |
| *2,885 variants removed due to allele frequency threshold* | | | |

Following the QC filtering steps listed above, the QC-ed UKB genotyping array was merged with 1KG data to facilitate analyses comparing UKB data with global references. The full intersection of SNPs (668,051 shared SNPs) yielded unexpected patterns of genetic diversity, prompting further investigation into the quality of the UKB and 1KG data.

#### 1.2.3: Genotype array ascertainment bias assessments

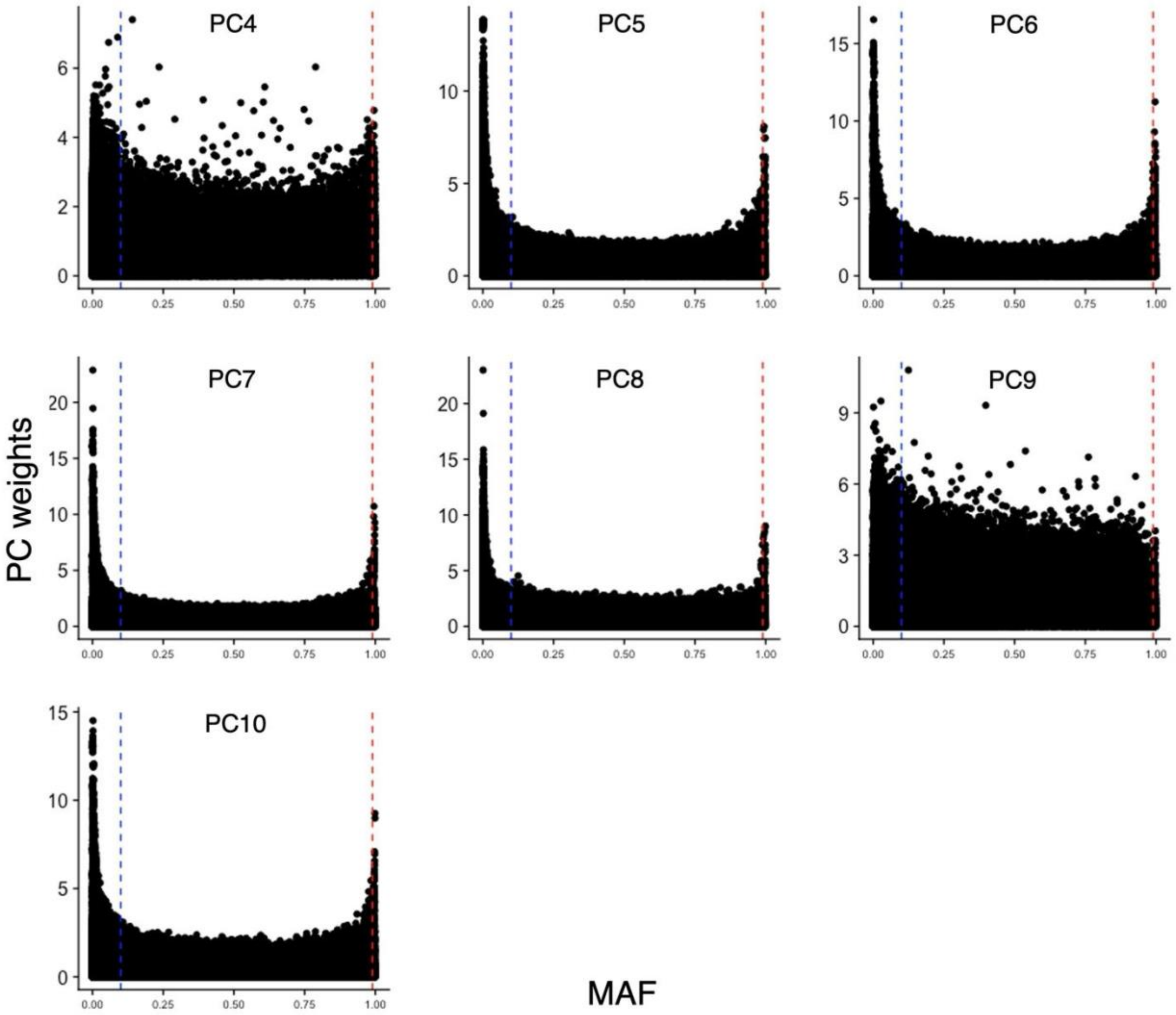

**Figure S1.2.** Absolute PC weights and minor allele frequency (MAF) of the full SNP array intersection of 1KG and UKB data. Alleles with MAF > 0.99 (red dashed line) and < 0.1 MAF (blue dashed line) in 1KG data were removed from both datasets (*n* = 199,495 retained).

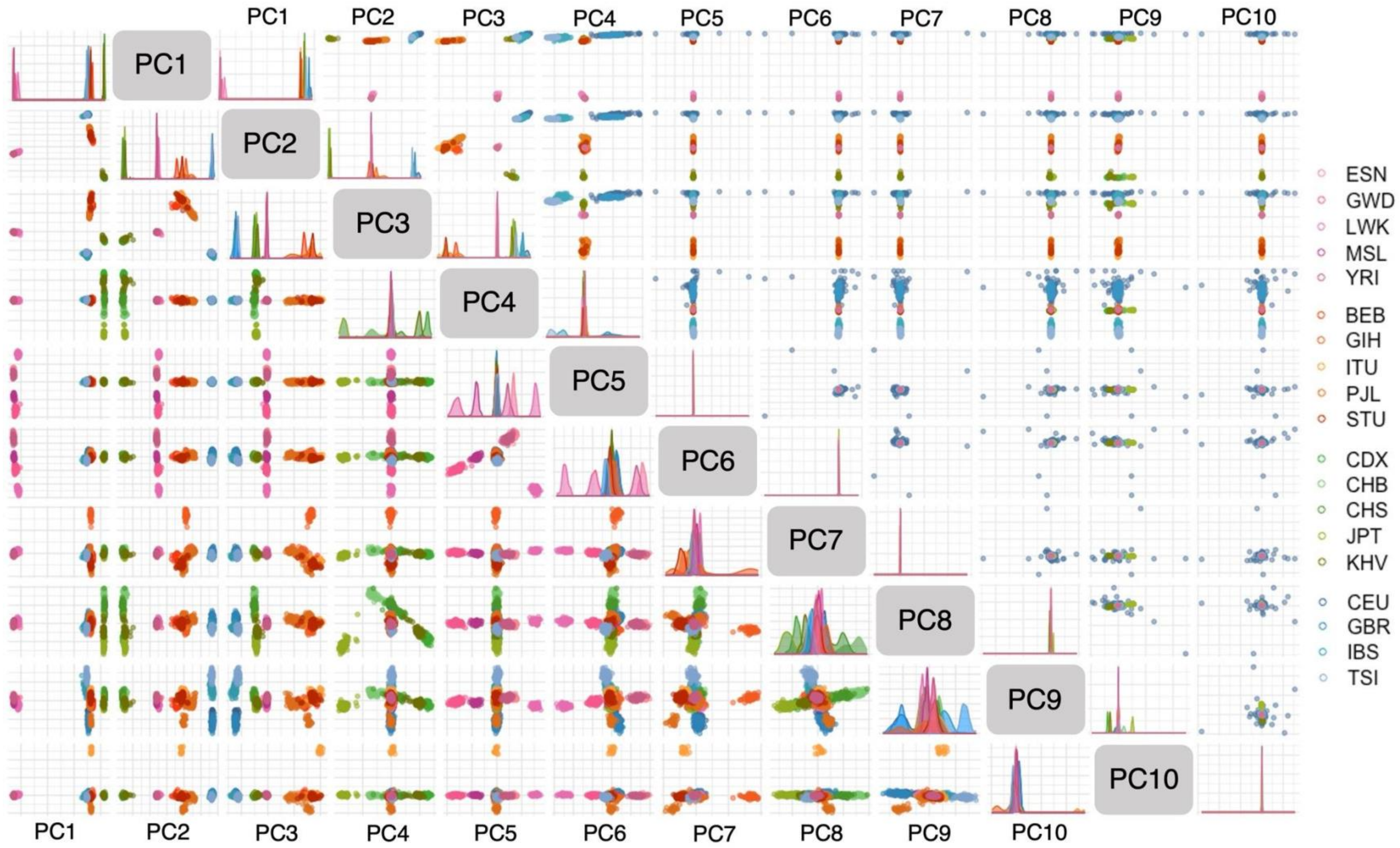

**Figure S1.3.** Comparison of 1KG PCA between the initial SNP intersection with the UKB array (upper) and the SNP intersection filtered to adjust for ascertainment bias (lower). Rows (PCs) are labeled along the diagonal. Column labels are listed on top for the upper triangle, and on the bottom for the lower triangle.

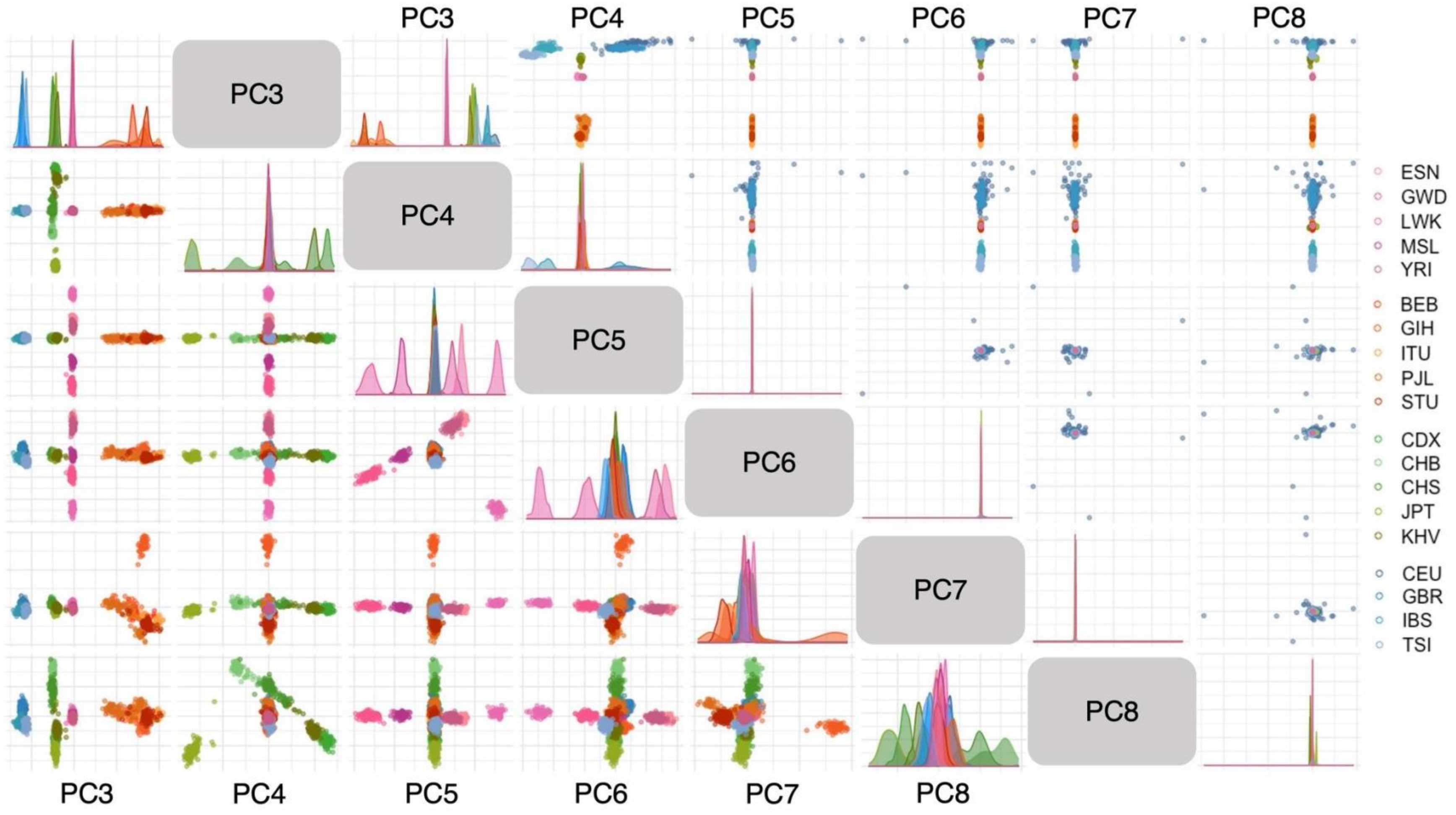

**Figure S1.4.** Enlarged comparison of 1KG PCA between the initial SNP intersection with the UKB array (upper) and the SNP intersection filtered to adjust for ascertainment bias (lower), for PCs 3 - 8. Rows (PC) are labeled along the diagonal. Column labels are listed on top for the upper triangle, and on the bottom for the lower triangle.

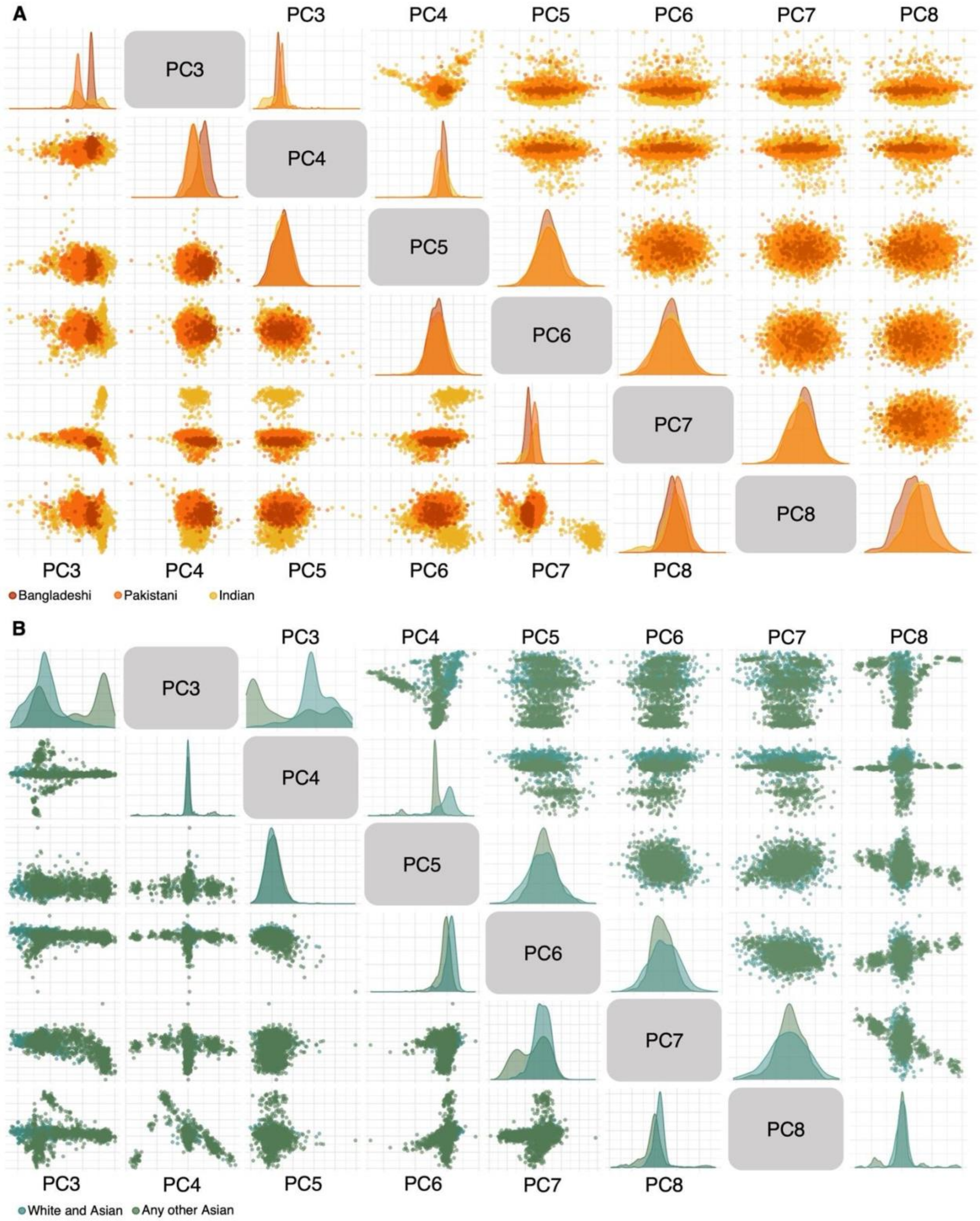

**Figure S1.5.** Comparisons of UKB PCA between the initial SNP intersection with the 1KG array (upper) and the SNP intersection filtered to adjust for ascertainment bias (lower), for PCs 3 - 8. Rows (PC) are labeled along the diagonal. Column labels are listed on top for the upper triangle, and on the bottom for the lower triangle. A) BIP participants; B) AOA and WA participants.

### 1.3: Genetic affinities of UKB participants

#### 1.3.1: PCA

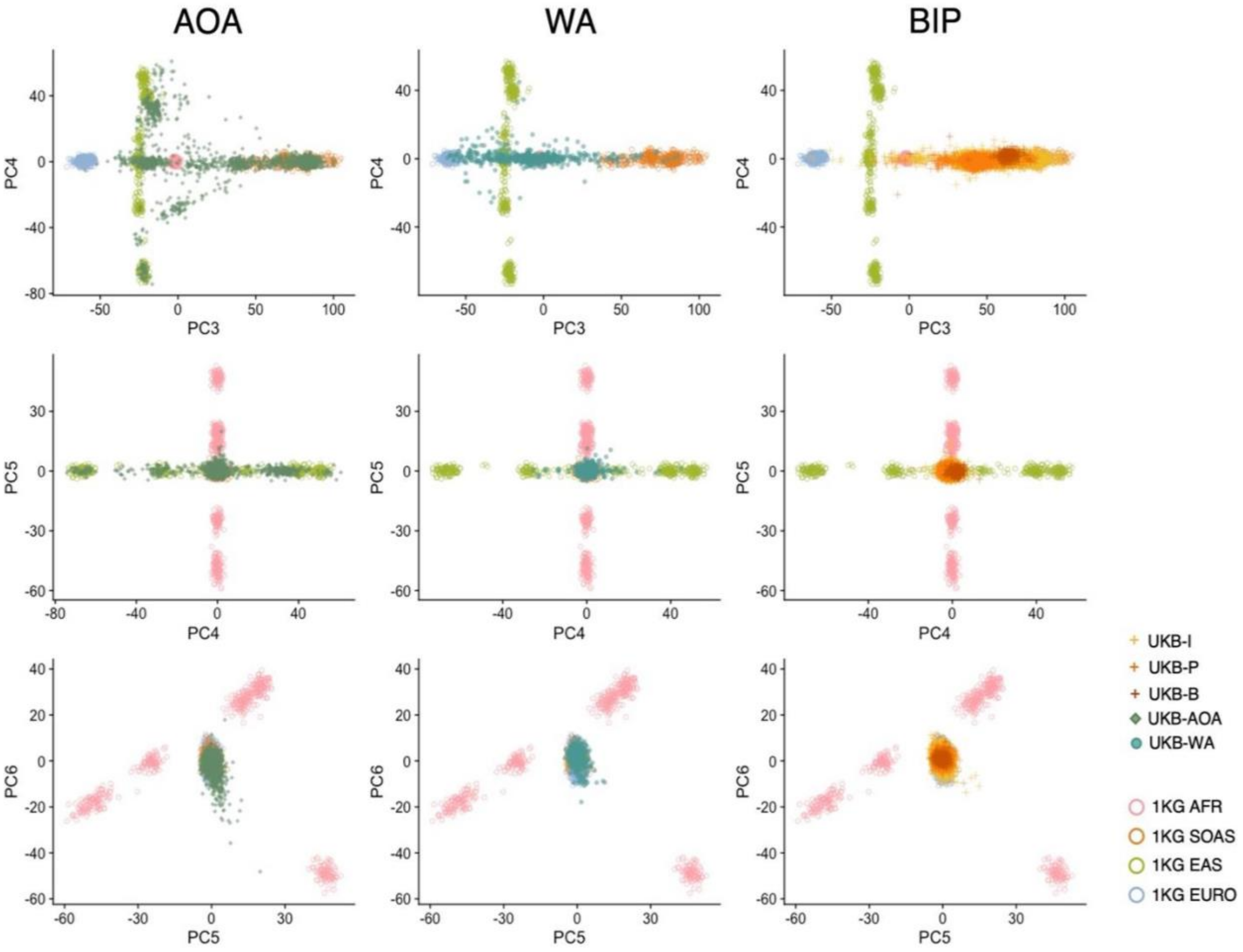

**Figure S1.6.** UKB participants projected onto 1KG PCs 3 - 6.

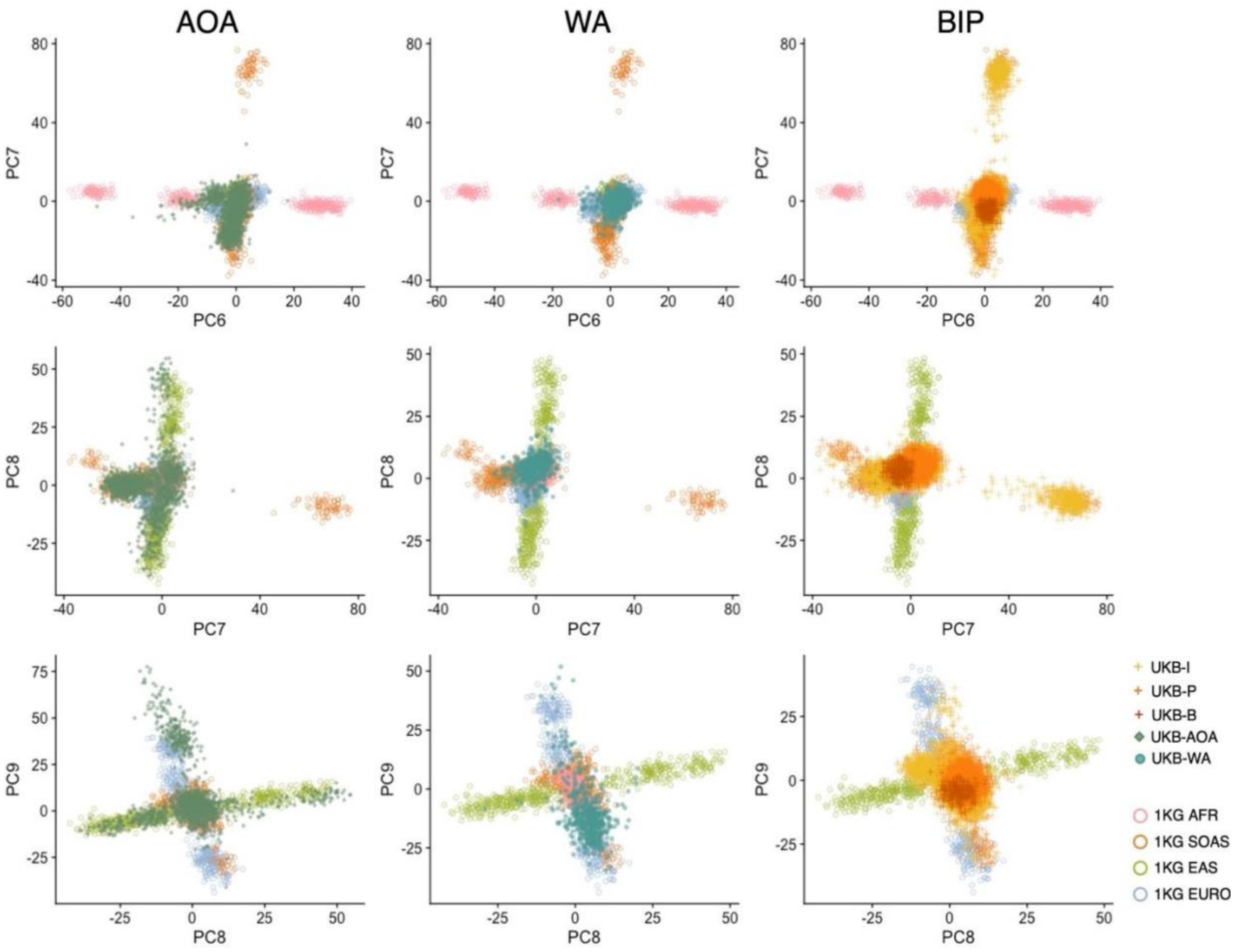

**Figure S1.7.** UKB participants projected onto 1KG PCs 6 - 9.

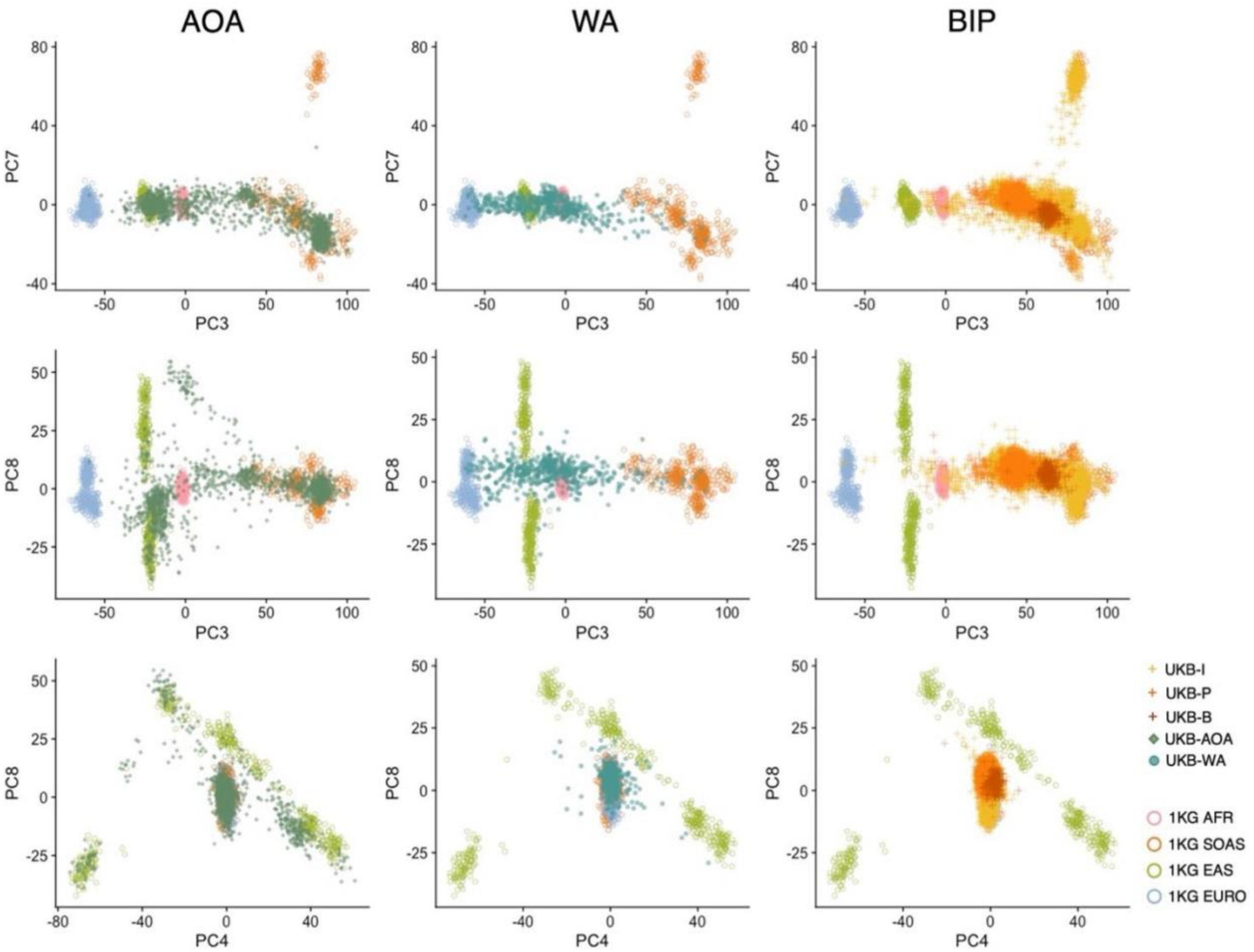

**Figure S1.8.** UKB participants projected onto 1KG PCs 3, 4, 7, and 8.

#### 1.3.2: SVM

Training data used for the SVM included PC1-15 scores from: UKB Bangladeshi (*n* = 208), UKB Pakistani (*n* = 1,484), all 1KG Asian populations, 1KG African samples collapsed into one group (*n* = 474), and 1KG European samples collapsed into one group (*n* = 394). UKB Bangladeshi and Pakistani outliers per PC (mean score +/- 4SD) were removed prior to SVM training. UKB Indian participants were not used to train the SVM in order to simplify the model, as both Bangladeshi and Pakistani genetic variance fall within the full range of Indian PC scores. The SVM model was then tuned to identify best cost and gamma parameters using 10-fold cross validation and was applied to the full training set. A class agreement confusion matrix was created to assess within sample accuracy, and the best-performing model was retained and used to classify WA and AOA participants (kappa = 0.94; corrected rand = 0.93).

Participants were included in the new UKB South Asian sample if the predicted classification was any South Asian group and the probabilities of all South Asian classifications (UKB Bangladeshi, UKB Pakistani, BEB, GIH, ITU, PJL, STU) summed to at least 0.7 (an arbitrarily chosen threshold). The SVM was additionally used to assess the genetic affinities of the UKB Indian participants (in part to assess the model's performance outside of the training data, but also to identify outliers within the UKB Indian sample), resulting in the removal of 10 UKB Indian participants due to a EURO SVM assignment or low South Asian probability sum (under 70%).

**Table S1.8**. SVM classifications of UKB AOA and WA participants per classification probability threshold.

| Probability | UKB Group | AFR | EUR | EAS | | | | | SAS | | | | | | |
|---|---|---|---|---|---|---|---|---|---|---|---|---|---|---|---|
| | | AFR | EUR | CDX | CHB | CHS | KHV | JPT | BEB | GIH | ITU | PJL | STU | UKB-B* | UKB-P* |
| ≥ 0 | AOA | 3 | 165 | 25 | 73 | 16 | 181 | 56 | 38 | 14 | 208 | | 373 | 66 | 397 |
| | WA | | 145 | 1 | 6 | 2 | 5 | | | | 3 | | 8 | 37 | 342 |
| ≥ 0.75 | AOA | 1 | 24 | 14 | 12 | 7 | 149 | 53 | 16 | 1 | 41 | | 249 | 20 | 271 |
| | WA | | 85 | | | | 2 | | | | | | 1 | 6 | 221 |
| ≥ 0.9 | AOA | 1 | 4 | 9 | 1 | | 84 | 43 | 12 | 1 | 6 | | 121 | 7 | 222 |
| | WA | | 51 | | | | | | | | | | | 3 | 101 |

*UKB Bangladeshi and Pakistani classifications are referred to as UKB-B and UKB-P, respectively. Shaded cells received 0 classifications.

**Table S1.9**: Subcontinental assignments of UKB AOA and WA participants derived from summed SVM classification probabilities (sum ≥ 0.7).

| | AFR | EUR | EAS | SAS |
|---|---|---|---|---|
| **AOA** | 1 | 39 | 332 | 1036 |
| **WA** | | 92 | 8 | 345 |

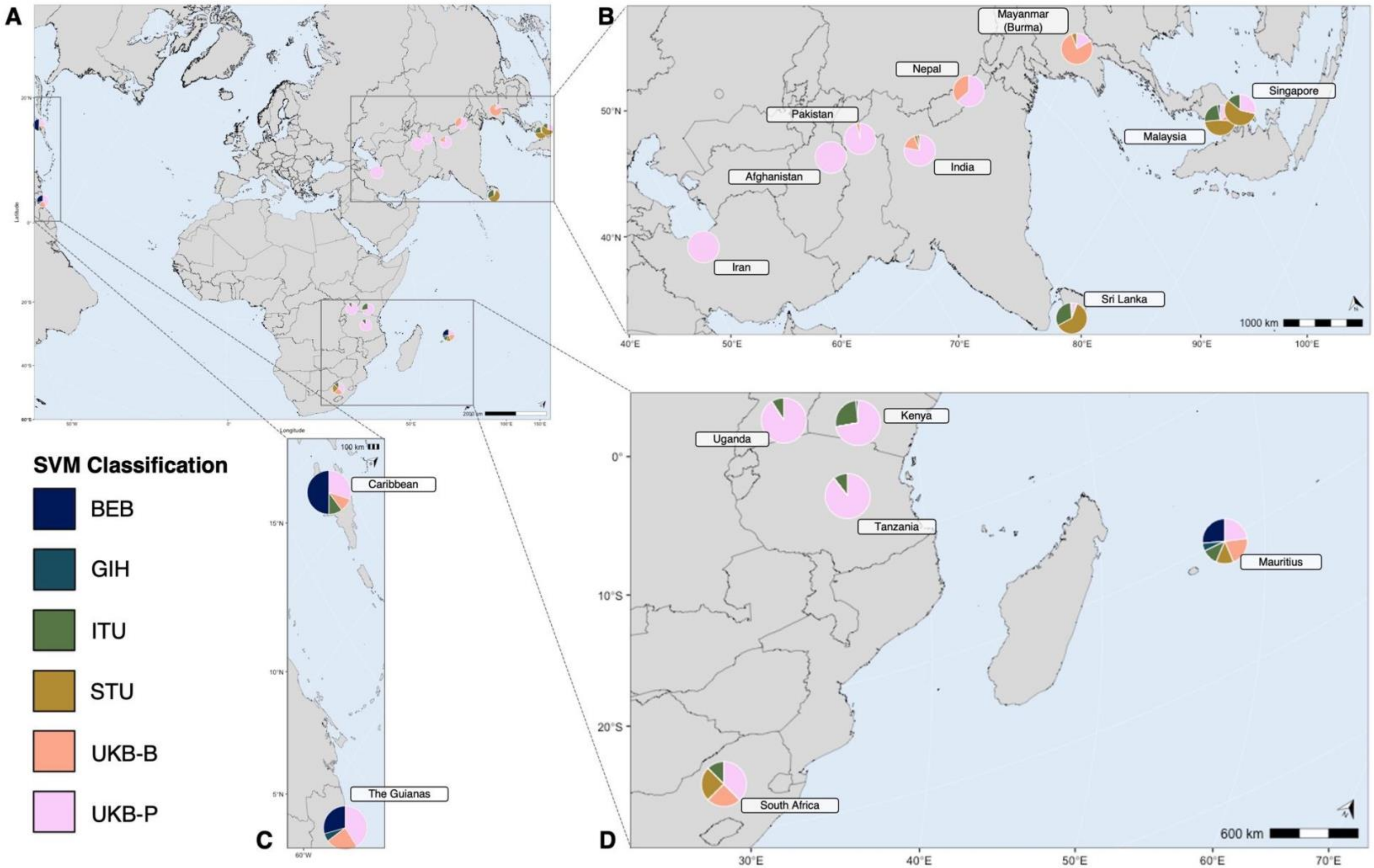

**Figure S1.9.** SVM assignments for AOA and WA participants and their reported region of origin data (UKB data field 20115; *n* = 1,063 [1,011 participants did not provide data]). Only regions with five or more participant responses are plotted. A) All regions of origin; B) Asia (Afghanistan, India, Iran, Malaysia, Myanmar / Burma, Nepal, Pakistan, Singapore, Sri Lanka); C) Caribbean and The Guianas; D) Africa (Kenya, South Africa, Tanzania, Uganda) and Mauritius.

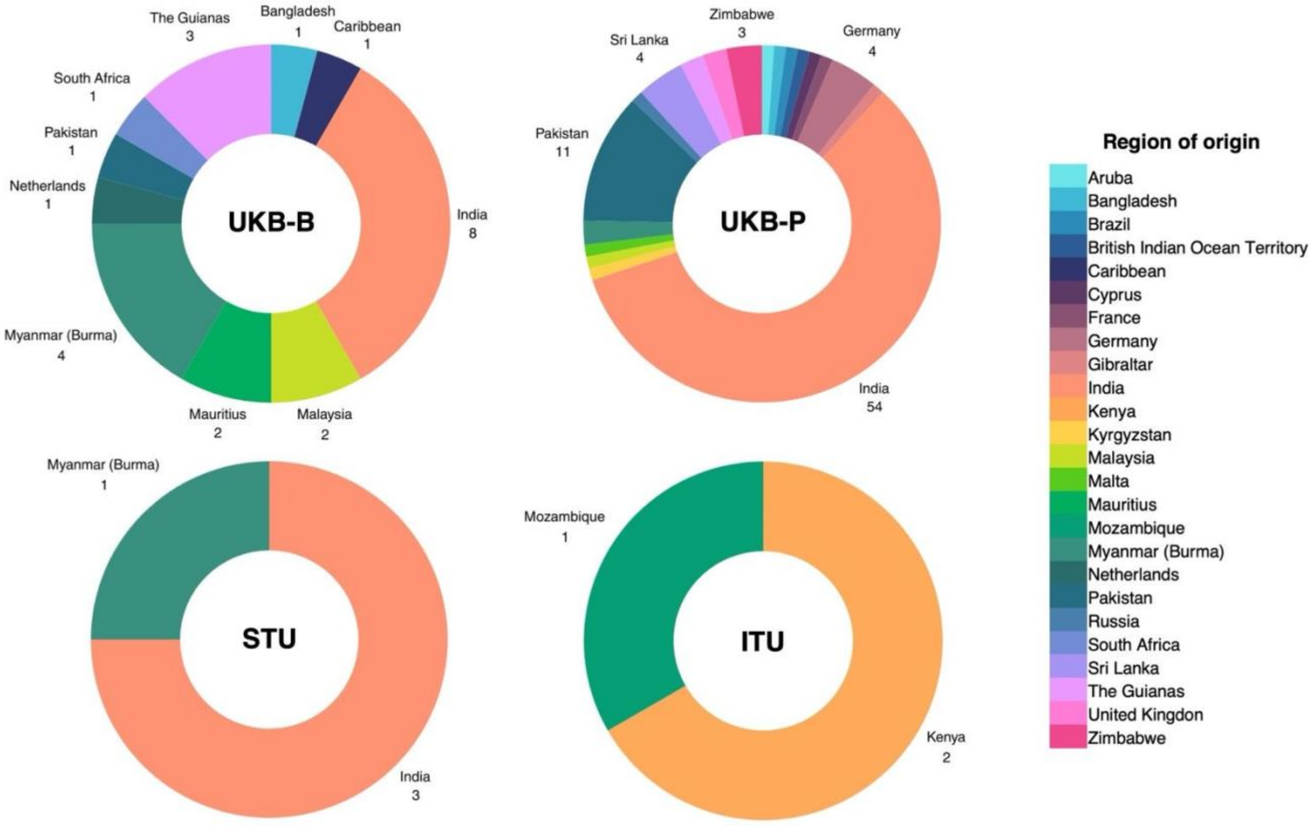

**Figure S1.10.** SVM classifications of WA participants, colored by region of origin (*n* listed beneath region when possible).

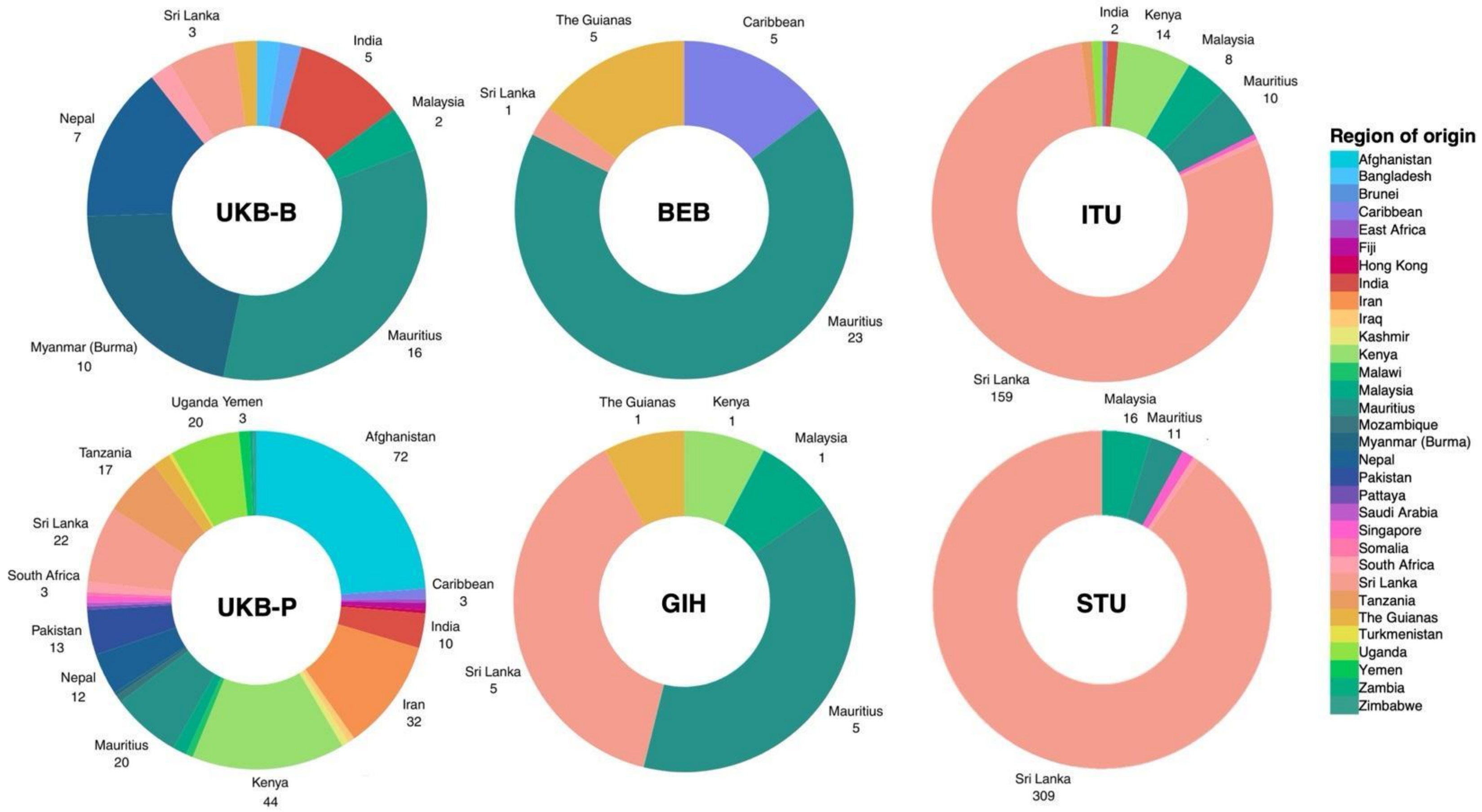

**Figure S1.11.** SVM classifications of AOA participants, colored by region of origin (*n* listed beneath region when possible).

#### 1.3.3: ADMIXTURE

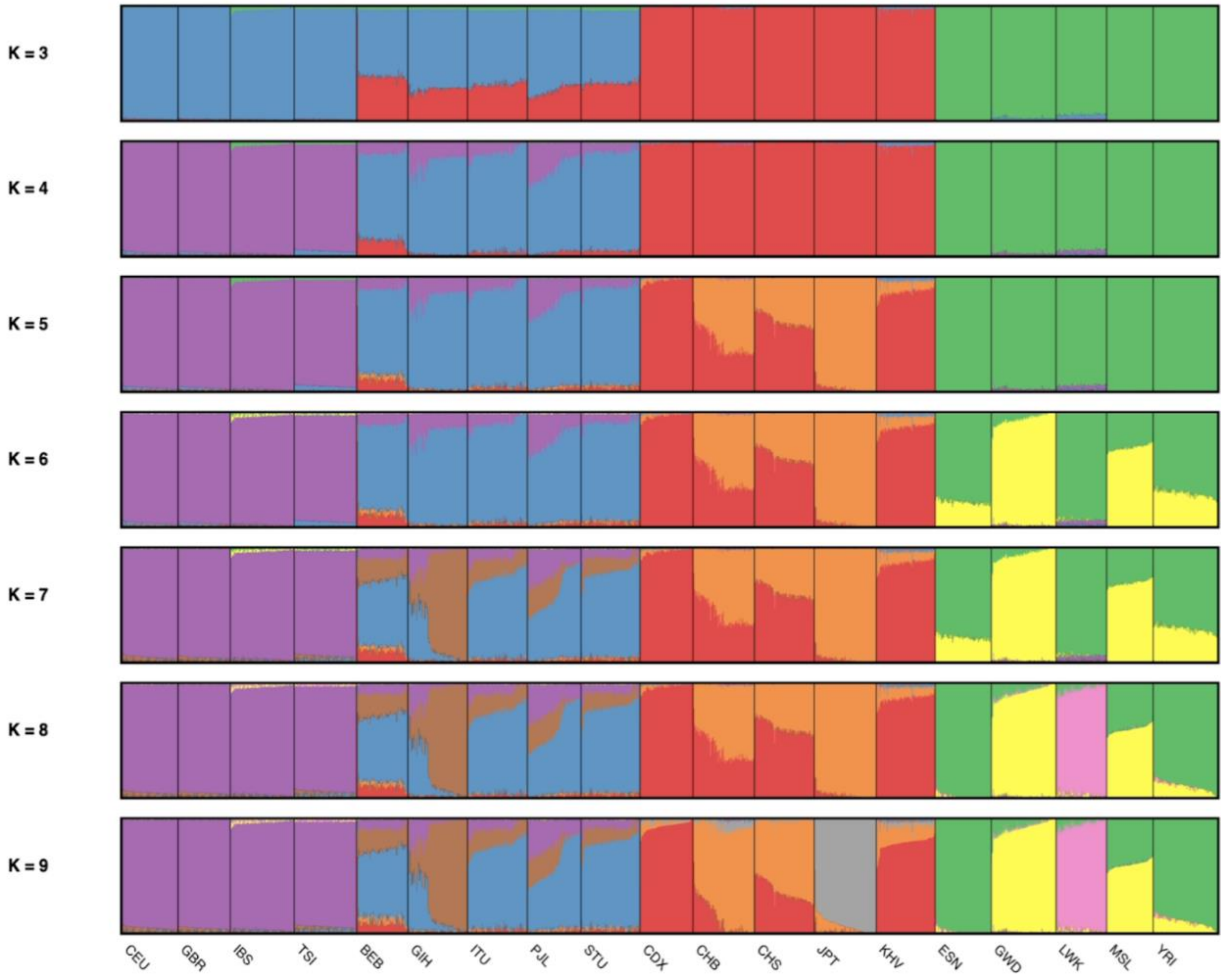

**Figure S1.12.** *pong* visualization of unsupervised ADMIXTURE analyses (for $k$ = 3 - 9) using 1000 Genomes reference data and 199,495 shared, QC-ed SNPs with the UK Biobank array. These data were then used to supervise the ADMIXTURE analyses of the UKB participants.

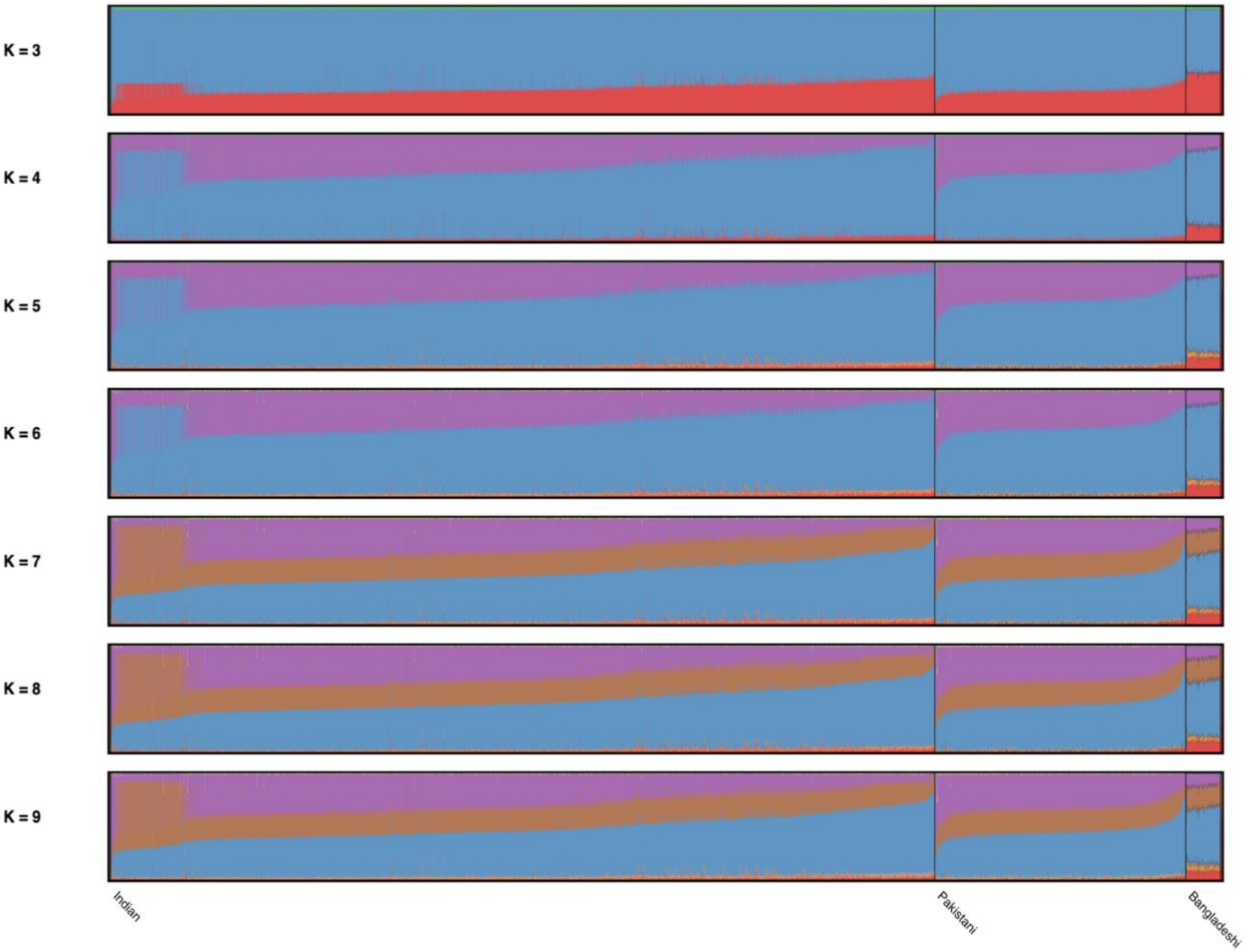

**Figure S1.13**. *pong* visualization of supervised ADMIXTURE analyses (for $k$ = 3 - 9) using 1000 Genomes reference data to infer population affinity proportions of UKB Indian, Pakistani, and Bangladeshi participants. Colors are consistent with Fig. S1.12 and Table S1.10.

**Table S1.10**. Average percent cluster membership for *k* = 3-9, output by the supervised ADMIXTURE analyses for UKB Bangladeshi, Indian, and Pakistani participants.

ADMIXTURE average percent cluster membership

| Group | *k* | | | | | | | | | |
|---|---|---|---|---|---|---|---|---|---|---|
| | | *Europe* | *Asia* | *Africa* | | | | | | |
| Bangladeshi | | 59.4% | 38.3% | 2.3% | | | | | | |
| Indian | **3** | 72.9% | 24.6% | 2.5% | | | | | | |
| Pakistani | | 75.4% | 22.1% | 2.5% | | | | | | |
| | | *Europe* | *South Asia* | *East Asia* | *Africa* | | | | | |
| Bangladeshi | | 13.9% | 70.3% | 15.4% | -- | | | | | |
| Indian | **4** | 28.5% | 67.9% | 3.0% | 0.7% | | | | | |
| Pakistani | | 34.5% | 62.3%% | 2.4% | 0.8% | | | | | |
| | | *Europe* | *South Asia* | *East Asia* | *Japan* | *Africa* | | | | |
| Bangladeshi | | 13.8% | 69.7% | 10.7% | 5.4% | -- | | | | |
| Indian | **5** | 28.3% | 67.3%% | 1.8% | 1.9% | 0.7% | | | | |
| Pakistani | | 34.3%% | 61.8%% | 1.2% | 2.0% | 0.8% | | | | |
| | | *Europe* | *South Asia* | *East Asia* | *Japan* | *Africa - GWD* | *Africa - LWK* | | | |
| Bangladeshi | | 13.8% | 69.7% | 10.7% | 5.4% | -- | -- | | | |
| Indian | **6** | 28.2% | 67.3% | 1.8% | 1.9% | -- | -- | | | |
| Pakistani | | 34.2% | 61.8%% | 1.1% | 1.9% | -- | 0.5% | | | |
| | | *Europe* | *South Asia* | *South Asia - GIH* | *East Asia* | *Japan* | *Africa - GWD* | *Africa - LWK* | | |
| Bangladeshi | | 11.9% | 51.3% | 21.0% | 10.2% | 5.2% | -- | -- | | |
| Indian | **7** | 25.7% | 45.3% | 25.3% | 1.4% | 1.6% | -- | -- | | |
| Pakistani | | 32.0% | 42.6% | 22.1% | 0.9% | 1.7% | -- | 0.5% | | |
| | | *Europe* | *South Asia* | *South Asia - GIH* | *East Asia* | *Japan* | *Africa - GWD* | *Africa - LWK* | *Africa - ESN* | |
| Bangladeshi | | 11.9% | 51.4% | 20.9% | 10.2% | 5.2% | -- | -- | -- | |
| Indian | **8** | 25.6% | 45.4% | 25.2% | 1.4% | 1.6% | -- | -- | -- | |
| Pakistani | | 31.9% | 42.6% | 22.0% | 0.9% | 1.6% | -- | -- | -- | |
| | | *Europe* | *South Asia* | *South Asia - GIH* | *East Asia - China* | *Southeast Asia* | *Japan* | *Africa - GWD* | *Africa - LWK* | *Africa - ESN* |
| Bangladeshi | | 11.8% | 51.1% | 20.3% | 5.7% | 8.3% | 2.4% | -- | -- | -- |
| Indian | **9** | 25.5% | 45.1% | 24.7% | 1.8% | 1.1% | 1.1% | -- | -- | -- |
| Pakistani | | 31.8% | 42.4% | 21.5% | 2.1% | 0.8% | 0.5% | -- | -- | -- |

Clusters are colored by the reference groups most representative of that cluster.

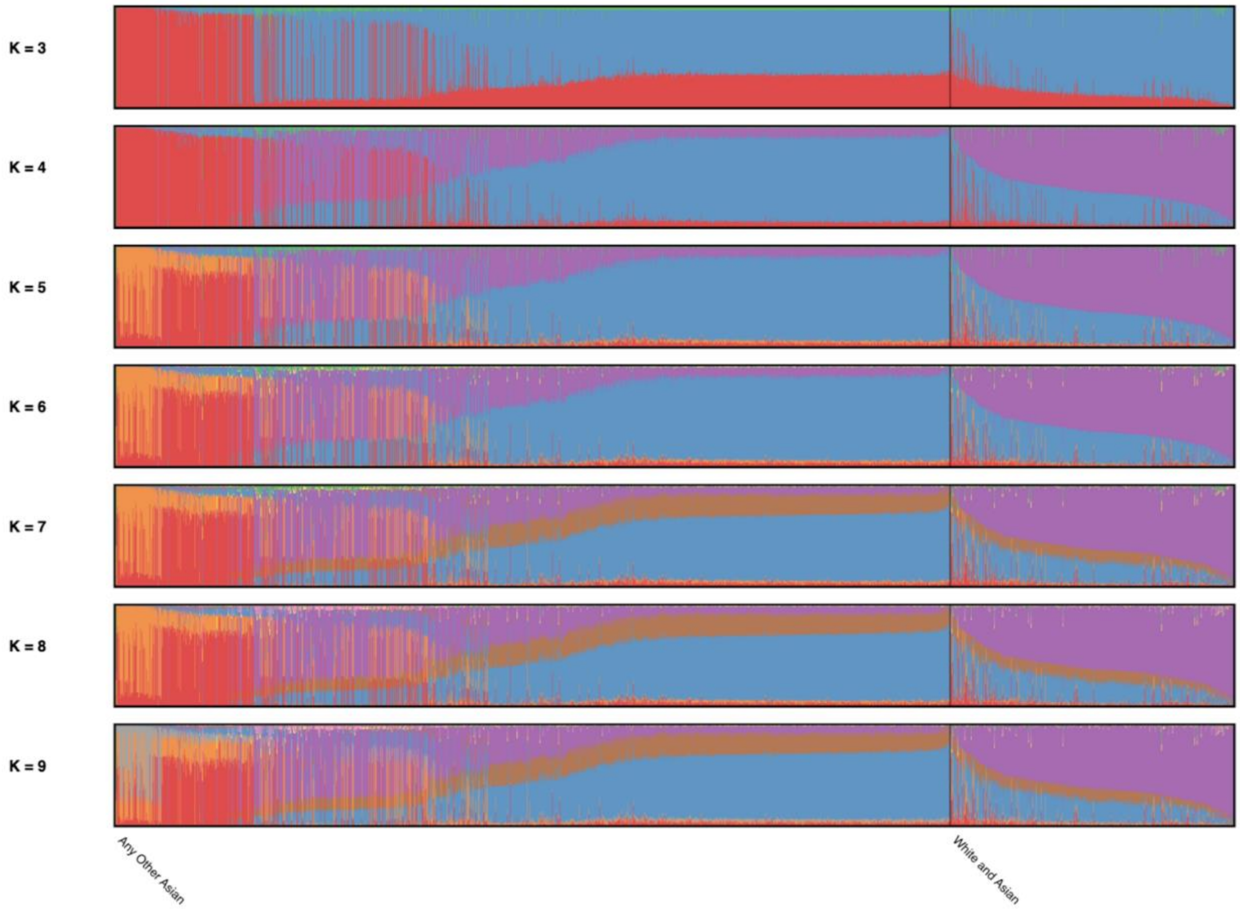

**Figure S1.14.** *pong* visualization of supervised ADMIXTURE analyses (for $k$ = 3 - 9) using 1000 Genomes reference data to infer population affinity proportions of UKB "Any other Asian" and "White and Asian" participants. Colors are consistent with Fig. S1.12 and Table S1.11.

**Table S1.11.** Average percent cluster membership for *k* = 3-9, output by the supervised ADMIXTURE analyses for UKB "Any Other Asian" (AOA) and "White and Asian" (WA) participants.

Clusters are colored by the reference groups most representative of that cluster.

ADMIXTURE average percent cluster membership

| Group | *k* | | | | | | | | | |
|---|---|---|---|---|---|---|---|---|---|---|
| | | *Europe* | *Asia* | *Africa* | | | | | | |
| AOA | 3 | 56.6% | 40.4% | 3.0% | | | | | | |
| WA | | 81.2% | 16.7% | 2.1% | | | | | | |
| | | *Europe* | *South Asia* | *East Asia* | *Africa* | | | | | |
| AOA | 4 | 22.8% | 51.9% | 23.7% | 1.6% | | | | | |
| WA | | 61.0% | 30.3% | 7.3% | 1.5% | | | | | |
| | | *Europe* | *South Asia* | *East Asia* | *Japan* | *Africa* | | | | |
| AOA | 5 | 22.7% | 51.5% | 14.4% | 9.8% | 1.6% | | | | |
| WA | | 60.8% | 30.0% | 4.8% | 2.9% | 1.5% | | | | |
| | | *Europe* | *South Asia* | *East Asia* | *Japan* | *Africa - GWD* | *Africa - LWK* | | | |
| AOA | 6 | 22.7% | 51.5% | 14.4% | 9.8% | 1.6% | 1.1% | | | |
| WA | | 60.8% | 30.0% | 4.8% | 2.9% | 0.6% | 1.0% | | | |
| | | *Europe* | *South Asia* | *South Asia - GIH* | *East Asia* | *Japan* | *Africa - GWD* | *Africa - LWK* | | |
| AOA | 7 | 21.3% | 38.7% | 14.8% | 14.1% | 9.6% | 0.5% | 1.0% | | |
| WA | | 59.4% | 21.6% | 10.2% | 4.6% | 2.8% | 0.5% | 1.0% | | |
| | | *Europe* | *South Asia* | *South Asia - GIH* | *East Asia* | *Japan* | *Africa - GWD* | *Africa - LWK* | *Africa - ESN* | |
| AOA | 8 | 21.3% | 38.7% | 14.7% | 14.1% | 9.6% | -- | 1.0% | -- | |
| WA | | 59.3% | 21.6% | 10.1% | 4.5% | 2.8% | -- | 0.7% | 0.5% | |
| | | *Europe* | *South Asia* | *South Asia - GIH* | *East Asia - China* | *South East Asia* | *Japan* | *Africa - GWD* | *Africa - LWK* | *Africa - ESN* |
| AOA | 9 | 21.2% | 38.5% | 14.3% | 8.2% | 11.1% | 5.2% | -- | 1.0% | -- |
| WA | | 59.2% | 21.5% | 9.8% | 2.8% | 3.6% | 1.4% | -- | 0.7% | 0.5% |

**SECTION 2: GWAS and PGS**
**2.1 UKB South Asian height data**

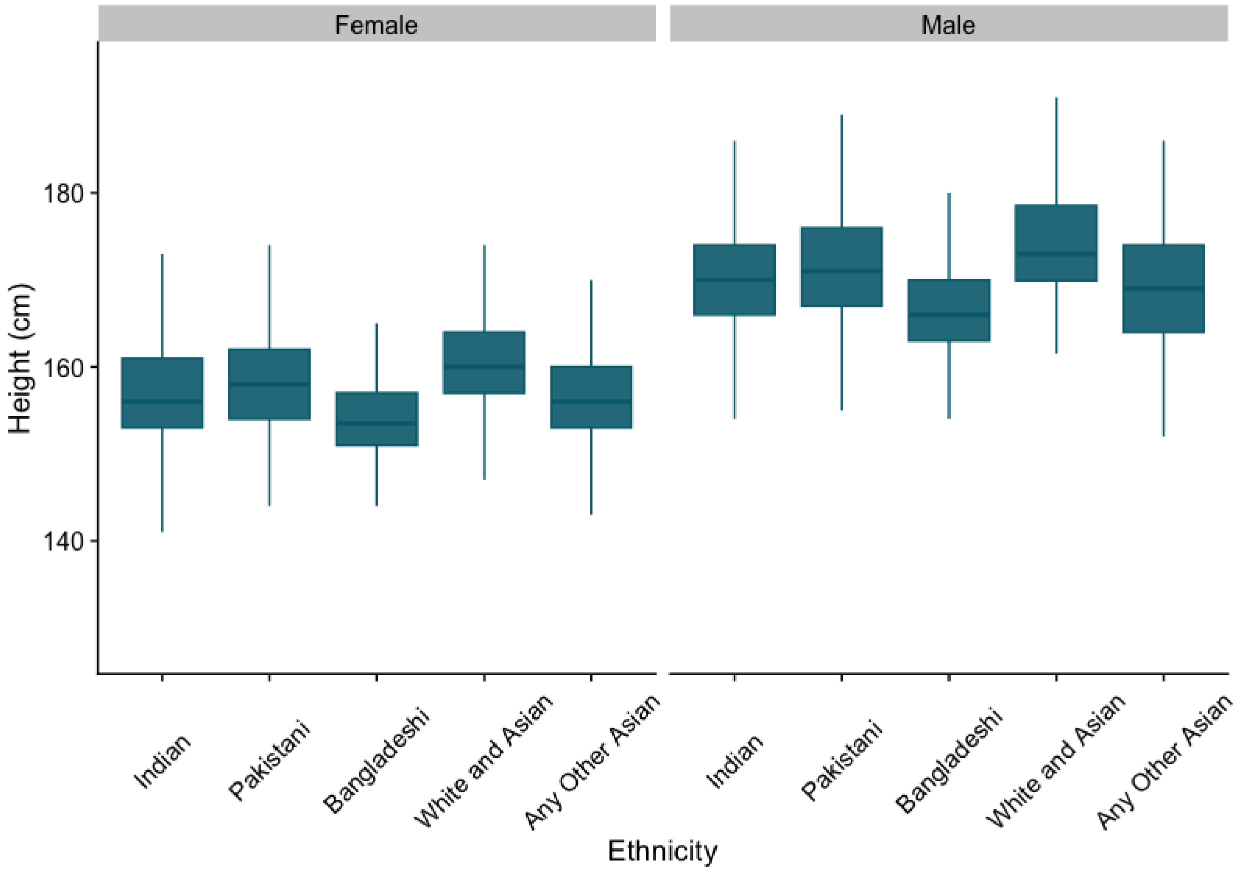

**Figure S2.1**. Height (cm) of UKB South Asian participants. “Ethnicity” reflects the UKB self-selected ethnic category (UKB data field 21000).

### 2.2: Environmental covariates
#### 2.2.1: Methods

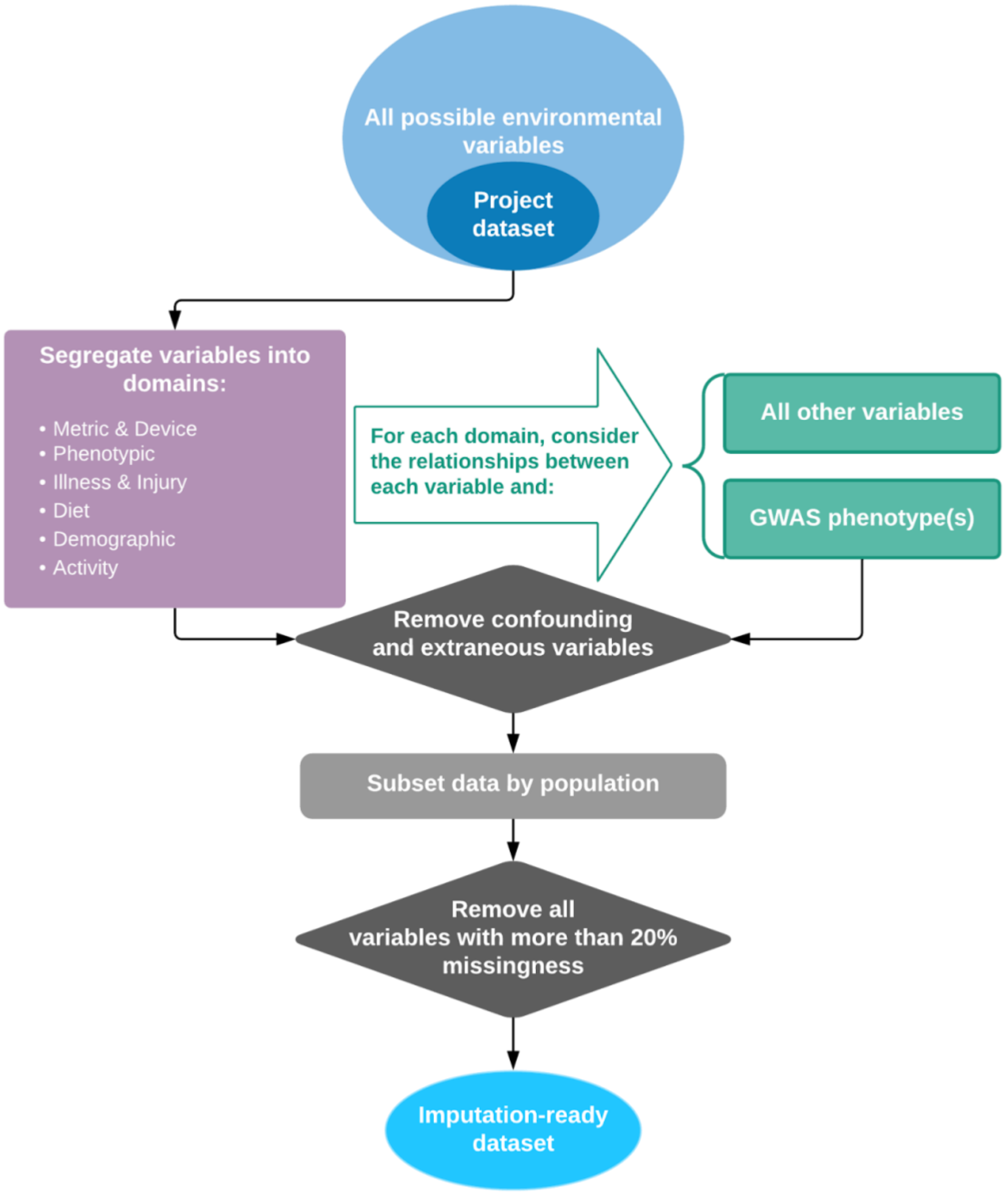

**Figure S2.2**. Environmental covariate QC workflow. Detail is provided in the main text Methods section.

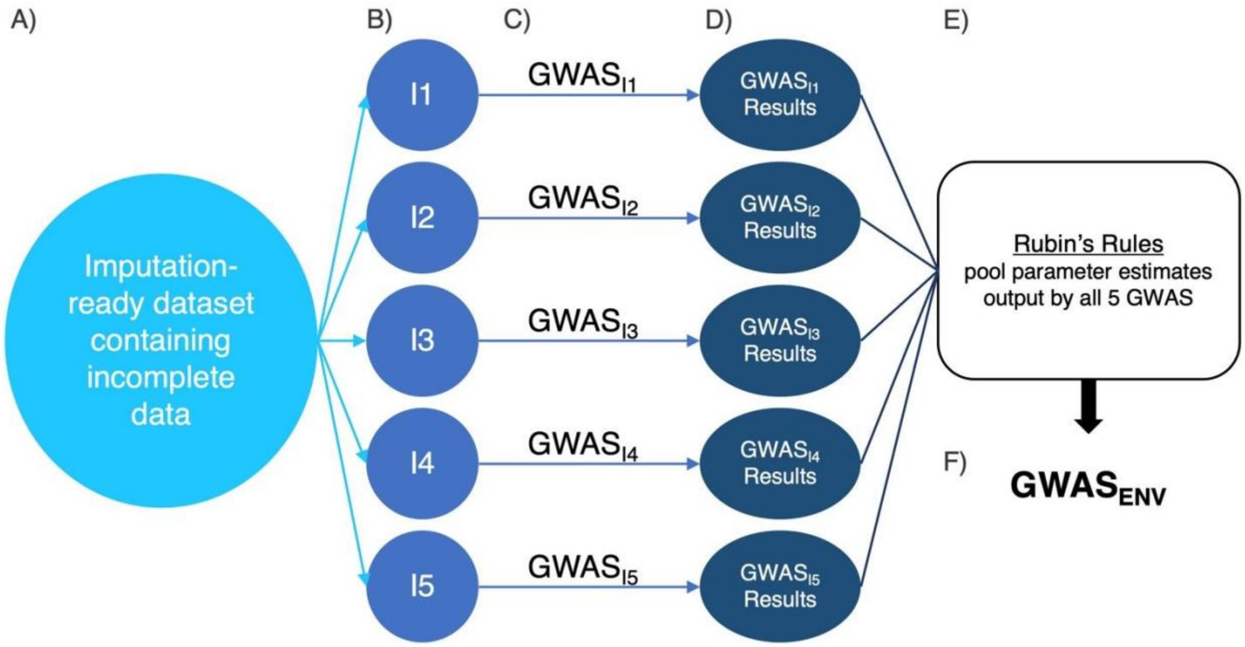

**Figure S2.3**. Environmental covariate multiple imputation (MI) procedure. A) Incomplete UKB data QC-ed as outlined in Fig. S2.2 are used to create five complete datasets (B) via classification and regression trees (CART) with R package 'MICE' [3, 4]; C) GWAS covariates are then extracted from each completed dataset and five identical GWAS are run (i.e. one GWAS per impute), resulting in a set of five GWAS summary statistics (D); E) Lastly, Rubin's rules are used to pool the parameter estimates output by each GWAS, creating new p-values, ®, and ® standard errors (Little and Rubin, 2019 [5]: section 5.4); F) $GWAS_{ENV}$ reflects the pooled results of $GWAS_{I1-5}$.

#### 2.2.2: Extended results

**Table S2.1.** Summary statistics of an environmental-only OLS linear regression model for height (*n* = 7,331):
$Height_i = \beta_0 + \beta_1(Sex_i) + \beta_2(TDI_i) + \beta_3(YOB_i) + \beta_4(\text{Birth location}_i) + \beta_5(\text{Eats pork}_i) + \beta_6(\text{Eats beef}_i) + \beta_7(\text{Excludes dairy}_i) + \beta_8(\text{Health}_i) + \beta_9(\text{Number of live births}_i) + \epsilon_i$

| **Coefficient** | **Estimate** | **Std. Error** | ***P*** |
|---|---|---|---|
| Intercept | -76.18 | 17.45 | < 0.0000 |
| Sex - Male | 13.42 | 0.22 | < 0.0000 |
| TDI | -0.14 | 0.02 | < 0.0000 |
| YOB | 0.12 | 0.01 | < 0.0000 |
| Birth location - Elsewhere | -1.90 | 0.23 | < 0.0000 |
| Eats pork - linear trend* | 0.32 | 0.12 | 0.0064 |
| Eats beef - linear trend* | 0.60 | 0.11 | < 0.0000 |
| Excludes dairy - linear trend* | -1.13 | 0.32 | 0.0004 |
| Health - linear trend* | 0.51 | 0.18 | 0.0035 |
| Number of births | -0.02 | 0.07 | 0.7732 |

*R standard orthogonal polynomial conversion (contrasts) for ordinal factor variables.

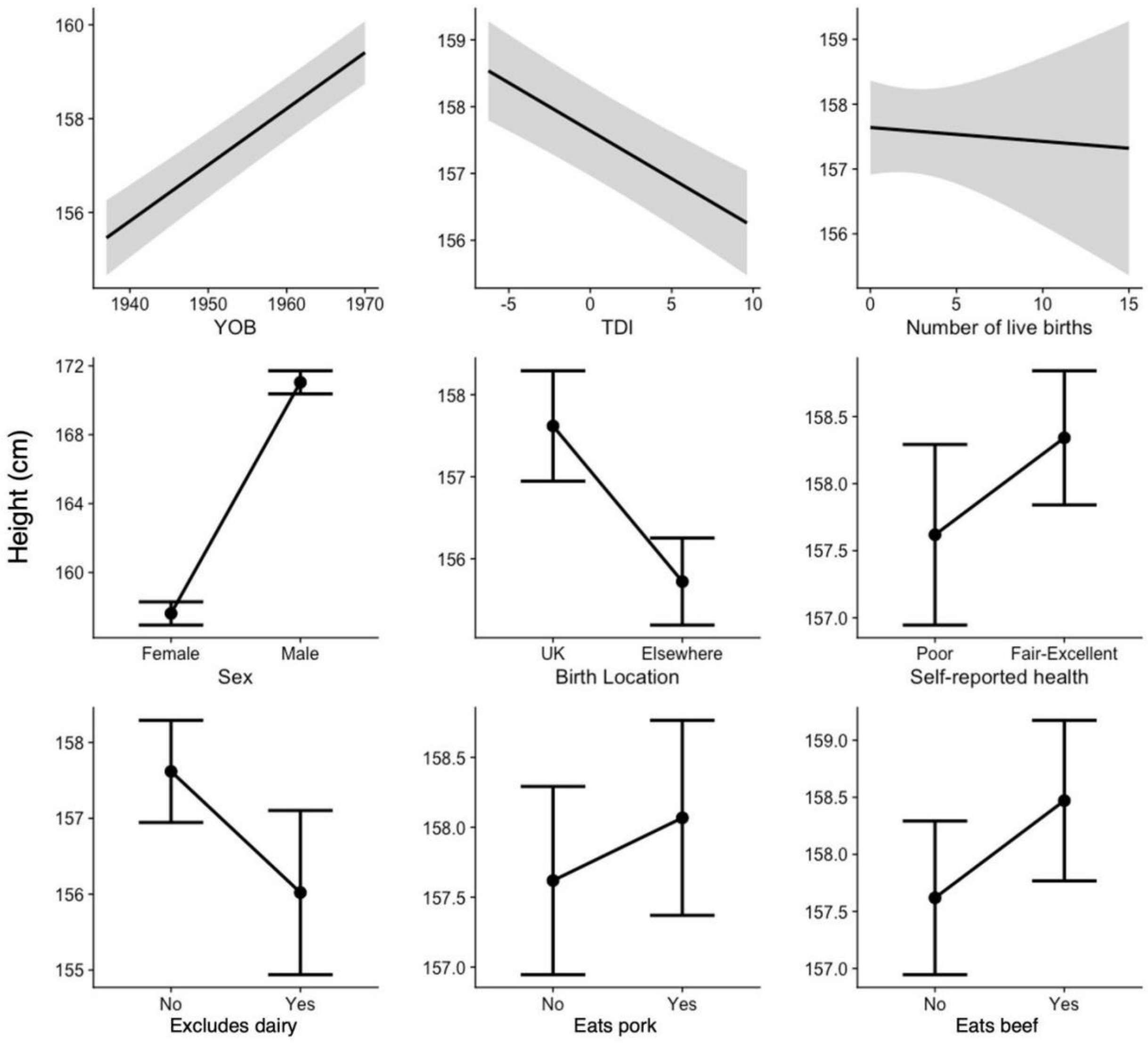

**Figure S2.4.** Effects on height for each environmental variable included as a covariate in $GWAS_{env}$. All non-genetic covariates were included in this environmental-only model, utilizing un-imputed data within the full UKB South Asian sample. The R statistical package 'jtools' was used to visualize these effects [6].

**Table S2.2.** Summary statistics of sex-specific environmental-only OLS linear regression models for height ($n_F$ = 3,430; $n_M$ = 3,901).

| Coefficient | Estimate | | Std. Error | | *P* | |
|---|---|---|---|---|---|---|
| | F | M | F | M | F | M |
| Intercept | -82.86 | -60.53 | 25.36 | 24.10 | 0.0011 | 0.0119 |
| TDI | -0.14 | -0.15 | 0.03 | 0.03 | < 0.0000 | < 0.0000 |
| YOB | 0.12 | 0.12 | 0.01 | 0.01 | < 0.0000 | < 0.0000 |
| Birth location - Elsewhere | -1.90 | -1.86 | 0.31 | 0.35 | < 0.0000 | < 0.0000 |
| Eats pork - linear trend* | 0.18 | 0.39 | 0.17 | 0.16 | 0.2874 | 0.0141 |
| Eats beef - linear trend* | 0.81 | 0.47 | 0.17 | 0.15 | < 0.0000 | 0.0021 |
| Excludes dairy - linear trend* | -0.50 | -1.70 | 0.44 | 0.46 | 0.2538 | 0.0002 |
| Health - linear trend* | 0.53 | 0.47 | 0.24 | 0.25 | 0.0270 | 0.0643 |
| Number of births (female only) | -0.02 | | 0.07 | | 0.7516 | |

*R base orthogonal polynomial conversion (contrasts) for ordinal factor variables.

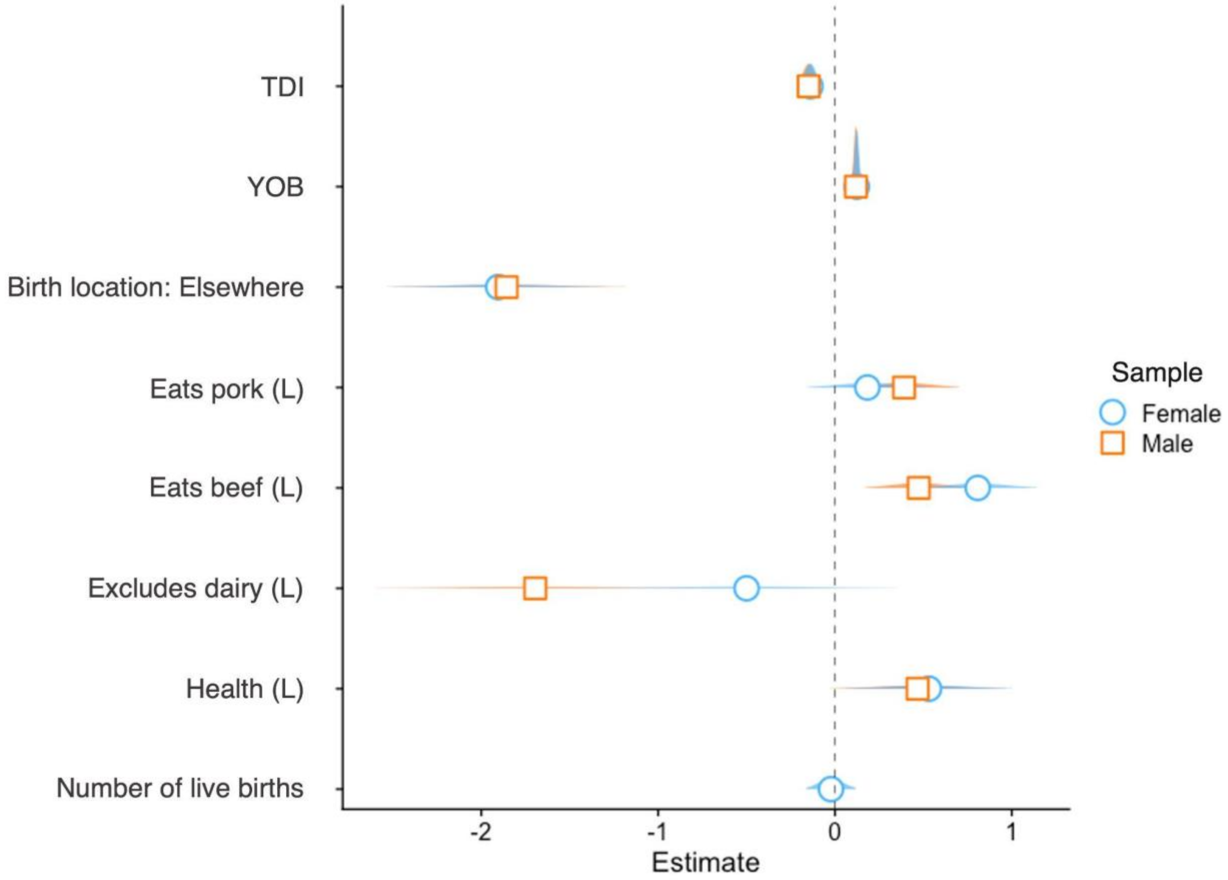

**Figure S2.5.** GWAS model environmental coefficient effects on female (blue / circle) and male (orange / square) participants. "L" refers to the statistical program R's base orthogonal polynomial conversion (contrasts) for ordinal factor variables.

### 2.3: GWAS extended results

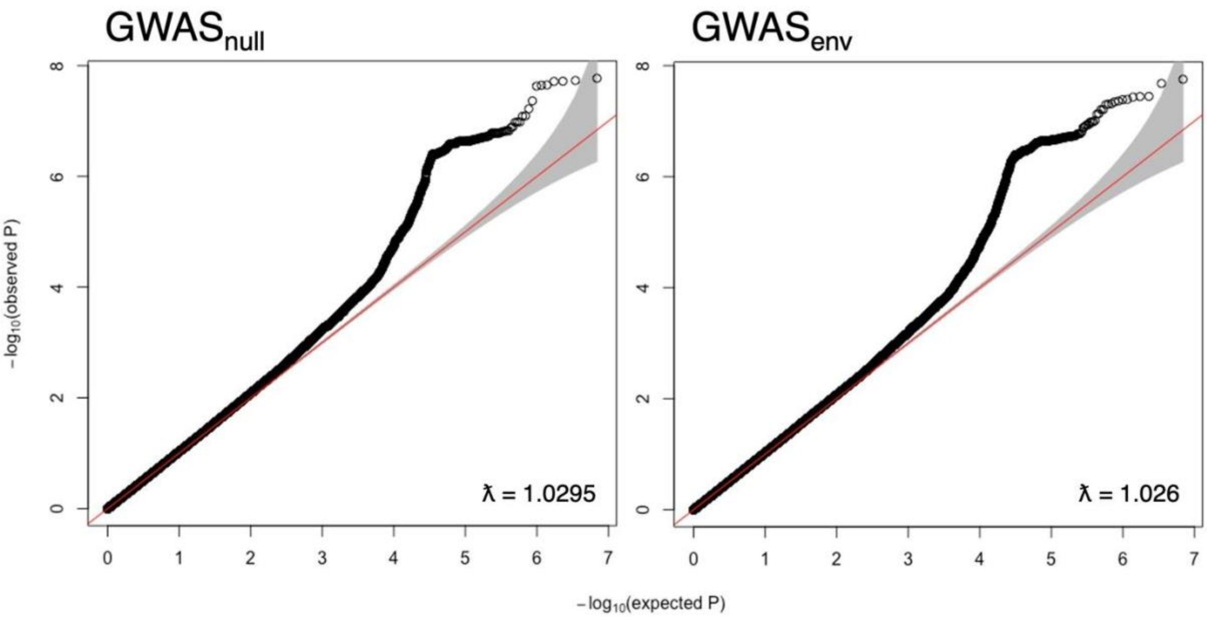

**Fig S2.6**: GWAS$_{null}$ and GWAS$_{env}$ QQ plots and genomic inflation (λ).

**Table S2.3**. The top two variants per chromosome with the largest relative differences in beta values between GWAS outputs, for loci significant at $p < 10^{-6}$.

| **SNP** | **Chr** | **Position*** | **Nearest gene(s)†** | **Effect allele** | **Other allele** | **$GWAS_{null}$** | | **$GWAS_{env}$** | | **$\|(\beta_{null}-\beta_{env}) / \beta_{null}\|$**** |
|---|---|---|---|---|---|---|---|---|---|---|
| | | | | | | **β** | **s.e.** | **β** | **s.e.** | |
| rs9822195 | 3 | 141096736 | *ZBTB38* | T | C | 0.661 | 0.128 | 0.650 | 0.126 | 1.53% |
| rs114097136 | 3 | 141114013 | *ZBTB38* | G | C | 1.108 | 0.226 | 1.134 | 0.223 | 2.31% |
| rs186069885 | 7 | 75978178 | *YWHAG* | C | T | 1.547 | 0.321 | 1.621 | 0.317 | 4.75% |
| rs192457049 | 7 | 76036583 | *ZP3, SRCRB4D* | A | T | 1.517 | 0.309 | 1.613 | 0.306 | 6.33% |
| rs71519447 | 8 | 57157543 | *CHCHD7, RP11-140I16.2* | A | G | -0.862 | 0.153 | -0.832 | 0.152 | 3.49% |
| rs67742458 | 8 | 57170647 | *RP11-140I16.2, SDR16C5* | G | A | -0.851 | 0.152 | -0.822 | 0.151 | 3.45% |
| rs224331 | 20 | 34022387 | *GDF5OS:GDF5* | C | A | 0.491 | 0.101 | 0.516 | 0.100 | 5.02% |
| rs143384 | 20 | 34025756 | *GDF5* | G | A | 0.551 | 0.102 | 0.563 | 0.100 | 2.26% |

*GRCh37 (hg19)
** This metric is simply the absolute percent difference in effect sizes (last column).
†If one gene is listed, then the variant is located within that gene. If two genes are listed, the variant is intergenic.

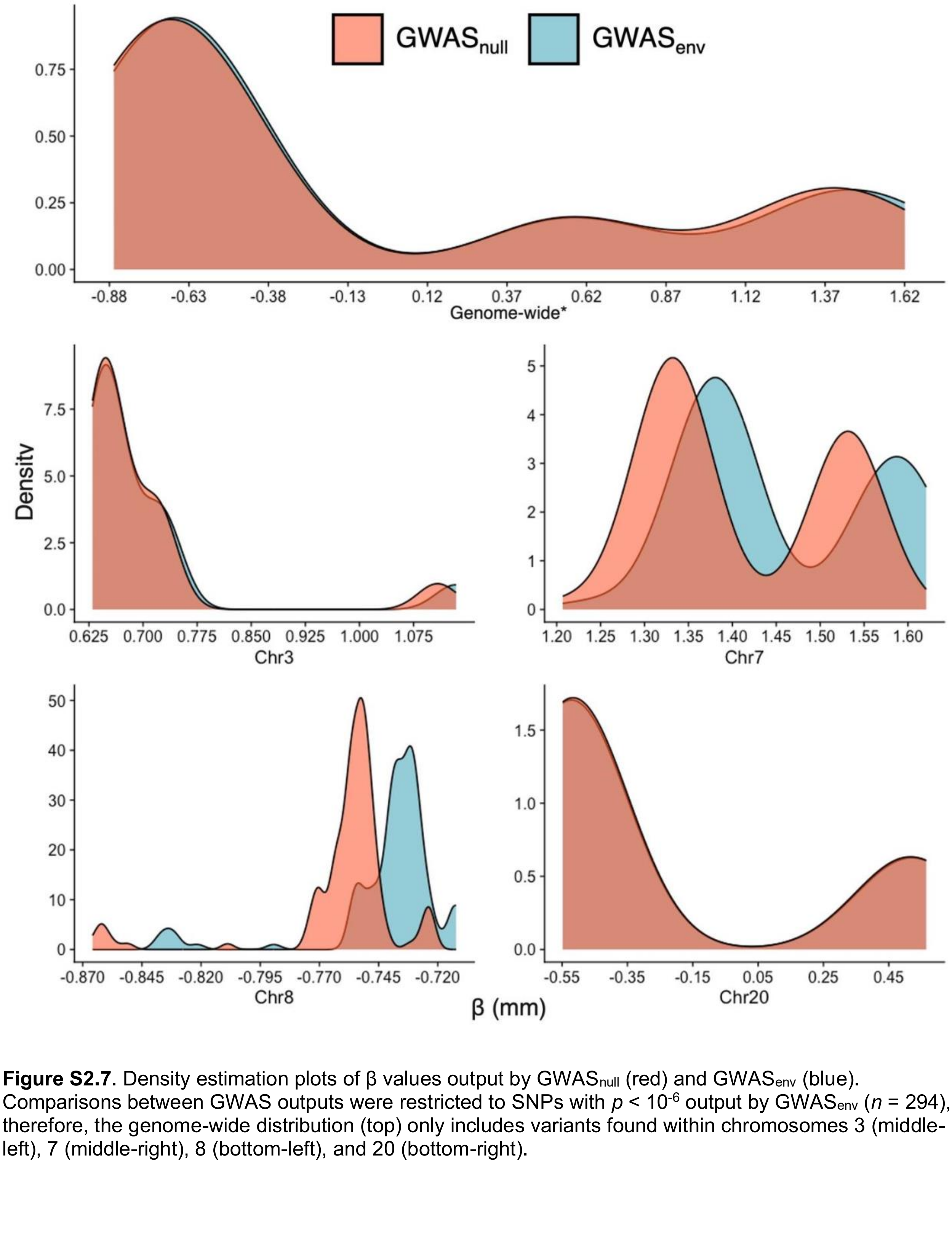

**Figure S2.7**. Density estimation plots of β values output by GWAS$_{null}$ (red) and GWAS$_{env}$ (blue). Comparisons between GWAS outputs were restricted to SNPs with $p < 10^{-6}$ output by GWAS$_{env}$ ($n$ = 294), therefore, the genome-wide distribution (top) only includes variants found within chromosomes 3 (middle-left), 7 (middle-right), 8 (bottom-left), and 20 (bottom-right).

**Table S2.4**. Summary of effect size changes from $GWAS_{null}$ to $GWAS_{env}$ ($p < 10^{-6}$, $n = 294$).

| **Effect change** | ***n*** |
|---|---|
| Negative effects strengthened | 65 |
| Negative effects weakened | 129 |
| Positive effects reduced | 2 |
| Positive effects increased | 98 |

**Table S2.5**. β SE differences between $GWAS_{null}$ and $GWAS_{env}$ summary statistics, per p-value threshold. Differences were calculated by subtracting $SE_{env}$ from $SE_{null}$ for each SNP meeting the significance threshold. The final column, "$n$ $SE_{env} > SE_{null}$" lists how many variants had larger SE in $GWAS_{env}$ relative to $GWAS_{null}$.

| ***p* <** | **Min** | **Mean** | **Max** | **$n$ $SE_{env} > SE_{null}$** |
|---|---|---|---|---|
| **1** | -5.53 | 0.007 | 0.49 | 5750 |
| **$10^{-6}$** | 0.001 | 0.002 | 0.01 | 0 |

### 2.3.1: Functional annotation

FUMA-GWAS [7] was used to summarize functional annotations of the $GWAS_{env}$ output and assess overlap with SNP associations currently reported in the GWAS Catalog [8]. At the SNP-level, three independent loci meet the genome-wide significance threshold ($p < 5^{-8}$), and 466 candidate SNPs (SNPs found in 1KG reference data that are in LD [$R^2 \geq 0.6$] with one of the significant independent SNPs) were identified (Tables S2.6 and S2.7). Of the candidate SNPs, 111 (~24%) are not currently reported for an association with height in the GWAS Catalog. GWAS SNPs were mapped to 18,687 protein coding genes, which were then used as the input for gene-based tests computed by MAGMA [9]. This identified three genes below the genome-wide significance threshold ($p < 2.68^{-6}$): *UQCC1*, *GDF5*, and *ACAN* (Tables S2.8 and S2.9; Figure S2.8 and S2.9). Each of these genes has previously been associated with height, skeletal disorders, growth, and/or cartilage composition ([10]; GeneCards – the human gene database [www.genecards.org]).

**Table S2.6.** Overview of FUMA-GWAS 'SNP2GENE' results for $GWAS_{env}$ summary statistics. SNP2GENE parameters are listed in Table S2.10.

| **Result** | **Description** | ***n*** |
|---|---|---|
| Genomic risk loci | Loci derived from independent significant SNPs by merging LD blocks within 250kb of each other. | 3 |
| Independent significant SNPs | SNPs with genome-wide significance ($p < 5e^{-8}$) and are independent from each other ($R^2 < 0.6$). | 3 |
| Lead SNPs | Independent significant SNPs with $R^2 \leq 0.1$. | 3 |
| Candidate SNPs | SNPs in LD with one of the independent significant SNPs, including variants not included in the GWAS data but included in the 1000 Genomes SAS reference panel. | 466 |
| Candidate GWAS tagged SNPs | Candidate SNPs that exist in the GWAS data. | 355 |
| Mapped genes | Genes with candidate SNPs located on the gene body or within 1kb of the gene transcription start site. | 10 |

**Table S2.7.** FUMA-derived lead SNPs with associated $GWAS_{env}$ summary statistics.

| SNP | Chr | Position | Effect allele | Other allele | β (mm) | Nearest gene |
|---|---|---|---|---|---|---|
| rs13091182 | 3 | 141133960 | A | G | 0.73 | *ZBTB38* |
| rs36063247 | 8 | 57154653 | C | T | -0.84 | *RP11-140I16.2* |
| rs143383 | 20 | 34025983 | A | G | 0.57 | *GDF5* |

**Table S2.8.** MAGMA gene-based test results as computed using $GWAS_{env}$ summary statistics in FUMA-GWAS.

| Gene | Ensembl ID | Chr | Position (start – stop) | Number of SNPs mapped to gene | *p* |
|---|---|---|---|---|---|
| *ZBTB38* | ENSG00000177311 | 3 | 141043055 - 141168634 | 250 | $2.77^{-6}$ |
| *HHIP* | ENSG00000164161 | 4 | 145567173 - 145666423 | 164 | $2.17^{-5}$ |
| *ABCE1* | ENSG00000164163 | 4 | 146019084 - 146050331 | 21 | $8.19^{-5}$ |
| *SRRM3* | ENSG00000177679 | 7 | 75831216 - 75916605 | 244 | $2.06^{-5}$ |
| *PLAG1* | ENSG00000181690 | 8 | 57073463 - 57123883 | 78 | $5.95^{-5}$ |
| *RBMXL2* | ENSG00000170748 | 11 | 7110165 - 7112379 | 5 | $8.17^{-5}$ |
| *ACAN** | ENSG00000157766 | 15 | 89346674 - 89418585 | 212 | $1.75^{-6}$ |
| *EDEM2* | ENSG00000088298 | 20 | 33703167 - 33865928 | 401 | $3.47^{-5}$ |
| *UQCC1** | ENSG00000101019 | 20 | 33890369 - 33999944 | 163 | $2.91^{-7}$ |
| *GDF5OS* | ENSG00000204183 | 20 | 34020827 - 34023248 | 6 | $8.23^{-5}$ |
| *GDF5** | ENSG00000125965 | 20 | 34021145 - 34042568 | 35 | $2.98^{-7}$ |
| *CEP250* | ENSG00000126001 | 20 | 34042985 - 34099804 | 92 | $1.05^{-5}$ |
| *KCTD17* | ENSG00000100379 | 22 | 37447779 - 37459430 | 3 | $8.68^{-5}$ |

* Genes meeting genome-wide significance ($p < 2.68^{-6}$). Input SNPs were mapped to 18,687 protein coding genes, but only those with $p < 1^{-4}$ are reported here.

**Table S2.9**. Overrepresentation of $GWAS_{env}$ output genes found in GWAS-catalog gene set, as calculated within the FUMA-GWAS pipeline [7, 8].

| **Phenotype** | ***n****** | **p-value†** | **Adj. *p*‡** | **Genes§** |
|---|---|---|---|---|
| Height | 1013 | $6.45e^{-10}$ | $2.85e^{-6}$ | *ZBTB38, PLAG1, CHCHD7, SDR16C5, FAM83C, UQCC1, GDF5, CEP250* |
| Hip circumference adjusted for BMI | 776 | $7.16e^{-9}$ | $1.58e^{-5}$ | *ZBTB38, PLAG1, CHCHD7, SDR16C5, FAM83C, UQCC1, GDF5* |
| Waist-to-hip ratio adjusted for BMI (age > 50) | 178 | $9.83e^{-7}$ | $1.45e^{-3}$ | *FAM83C, GDF5, CEP250, C20orf173* |
| Waist-to-hip ratio adjusted for BMI x sex x age interaction (4df test) | 299 | $7.73e^{-6}$ | $8.55e^{-3}$ | *FAM83C, GDF5, CEP250, C20orf173* |
| Refractive error | 1381 | $1.14e^{-5}$ | $1.01e^{-2}$ | *ZBTB38, PLAG1, CHCHD7, SDR16C5, GDF5, CEP250* |
| Infant length | 13 | $1.62e^{-5}$ | $1.02e^{-2}$ | *ZBTB38, GDF5* |
| Low hand grip strength (60 years and older) (EWGSOP) | 13 | $1.62e^{-5}$ | $1.02e^{-2}$ | *ZBTB38, GDF5* |
| Body fat distribution (leg fat ratio) | 167 | $5.73e^{-5}$ | $2.89e^{-2}$ | *ZBTB38, FAM83C, GDF5* |
| Brain morphology (MOSTest) | 1049 | $5.88e^{-5}$ | $2.89e^{-2}$ | *FAM83C, UQCC1, GDF5, CEP250, C20orf173* |
| Body fat distribution (trunk fat ratio) | 179 | $7.05e^{-5}$ | $3.12e^{-2}$ | *ZBTB38, FAM83C, GDF5* |

**n:* Number of genes associated with the phenotype in GWAS-catalog.
†p-value: Hypergeometric test (upper tail) p-value.
‡Adj. *p*: Adjusted p-value (false discovery rate - Benjamini and Hochberg step-up procedure [FDR-BH]).
§Genes: $GWAS_{env}$ genes overlapping with GWAS-catalog.

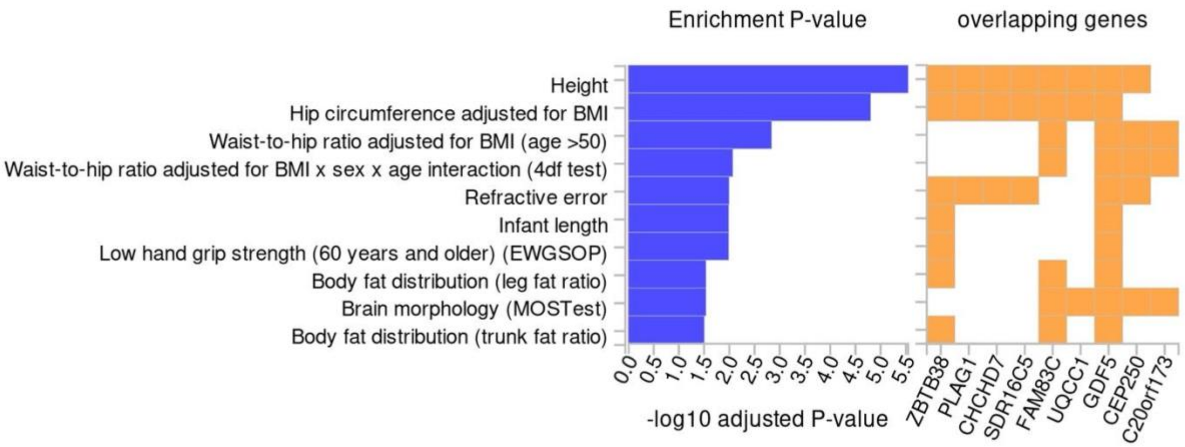

**Figure S2.8**: Enrichment of $GWAS_{env}$ output genes relative to the GWAS-catalog gene set, with adjusted p-value < 0.05. Modified from FUMA-GWAS results, June 2024 [7, 8].

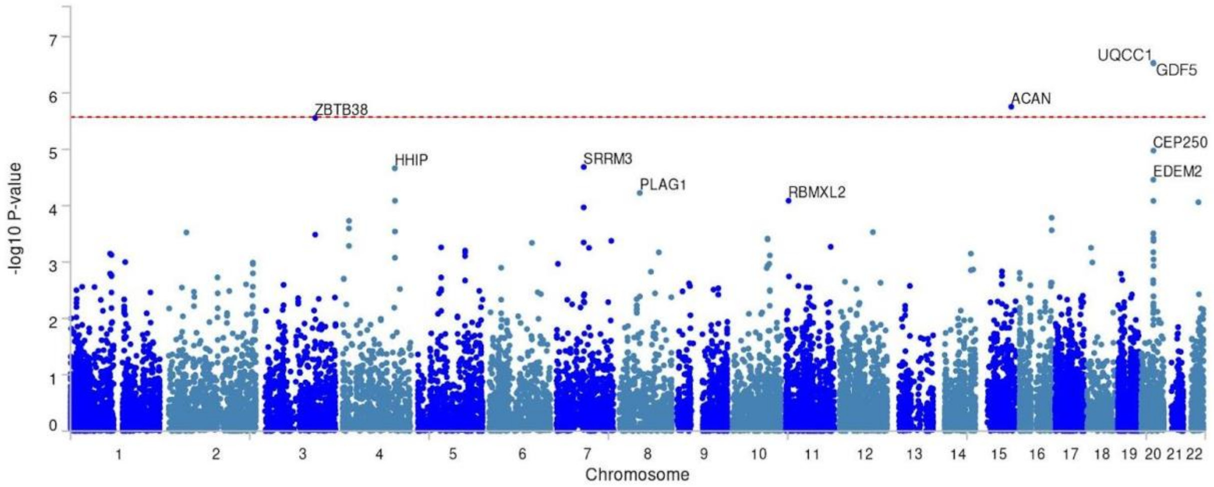

**Figure S2.9**: Gene-based Manhattan plot (10 genes labeled) as output from FUMA-GWAS.

**Table S2.10:** FUMA-GWAS SNP2GENE and GENE2FUNC information and relevant parameters.

| | | | |
|---|---|---|---|
| **Versions** | FUMA = v1.5.2 | | |
| | MAGMA = v1.08 | | |
| | GWAScatalog = e0_r2022-11-29 | | |
| | ANNOVAR = 2017-07-17 | | |
| **SNP2GENE parameters** | GRCh38 = 0 | **Positional mapping** | posMap = 1 |
| | N = 6937 | | posMapWindowSize = 10 |
| | Ncol = NA | | posMapAnnot = NA |
| | exMHC = 1 | | posMapCADDth = 0 |
| | MHCopt = annot | | posMapRDBth = NA |
| | extMHC = NA | | posMapChr15 = NA |
| | ensembl = v102 | | posMapChr15Max = NA |
| | genetype = protein_coding | | posMapChr15Meth = NA |
| | leadP = 5e-8 | | posMapAnnoDs = NA |
| | gwasP = 0.05 | | posMapAnnoMeth = NA |
| | r2 = 0.6 | **GENE2FUNC parameters** | gtype = text |
| | r2_2 = 0.1 | | gval = ENSG00000177311:ENSG00000181690:ENSG00000170791:ENSG00000170786:ENSG00000125998:ENSG00000101019:ENSG00000204183:ENSG00000125965:ENSG00000126001:ENSG00000125975 |
| | refpanel = 1KG/Phase3 | | bkgtype = select |
| | pop = SAS | | bkgval = protein_coding |
| | MAF = 0 | | MHC = 1 |
| | refSNPs = 1 | | ensembl = v102 |
| | mergeDist = 250 | | gsFileN = 0 |
| **MAGMA** | magma = 1 | | gsFiles = NA |
| | magma_window = 0 | | gene_exp = GTEx/v8/gtex_v8_ts_avg_log2TPM:GTEx/v8/gtex_v8_ts_general_avg_log2TPM |
| | magma_exp = GTEx/v8/gtex_v8_ts_avg_log2TPM:GTEx/v8/gtex_v8_ts_general_avg_log2TPM | | adjPmeth = fdr_bh |
| | | | adjPcut = 0.05 |
| | | | minOverlap = 2 |

### 2.4: PGS

#### 2.4.1: Methods

**Table S2.11**. PGS model harmonization summaries. All PGS scores were applied using ESCALATOR (Lin and Fisher, 2022: [https://github.com/menglin44/ESCALATOR]).

| **PGS** | **Variant information** |
|---|---|
| $PGS_{Y1}$ (Yengo PGS x GWAS SNPs) | 89,236 ambiguous A/T, C/G loci removed |
| | 169,019 SNPs not found in array |
| | 898,486 variants used |
| $PGS_{Y2}$ (Yengo PGS x all possible SNPs) | 89,236 ambiguous A/T, C/G loci removed |
| | 34,833 SNPs not found in array |
| | 1,032,672 variants used |

#### 2.4.2: Extended results

**Table S2.12**. Detailed predictive performance of Yengo et al. PGS equations and associated non-genetic and/or environmental models. Confidence intervals were calculated using package *psychometric* in R [https://CRAN.R-project.org/package=psychometric].

| Model | Model (R syntax) | Model description | Sample | $R^2$ | $R^2$ 95% CI | *p* |
|---|---|---|---|---|---|---|
| **Baseline covariates** | Height ~ Sex + YOB | Height explained by $GWAS_{null}$ covariates | Full test sample (*n* = 997) | 0.549 | 0.508 - 0.590 | $< 2.2 \times 10^{-16}$ |
| **$PGS_{Y1}$: PGS002800 – GWAS SNP intersection (*n* = 898,486)** *SNPs in the Yengo et al. PGS weight file and our GWAS model* | Height ~ Score | Height explained by only PGS scores (no other adjustments or covariates) | Full test sample (*n* = 997) | 0.027 | 0.008 - 0.047 | $1.43 \times 10^{-7}$ |
| | | | F (*n* = 473) | 0.099 | 0.048 - 0.150 | $2.51 \times 10^{-12}$ |
| | | | M (*n* = 524) | 0.042 | 0.009 - 0.076 | $1.88 \times 10^{-6}$ |
| | Height ~ Score + Sex + YOB | Height explained by PGS scores plus comparable covariates used in PGS002800 training model | Full test sample (*n* = 997) | 0.577 | 0.537 - 0.616 | $< 2.2 \times 10^{-16}$ |
| | Height ~ Score + YOB | | F (*n* = 473) | 0.142 | 0.085 - 0.200 | $< 2.2 \times 10^{-16}$ |
| | | | M (*n* = 524) | 0.070 | 0.028 - 0.112 | $6.37 \times 10^{-9}$ |
| **$PGS_{Y2}$: PGS002800 – "full" SNP intersection (*n* = 1,032,672)** *SNPs in the Yengo et al. PGS weight file and our imputed data (regardless of QC filtering thresholds)* | Height ~ Score | Height explained by only PGS scores (no other adjustments or covariates) | Full test sample (*n* = 997) | 0.024 | 0.005 - 0.042 | $9.18 \times 10^{-7}$ |
| | | | F (*n* = 473) | 0.104 | 0.053 - 0.156 | $5.8 \times 10^{-13}$ |
| | | | M (*n* = 524) | 0.041 | 0.008 - 0.074 | $3.16 \times 10^{-6}$ |
| | Height ~ Score + Sex + YOB | Height explained by PGS scores plus comparable covariates used in PGS002800 training model | Full test sample (*n* = 997) | 0.577 | 0.537 - 0.616 | $< 2.2 \times 10^{-16}$ |
| | Height ~ Score + YOB | | F (*n* = 473) | 0.148 | 0.089 - 0.206 | $< 2.2 \times 10^{-16}$ |
| | | | M (*n* = 524) | 0.068 | 0.027 - 0.109 | $1.09 \times 10^{-8}$ |
| **Extended environmental covariates** | Height ~ Sex + TDI + YOB + Birth location + Eats pork + Eats beef + Excludes dairy + Health + Number of births | Height explained by $GWAS_{env}$ covariates | Full test sample with unimputed covariate data (*n* = 907) | 0.569 | 0.527 - 0.611 | $< 2.2 \times 10^{-16}$ |
| **$PGS_{Y1}$** | Height ~ Score + Sex + TDI + YOB + Birth location + Eats pork + Eats beef + Excludes dairy + Health + Number of births | Height explained by PGS scores plus $GWAS_{env}$ covariates | Full test sample with unimputed covariate data (*n* = 907) | 0.597 | 0.557 - 0.637 | $< 2.2 \times 10^{-16}$ |
| | Height ~ Score + TDI + YOB + Birth location + Eats pork + Eats beef + Excludes dairy + Health + Number of births | | F (*n* = 427) | 0.169 | 0.106 - 0.232 | $3.74 \times 10^{-13}$ |
| | Height ~ Score + TDI + YOB + Birth location + Eats pork + Eats beef + Excludes dairy + Health | | M (*n* = 480) | 0.110 | 0.058 - 0.161 | $4.169 \times 10^{-9}$ |
| **$PGS_{Y2}$** | Height ~ Score + Sex + TDI + YOB + Birth location + Eats pork + Eats beef + Excludes dairy + Health + Number of births | Height explained by PGS scores plus $GWAS_{env}$ covariates | Full test sample with unimputed covariate data (*n* = 907) | 0.597 | 0.557 - 0.637 | $< 2.2 \times 10^{-16}$ |
| | Height ~ Score + TDI + YOB + Birth location + Eats pork + Eats beef + Excludes dairy + Health + Number of births | | F (*n* = 427) | 0.175 | 0.111 - 0.238 | $1.1 \times 10^{-13}$ |
| | Height ~ Score + TDI + YOB + Birth location + Eats pork + Eats beef + Excludes dairy + Health | | M (*n* = 480) | 0.108 | 0.057 - 0.159 | $6.75 \times 10^{-9}$ |

**Table S2.13**. Detailed predictive performance of GWAS-derived PGS equations.

| PGS | Model (R syntax) | Model description | Sample | Adj. $R^2$ | $R^2$ | $R^2$ 95% CI | *p* |
|---|---|---|---|---|---|---|---|
| **GWAS$_{null}$ PGS (*n* = 896,160)** | Height ~ Score | Height explained by only PGS scores (no other adjustments or covariates) | Full test sample (*n* = 997) | 0.025 | 0.026 | 0.006 - 0.045 | $3.46 \times 10^{-7}$ |
| | | | F (*n* = 473) | 0.068 | 0.070 | 0.026 - 0.113 | $5.62 \times 10^{-9}$ |
| | | | M (*n* = 524) | 0.045 | 0.047 | 0.012 - 0.082 | $5.61 \times 10^{-7}$ |
| | Height ~ Score + Sex + YOB | Height explained by PGS scores plus covariates used in discovery GWAS | Full test sample (*n* = 997) | 0.573 | 0.574 | 0.534 - 0.614 | $< 2.2 \times 10^{-16}$ |
| | Height ~ Score + YOB | | F (*n* = 473) | 0.116 | 0.120 | 0.065 - 0.174 | $9.31 \times 10^{-14}$ |
| | | | M (*n* = 524) | 0.074 | 0.077 | 0.034 - 0.121 | $7.98 \times 10^{-10}$ |
| | Height ~ Score + Sex + TDI + YOB + Birth location + Eats pork + Eats beef + Excludes dairy + Health + Number of births | Height explained by PGS scores plus covariates used in GWAS$_{env}$ | Full test sample with unimputed covariate data (*n* = 907) | 0.585 | 0.590 | 0.550 - 0.630 | $< 2.2 \times 10^{-16}$ |
| | Height ~ Score + TDI + YOB + Birth location + Eats pork + Eats beef + Excludes dairy + Health + Number of births | | F (*n* = 427) | 0.114 | 0.133 | 0.075 - 0.191 | $1.27 \times 10^{-9}$ |
| | Height ~ Score + TDI + YOB + Birth location + Eats pork + Eats beef + Excludes dairy + Health | | M (*n* = 480) | 0.095 | 0.110 | 0.058 - 0.161 | $4.26 \times 10^{-9}$ |
| **GWAS$_{env}$ PGS (*n* = 896,160)** | Height ~ Score | Height explained by only PGS scores (no other adjustments or covariates) | Full test sample (*n* = 997) | 0.021 | 0.022 | 0.004 - 0.040 | $2.14 \times 10^{-6}$ |
| | | | F (*n* = 473) | 0.056 | 0.058 | 0.017 - 0.098 | $1.17 \times 10^{-7}$ |
| | | | M (*n* = 524) | 0.038 | 0.040 | 0.007 - 0.073 | $3.75 \times 10^{-6}$ |
| | Height ~ Score + Sex + YOB | Height explained by PGS scores and covariates used in GWAS$_{null}$ model | Full test sample (*n* = 997) | 0.569 | 0.570 | 0.530 - 0.610 | $< 2.2 \times 10^{-16}$ |
| | Height ~ Score + Sex + TDI + YOB + Birth location + Eats pork + Eats beef + Excludes dairy + Health + Number of births | Height explained by PGS scores plus covariates used in discovery GWAS | Full test sample with unimputed covariate data (*n* = 907) | 0.582 | 0.587 | 0.546 - 0.628 | $< 2.2 \times 10^{-16}$ |
| | Height ~ Score + TDI + YOB + Birth location + Eats pork + Eats beef + Excludes dairy + Health + Number of births | | F (*n* = 427) | 0.104 | 0.123 | 0.065 - 0.180 | $1.16 \times 10^{-8}$ |
| | Height ~ Score + TDI + YOB + Birth location + Eats pork + Eats beef + Excludes dairy + Health | | M (*n* = 480) | 0.091 | 0.106 | 0.055 - 0.157 | $1.01 \times 10^{-8}$ |

**Table S2.14.** Results of testing for significant differences between PGS $R^2$. Significance testing was conducted using R package *r2redux*, function *r2_diff()* [https://cran.r-project.org/web/packages/r2redux/r2redux.pdf].

| Covariates included | PGS models compared | Sample | Mean diff.* | Upper diff.* | Lower diff.* | *p* |
|---|---|---|---|---|---|---|
| Sex + YOB | $GWAS_{env}$, $GWAS_{null}$ | All | -0.0091 | -0.0029 | -0.0153 | 0.0041 |
| | $GWAS_{env}$, Y1 | | -0.0138 | 0.0194 | -0.0469 | 0.4159 |
| | $GWAS_{env}$, Y2 | | -0.0142 | 0.0194 | -0.0477 | 0.4075 |
| | $GWAS_{null}$, Y1 | | -0.0047 | 0.0298 | -0.0392 | 0.7906 |
| | $GWAS_{null}$, Y2 | | -0.0051 | 0.0298 | -0.0400 | 0.7752 |
| YOB | $GWAS_{env}$, $GWAS_{null}$ | F | -0.0130 | -0.0037 | -0.0224 | 0.0065 |
| | $GWAS_{env}$, Y1 | | -0.0364 | 0.0174 | -0.0901 | 0.1845 |
| | $GWAS_{env}$, Y2 | | -0.0418 | 0.0134 | -0.0969 | 0.1377 |
| | $GWAS_{null}$, Y1 | | -0.0233 | 0.0328 | -0.0795 | 0.4155 |
| | $GWAS_{null}$, Y2 | | -0.0287 | 0.0288 | -0.0863 | 0.3277 |
| | $GWAS_{env}$, $GWAS_{null}$ | M | -0.0064 | 0.0018 | -0.0147 | 0.1263 |
| | $GWAS_{env}$, Y1 | | 0.0012 | 0.0421 | -0.0396 | 0.9534 |
| | $GWAS_{env}$, Y2 | | 0.0032 | 0.0440 | -0.0376 | 0.8774 |
| | $GWAS_{null}$ Y1 | | 0.0076 | 0.0500 | -0.0347 | 0.7237 |
| | $GWAS_{null}$, Y2 | | 0.0096 | 0.0520 | -0.0328 | 0.6560 |
| Sex + YOB + TDI + Eats pork + Eats beef + Birth location + Excludes dairy + Health + Number of births | $GWAS_{env}$, $GWAS_{null}$ | All | -0.0075 | -0.0015 | -0.0136 | 0.0152 |
| | $GWAS_{env}$, Y1 | | -0.0191 | 0.0152 | -0.0534 | 0.2744 |
| | $GWAS_{env}$, Y2 | | -0.0193 | 0.0154 | -0.0539 | 0.2759 |
| | $GWAS_{null}$, Y1 | | -0.0116 | 0.0240 | -0.0471 | 0.5226 |
| | $GWAS_{null}$, Y2 | | -0.0117 | 0.0242 | -0.0477 | 0.5223 |
| YOB + TDI + Eats pork + Eats beef + Birth location + Excludes dairy + Health + Number of births | $GWAS_{env}$, $GWAS_{null}$ | F | -0.0113 | -0.0019 | -0.0206 | 0.0180 |
| | $GWAS_{env}$, Y1 | | -0.0491 | 0.0080 | -0.1061 | 0.0921 |
| | $GWAS_{env}$, Y2 | | -0.0544 | 0.0041 | -0.1128 | 0.0681 |
| | $GWAS_{null}$, Y1 | | -0.0378 | 0.0215 | -0.0970 | 0.2114 |
| | $GWAS_{null}$, Y2 | | -0.0431 | 0.0175 | -0.1036 | 0.1632 |
| YOB + TDI + Eats pork + Eats beef + Birth location + Excludes dairy + Health | $GWAS_{env}$, $GWAS_{null}$ | M | -0.0046 | 0.0033 | -0.0124 | 0.2537 |
| | $GWAS_{env}$, Y1 | | -0.0020 | 0.0392 | -0.0433 | 0.9227 |
| | $GWAS_{env}$, Y2 | | 0.0001 | 0.0412 | -0.0410 | 0.9966 |
| | $GWAS_{null}$, Y1 | | 0.0025 | 0.0451 | -0.0400 | 0.9071 |
| | $GWAS_{null}$, Y2 | | 0.0047 | 0.0472 | -0.0379 | 0.8298 |

*Mean, upper, and lower differences reflect the mean difference and upper/lower bounds of the 95% CI for the difference between the two PGS $R^2$ compared, per model. The mean difference is relative to the first PGS listed in the 'PGS models compared' column.

**Table S2.15.** Correlation between PGS scores and environmental covariates. No models meet significance at $p < 0.01$.

| PGS | Model (R syntax) | Sample | $R^2$ | Adj. $R^2$ |
|---|---|---|---|---|
| **$PGS_{Y1}$ (*n* = 898,486)** | Score ~ Sex + TDI + Birth location + Eats pork + Eats beef + Excludes dairy + Health + Number of births | Full test sample with unimputed environmental covariate data (*n* = 907) | 0.011 | 0.001 |
| | | F (*n* = 427) | 0.023 | 0.004 |
| | | M (*n* = 480) | 0.01 | -0.005 |
| **$PGS_{Y2}$ (*n* = 1,032,672)** | Score ~ Sex + TDI + Birth location + Eats pork + Eats beef + Excludes dairy + Health + Number of births | Full test sample with unimputed environmental covariate data (*n* = 907) | 0.013 | 0.003 |
| | | F (*n* = 427) | 0.022 | 0.004 |
| | | M (*n* = 480) | 0.011 | -0.004 |
| **$GWAS_{env}$ PGS (*n* = 896,160)** | Score ~ Sex + TDI + YOB + Birth location + Eats pork + Eats beef + Excludes dairy + Health + Number of births | Full test sample with unimputed environmental covariate data (*n* = 907) | 0.008 | -0.002 |
| | Score ~ TDI + YOB + Birth location + Eats pork + Eats beef + Excludes dairy + Health + Number of births | F (*n* = 427) | 0.017 | -0.002 |
| | Score ~ TDI + YOB + Birth location + Eats pork + Eats beef + Excludes dairy + Health | M (*n* = 480) | 0.014 | -0.001 |
| **$GWAS_{null}$ PGS (*n* = 896,160)** | Score ~ Sex + TDI + YOB + Birth location + Eats pork + Eats beef + Excludes dairy + Health + Number of births | Full test sample with unimputed environmental covariate data (*n* = 907) | 0.011 | 0.001 |
| | Score ~ TDI + YOB + Birth location + Eats pork + Eats beef + Excludes dairy + Health + Number of births | F (*n* = 427) | 0.018 | -0.001 |
| | Score ~ TDI + YOB + Birth location + Eats pork + Eats beef + Excludes dairy + Health | M (*n* = 480) | 0.018 | 0.004 |

**Environmental adjustments and population stratification**

Following the guidance of Zaidi and Mathieson [11] for evaluating evidence of residual population stratification in GWAS-PGS results, we implemented an additional assessment to: A) ascertain if environmental adjustments in GWAS modeling mitigate the downstream effects of population stratification on PGS development; and B) identify which of our PGS equations were likely to be least influenced by population stratification. Using the PGS development sample, we regressed $PGS_{Y1}$, $PGS_{Y2}$, $GWAS_{null}$ PGS, and $GWAS_{env}$ PGS scores on participant region of origin and compared the resulting summary statistics for significant associations between region and score. Regions of origin with fewer than five participants were removed prior to analyses to reduce the likelihood of identifying significant associations resulting from individual outliers (*n* = 832). Results indicate that all PGS scores reflect a small, but significant, effect of population stratification tracked by participant region of origin (Table S2.15). $GWAS_{env}$ PGS score reflects the least evidence of residual population stratification, with participant origin explaining ~2% less variance than for $PGS_Y$ scores (Table S2.16). At $p < 0.05$, no significant associations between the $GWAS_{env}$ PGS score and participant region of origin were identified, however significant associations were identified for all other PGS scores (Figure S2.10). The lack of significant associations derived from the $GWAS_{env}$ PGS in comparison to $GWAS_{null}$ and $PGS_Y$ scores suggests that including appropriate environmental covariates can help reduce the effects of population stratification on PGS development that are not explained by genetic PCs and GRM.

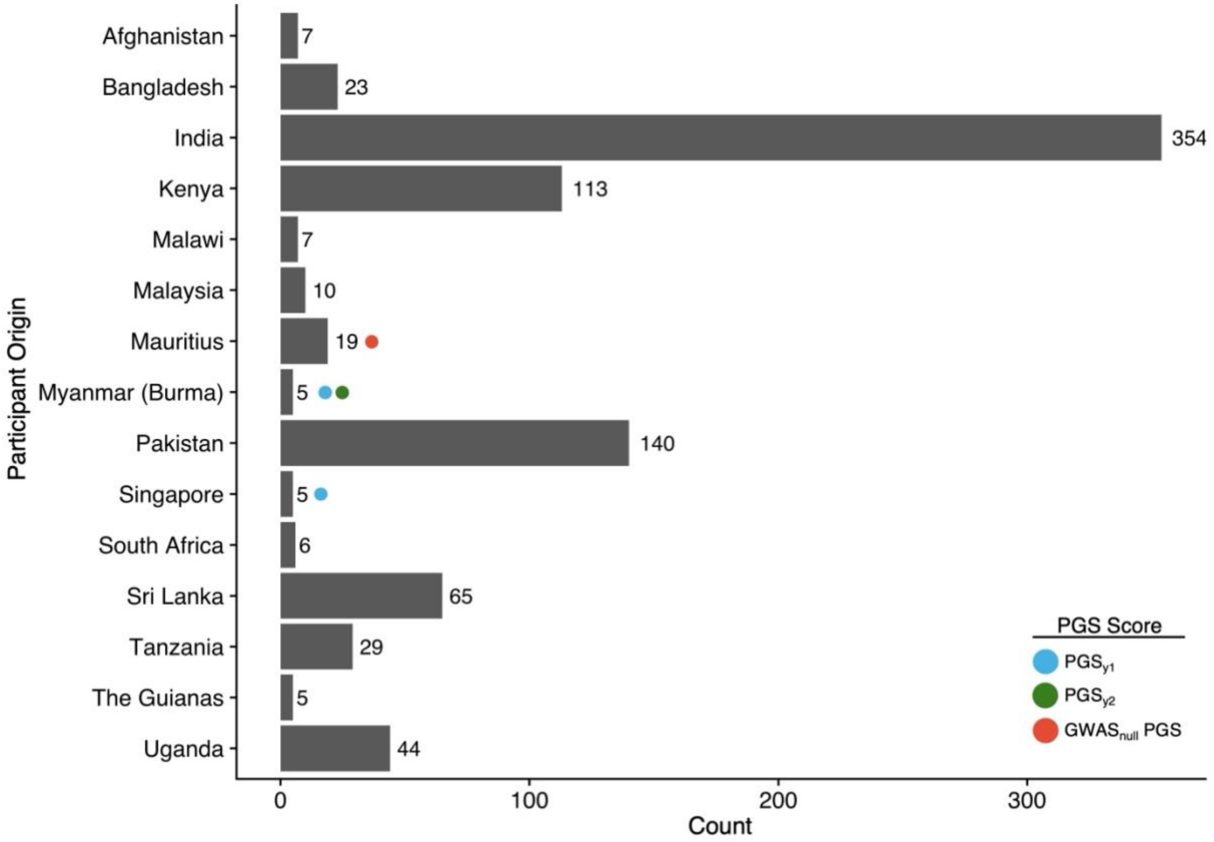

**Figure S2.10.** Sample counts of participant region of origin used for assessment of population stratification on PGS scores ($PGS_{Y1}$ [blue], $PGS_{Y2}$ [green], $GWAS_{null}$ PGS [red], and $GWAS_{env}$ PGS [not visualized]. Participant origins significant at $p < 0.05$ are marked with a circle. No significant associations were found between the $GWAS_{env}$ PGS score and participant origin.

**Table S2.16.** Summary statistics of OLS model regressing PGS score on participant region of origin (as calculated using base functions within the statistical package R).

| PGS | *p* | $R^2$ | Adj. $R^2$ |
|---|---|---|---|
| **$PGS_{Y1}$** | < 0.001 | 0.046 | 0.030 |
| **$PGS_{Y2}$** | < 0.001 | 0.048 | 0.032 |
| **$GWAS_{null}$ PGS** | 0.01 | 0.034 | 0.018 |
| **$GWAS_{env}$ PGS** | 0.03 | 0.031 | 0.014 |

### 2.5: Extended discussion

#### Heritability of environmental covariates

The heritability of each environmental covariate used in the $GWAS_{env}$ model was investigated to ascertain the potential magnitude of shared genetic causality with height. As of the submission date of this manuscript, the only published SNP-based heritability ($h^2$) estimates available for our variables of interest were those presented in the Neale Lab SNP-Heritability Browser [http://www.nealelab.is/uk-biobank/], which are derived solely from European UKB participants. The dietary variables published by the Neale Lab have low levels of $h^2$ (~0.03 - 0.04 for pork, beef, and dairy variables). Townsend deprivation index (TDI) is comparably low (~0.03) as is "Number of live births" (~0.06). Overall health, however, has a larger $h^2$ estimate of ~0.1, suggesting that inclusion of this variable as a covariate may result in biased genetic effect estimates. As these $h^2$ estimates are not directly comparable to the UKB South Asian sample, we ran additional GWAS for overall health (following the same procedures as our height GWAS) and calculated observed scale $h^2$ from these summary statistics using LDSC [18]. 1000 Genomes South Asian data were used to create the LD scores and regression weights required to use LDSC (after merging, 4,022,277 SNPs were used in this analysis). Our results suggest a lower $h^2$ estimate than derived from the UKB European sample, however, SEs are large and 95% CI's extend past 0 (Figure S2.11 and Table S2.17). This suggests that "overall health rating" is a (predictably) noisy variable likely capturing many proxies for actual health, and our data are underpowered for a reliable $h^2$ estimate. The low $h^2$ estimates suggest that any shared causal loci between height and health likely do not represent a large, direct genetic effect on height.

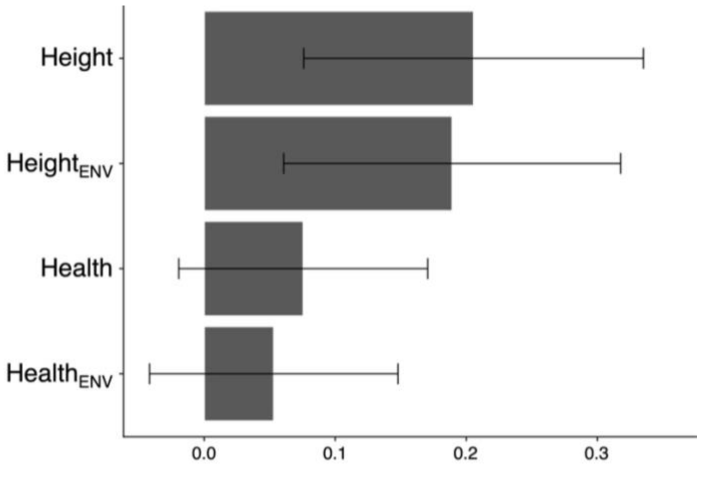

**Figure S2.11.** LDSC observed scale heritability estimates for height and health, derived from GWAS using either the $GWAS_{null}$ covariates (designated above without subscripts) or $GWAS_{env}$ covariates (designated by the "ENV" subscript). Health was not used as a covariate in the environmentally-adjusted GWAS when it was the outcome phenotype. Error bars reflect 95% CI intervals.

**Table S2.17.** LDSC observed scale heritability estimates for height and health, derived from GWAS using either the $GWAS_{null}$ covariates ("Null") or $GWAS_{env}$ covariates ("Environmental").

| Trait | GWAS Model | $h^2$ | SE |
|---|---|---|---|
| Height | Environmental | 0.19 | 0.064 |
| | Null | 0.21 | 0.065 |
| Health | Environmental | 0.05 | 0.047 |
| | Null | 0.08 | 0.048 |

### GWAS and PGS comparisons

Sex biases

PGS performance has higher $R^2$ for female height than for male height. However, this bias is reduced when adjusting for environmental contributors (Table S2.13). Though not explicitly tested here, these results may be a consequence of overall higher variance in male height than female height [12]; such variance may reflect greater male plasticity [13]. Del Giudice et al. [13] hypothesize that androgen exposure contributes to phenotypic plasticity, ergo, for traits that differ in their variability between males and females, males are more likely to have higher variance in phenotypic expression. Additionally, androgens have repeatedly been implicated in skeletal homeostasis and development [14, 15, 16], lending credence to their role in male plasticity for height. Enrichment of *C20orf173* within the $GWAS_{env}$ FUMA-GWAS functional annotation results (Table S2.9; Fig. S2.8) further supports this relationship as *C20orf173* is over-expressed in the testis (Human Protein Atlas [17]). *C20orf173* has not been directly associated with height prior to our analysis (per GWAS Catalog results, September 2024). Therefore, if male height is more plastic than female height, differential environmental exposures are likely to have a stronger effect on males during ontogeny. By including environmental covariates in our GWAS, we are able to improve male height PGS performance (and reduce the discrepancy in $R^2$ between sex-stratified samples).

GWAS and PGS variant sets

Following common practice, and recommendations for PGS development with LDpred2, we intersected our (GAsP imputed) GWAS summary statistics with the HapMap 3 plus (HM3+) variants. This reduces the total possible number of PGS variants to ~1.4 million SNPs in HM3+, of which 1,069,741 overlap with our GWAS SNPs ($n$ = 6,946,575). By removing the majority of our GWAS SNPs prior to PGS development, we may have limited the full variance in height possible to be explained by the PGS SNPs. As a post-hoc assessment, we investigated the relationship between the GWAS summary statistics and HM3+ variants. When assessing the intersection of GWAS and HM3+ SNPs by alpha thresholds (as in Table 4), 0 SNPs meeting genome-wide significance can be found in HM3+ (Table S2.18). This alone, is not cause for concern given the likelihood that many GWAS SNPs may be "tagged" by (i.e., in high LD with) the SNPs in HM3+. The effects associated with the top GWAS SNPs that are found in both variant sets do appear to be attenuated (Figure S2.12), suggesting that if the top SNPs are not tagged by those in HM3+, the full range of genetic effects may not be represented in the PGS models. To visualize the genome-wide representativeness of GWAS SNPs by HM3+, we created overlaid Manhattan plots, with SNPs annotated with the nearest gene, using the *topr* package in R [19]. Qualitatively, most "peaks" (loci with smallest p-values) are, in fact, tagged by HM3+ (Figure S2.13). However, some regions with sub-genome-wide significance are fully omitted in the HM3+ array (e.g. see chromosome 7, gene POR in Fig. S2.13), again hinting at the possibility that this SNP intersection may not allow for the ideal set of genetic effects to be included in PGS model. Future GWAS-PGS development may benefit from exploring these limitations in more depth, and directly comparing the outcomes between PGS equations derived from arrays meant to provide broad, globally diverse variants and those curated to highlight the genetic variance for a specific meta-population.

**Table S2.18**. Number of SNPs meeting different significance thresholds in GWAS summary statistics that are also found in the HM3+ array.

| *p* < | $10^{-5}$ | $10^{-6}$ | $10^{-7}$ | $5^{-8}$ |
|---|---|---|---|---|
| $GWAS_{null}$ | 45 | 25 | 0 | 0 |
| $GWAS_{env}$ | 50 | 24 | 0 | 0 |

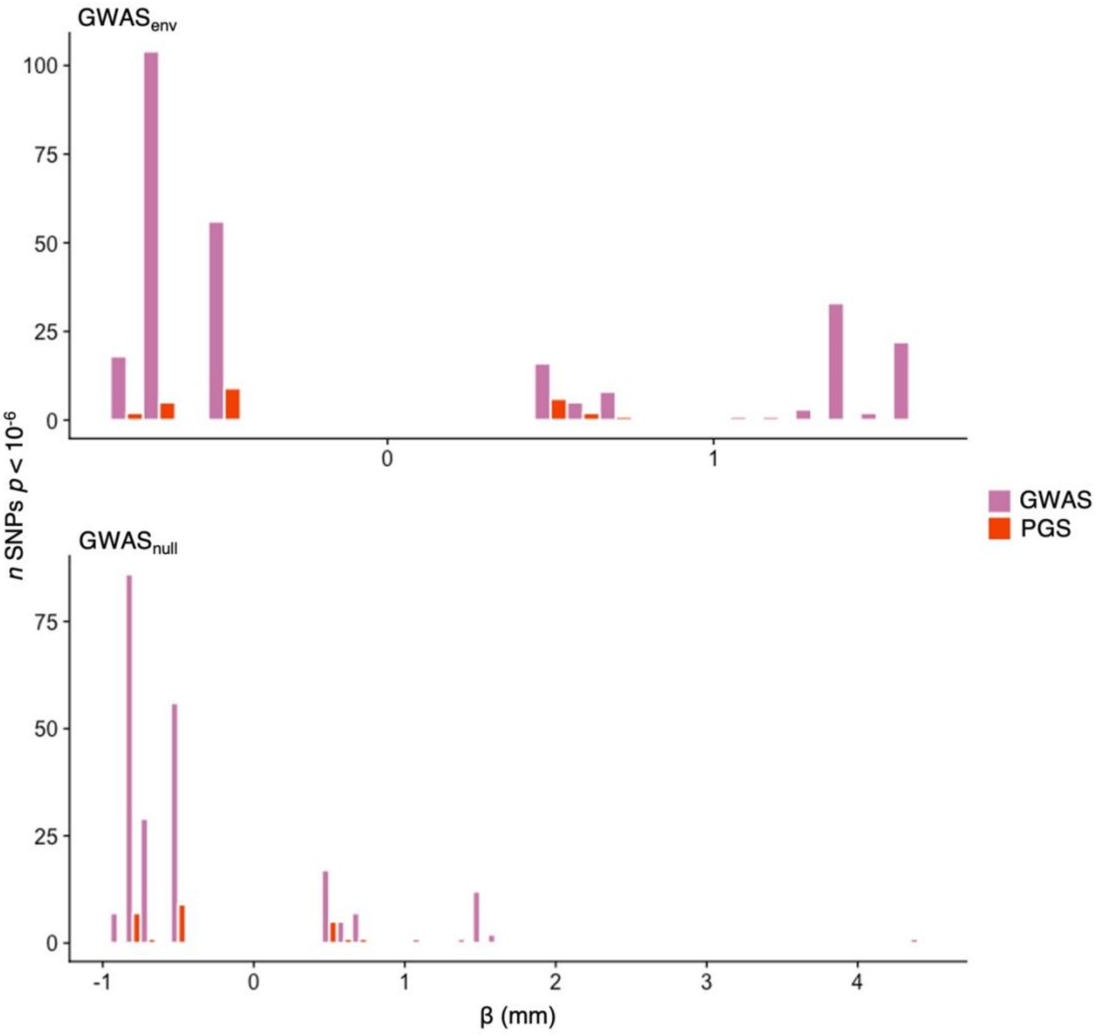

**Figure S2.12.** Distribution of effect estimates for $GWAS_{env}$ SNPs (top) and $GWAS_{null}$ SNPs (bottom) with significance $p < 10^{-6}$. GWAS variants meeting the alpha threshold that are not found in the associated HM3+ PGS model are pink. GWAS variants that meet the alpha threshold and are included in the HM3+ PGS model are red.

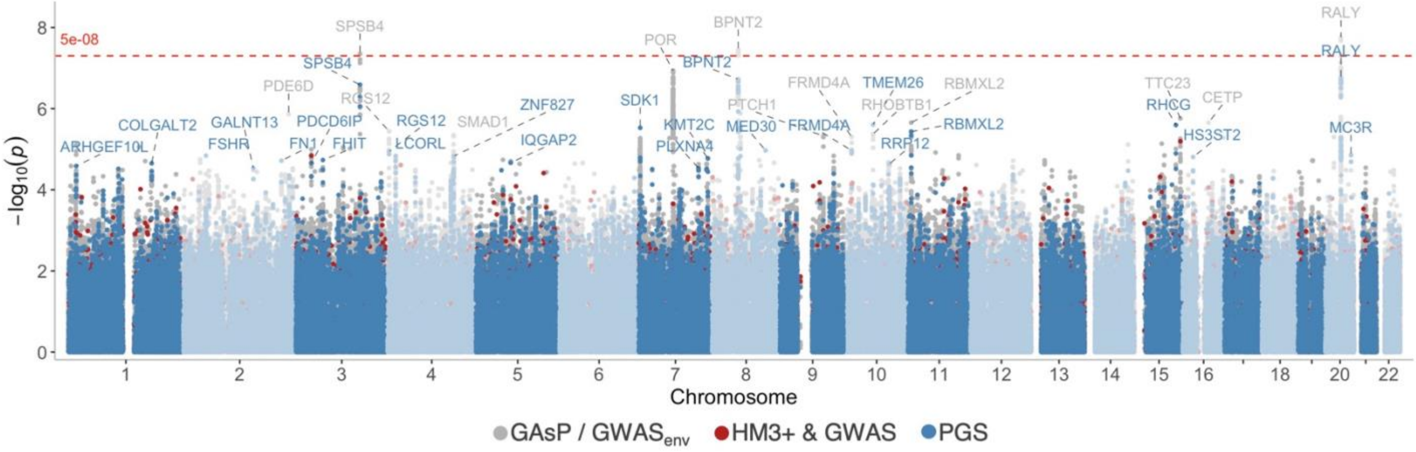

**Figure S2.13.** Manhattan plot of $GWAS_{env}$ SNPs colored by array inclusion. All variants plotted are found in the GAsP panel and our GWAS. Variants in grey are not found in HM3+ or the downstream PGS model. Variants in red are found in both HM3+ and GAsP but were not included in the PGS. Variants in blue are found in both HM3+ and GAsP and are included in the PGS. SNPs are annotated with the nearest gene [19]. Genome-wide significance is denoted by the red dashed line.
